# Visualizing the Rapid Degradation and Differential Binding of Membrane-Bound Human ACE2 Protein Upon Binding and Internalization of the SARS-CoV-2 Spike RBD Protein

**DOI:** 10.1101/2024.03.07.583884

**Authors:** Paul Feinstein

## Abstract

The SARS-CoV-2 betacoronavirus infects humans through binding the Angiotensin Converting Enzyme 2 (ACE2) protein that lines the nasal cavity and lungs, followed by import into a cell utilizing the Transmembrane Protease, Serine 2 (TMPRSS2) cofactor. ACE2 binding is mediated by an approximately 200-residue portion of the SARS-CoV-2 extracellular spike protein, the receptor binding domain (RBD). Robust interactions are shown using a novel cell-based assay between an RBD membrane tethered-GFP fusion protein and a membrane-bound ACE2-Cherry fusion protein. Several observations were not predicted, including rapid and sustained interactions leading to internalization of the RBD fusion protein into ACE2-expressing cells and rapid downregulation of ACE2-Cherry fluorescence, suggesting that a membrane-associated form of RBD found on the viral coat may have long-term system-wide consequences on ACE2-expressing cells. Targeted mutation in the RBD disulfide Loop 4 led to a loss of internalization for several variants tested. However, a secreted RBD did not cause ACE2 downregulation of ACE2-Cherry fluorescence. Omicron BA.1 and BA.2 variants have altered their dependency on the amino terminus (Nt) of the ACE2 protein. In contrast, the H-CoV-NL63 RBD is only dependent on the ACE2 internal region for binding, leading to the conclusion that the RBD binding surface of ACE2 appears relatively fluid and amenable to internalizing a range of novel variants.

## Introduction

The COVID-19 worldwide pandemic continues to produce widespread illness and death. The causative agent of this disease is the Severe Acute Respiratory Syndrome coronavirus 2 (SARS-CoV-2 aka SARS2), which is similar the SARS-CoV (SARS1) virus that had a limited outbreak between 2002-2003 (1). Both SARS1 and SARS2 belong to the betacoronavirus subfamily. The spread of SARS1 was cut short by lockdowns, but followed by a second betacoronavirus, Middle East respiratory syndrome (MERS-CoV) in 2012, which continues to infect individuals (2). Vaccines against SARS2 are reducing the death caused by the proliferation of viral particles (3-5). But vaccinations tend not to halt the transmission of viral particles, which occurs in just days after infection, prior to the rise in immune response (6). In addition, the mutation rate of SARS2 is unyielding, leading to the prospect of infections by novel variants for the foreseeable future. The medical industry must find viral transmission mitigation strategies (7-9). While masks form a reasonable barrier to SARS2 infection, societies have not embraced this as a long-term solution to mitigate infections. Both vaccine protocols and drugs such as Paxlovid (10), have significantly reduced morbidities once an infection has occurred.

How are SARS1 and SARS2 infections initiated? It has been shown that the trimeric Spike coat protein for both viruses latch onto a cell via binding to a dimer of the human Angiotensin-Converting Enzyme 2 protein, ACE2, that resides in the plasma membrane. ACE2 provides 740 amino-terminal (Nt) residues in the extracellular space, whilst only a short 40-residue carboxy terminus (Ct) extends cytosolically (11-13). Unlike SARS1, SARS2 has been pre-cleaved to generate S1 and S2 subunits that remain associated by non-covalent interactions. Trimeric S1 proteins loosely associated with an uncleaved S2 subunit cannot be mimicked without using a pseudovirus. Once the SARS2 Spike S1 subunit binds an ACE2 dimer at the plasma membrane, the Transmembrane Protease, serine 2, TMPRSS2, catalyzes the endocytosis of the virus into the cell by cleaving the second Spike S2 subunit, revealing the S2’fusion peptide (14-16). Mitigating either binding of ACE2, functionality of TMPRSS2 (7, 8) or endocytosis could prove an effective means of suppressing disease onset. Small molecules, although not volatile, that target TMPRSS2 functionality have been identified that may limit viral load (7)

ACE2 is well expressed in the cilia or microvilli throughout epithelial cells from the nose to the lungs. In the olfactory epithelium, ACE2 is not expressed by olfactory sensory neurons (OSN) but robustly expressed on the intertwined supporting sustentacular cells (SUS) as well as Bowman’s glands (17, 18). The nasal infection leads to a down-regulation of OSN-produced odorant receptor (OR) signal transduction components (19). It is not completely understood how SUS cell infection by SARS2 contributes to anosmia (6). In our pursuit of studying odorant receptor (OR) protein trafficking to the plasma membrane, we have fused ORs and other G-protein Coupled Receptors to Green Fluorescent Protein (GFP) and expressed them in an olfactory placodal cell line (OP6) with robust filopodia (20-22). Here, I have repurposed the trafficking assay to study the molecular details of how the SARS2 Spike S1 and ACE2 proteins interact.

Cellular entry for SARS2 is initiated by the cleaved Spike S protein, the 769 residue S1 subunit. Detailed analysis of the SARS1-ACE2 binding (the 193 amino acid region- the receptor binding domain, RBD) led to the identification of the SARS2-S1 RBD between residues 333-529 (23). This 197-residue region recognizes three domains of ACE2 within residues 19-357. By contrast, angiotensin II binds a different portion of ACE2 (24). The sufficiency of the RBD was shown in binding studies using a soluble form of ACE2, sACE2, in EC50 measurements from neutralization with RBD in the micromolar range and to a dimeric ACE2 protein in the nanomolar range (25). The ACE2 protein traffics to the plasma membrane with a signal peptide that is cleaved, resulting in the mature protein, residues 19-805. The RBD region binds to sixteen ACE2 residues distributed within its amino terminal (Nt) 19-83 residues, as well as another five residues within an internal region containing residues 330, 353-357. These interactions and co-crystallographic studies of SARS1 and SARS2 lay the foundation of this current work (1, 26-28).

Here, the RBD domain and full-length ACE2 proteins are linked to different fluorescent proteins and determined if they could interact in OP6 cells. For the Type I transmembrane ACE2 protein an intracellular Cherry fusion is generated at its Carboxy terminus (Ct), whereas the 197-residue SARS2 RBD was fused to a secretion peptide at its Nt and either a single transmembrane (TM) domain protein or the 7-TM domain β2AR at its Ct along with an intracellular located GFP fusion.

This in vitro assay allowed us to observe fast and viral-free SARS2-RBD protein binding to ACE2 in cells, which mimic some of the binding and internalization properties of the pseudotype virus infection of ACE2-expressing cells: 1) rapid (a few minutes) and sustained binding between SARS2-RBD protein and ACE2; 2) a Spike S2 subunit independent uptake of a membrane-bound RBD from one cell by other cells expressing ACE2; and 3) a downregulation of ACE2-Cherry protein upon binding to RBD suggesting an unsuspected effect on ACE2 accessibility for binding of additional viral particles. This novel in vitro assay provides the basis to identify the critical amino acid sequences involved in binding of SARS2-RBD to ACE2 and its internalization. By contrast, pseudotype viruses provide data on viral replication as a surrogate for RBD binding and viral membrane internalization by ACE2-expressing cells.

A multitude of generated mutations in conserved residues, despite being predicted to have important roles in binding from crystal structures, do not abrogate SARS2-RBD binding to ACE2 revealing the distributed nature of interactions. One critical residue that did not affect plasma membrane trafficking of the protein, Y489A, nearly abolished the RBD-ACE2 interactions. This residue juxtaposes C488 that is critical for disulfide Loop 4 formation (C480-C488). A debilitating C480A, C488A SARS2-RBD double mutant was produced, which suggests disruption of disulfide bridges would abolish binding to ACE2. RBD-ACE2 binding deficits were recapitulated in our assay using concentrations of disulfide-reducing agent dithiothreitol (DTT) analogous to what has been demonstrated with pseudotype viruses (29).

The alphacoronavirus H-CoV-NL63 (NL63) also binds ACE2 through a 136-residue RBD domain with little homology to SARS2-RBD (7% amino acid identity). Thus, it seems likely that the biological properties of ACE2 make it an attractive target for cellular entry of viruses. NL63-RBD contains three potential disulfide bonds with one cysteine residue common to SARS2 disulfide Loop 4 and contains three other homologous residues. Again, DTT blocked NL63-RBD binding to ACE2 as well. It is plausible that these or other volatile disulfide-reducing agents could mitigate viral infections despite their pleiotropic effects. The ability for RBD domains alone to affect ACE2 protein suggests that vaccines containing “binding competent” RBD domains may alter in vivo ACE2 protein levels and lead to adverse reactions. The data suggest that vaccines with mutation(s) in RBD, which cannot bind to ACE2 may lead to fewer side effects. Some of the Long COVID symptoms may be a result of sustained ACE2 down-regulation (30).

To further pursue how RBD binds to ACE2, a structure-function mutagenesis on ACE2 was undertaken in hopes of identifying an antagonist of RBD binding. Mutations around disulfide Loop 4 within SARS2-RBD that reduced or abolished binding to ACE2, suggested ACE2 Nt proximal regions that might be critical for binding. Here, I show mutagenesis of the ACE2 Nt region identifies a swap of eight residues from the Least Horseshoe Bat (*Rhinolophus pusillus*) into the Nt proximal portion of the human ACE2 (residues between 20-34), which can abolish SARS2-RBD binding. But both Omicron BA.1 and BA.2-RBDs were still able to interact with this ACE2 mutant. An addition of three mutations (L79A, M82A, Y83A: LMY➔A) in the Nt distal region of this chimeric ACE2 can completely abolish Omicron BA.1 or BA.2-RBD internalization, while the control NL63-RBD from alphacoronavirus is not affected by any Nt proximal or distal mutations.

However, both NL63-RBD and SARS2-RBD were both unable to bind ACE2 when its internal residues K353 and D355 were both mutated. Using co-crystallization data as a guide, a double mutant T500A, G502A on SARS2-RBD is also able to abolish binding to ACE2. A set of peptides based on K353-G354-D355 of ACE2 were not able to inhibit either SARS2-RBD nor NL63-RBD binding. The totality of the data suggests: 1) multiple weak binding sites on ACE2 sum to create high affinity; 2) the internal region residues may be critical for all RBD binding sites necessary for ACE2 internalization; and 3) there are multiple permutations for RBD binding to ACE2. The changing nature of RBD variants preferentially binding to different ACE2 residues may correlate with the severity of symptoms by altering the degree of internalization by the ACE2 protein.

## Results

### RBD Mutants

#### The RBD➔ACE2 cell-based binding assay

The SARS-CoV-2 (SARS2) betacoronavirus infects cells via its furin-pre-cleaved Spike protein, generating S1 and S2 subunits that remain non-covalently associated. Internalization of the SARS2 virus is achieved by its membrane localization mediated by the binding of the Spike S1 subunit to its cellular receptor, the Angiotensin Converting Enzyme 2 (ACE2), followed by the Transmembrane Protease, Serine 2 (TMPRSS2) cleavage of the Spike S2 subunit, revealing the S2’ fusion protein. There have been many strategies to characterize the molecular interactions between the Spike S1 subunit and the ACE2 protein. Here, I set up an in vitro cell interaction assay based on the experiments by Chan et al. (25), with a subdomain of the S1 subunit-the receptor binding domain (RBD) and ACE2, both fused to different fluorescent proteins and expressed in heterologous cells **(Figure 1A)**(25).

**Figure 1:**
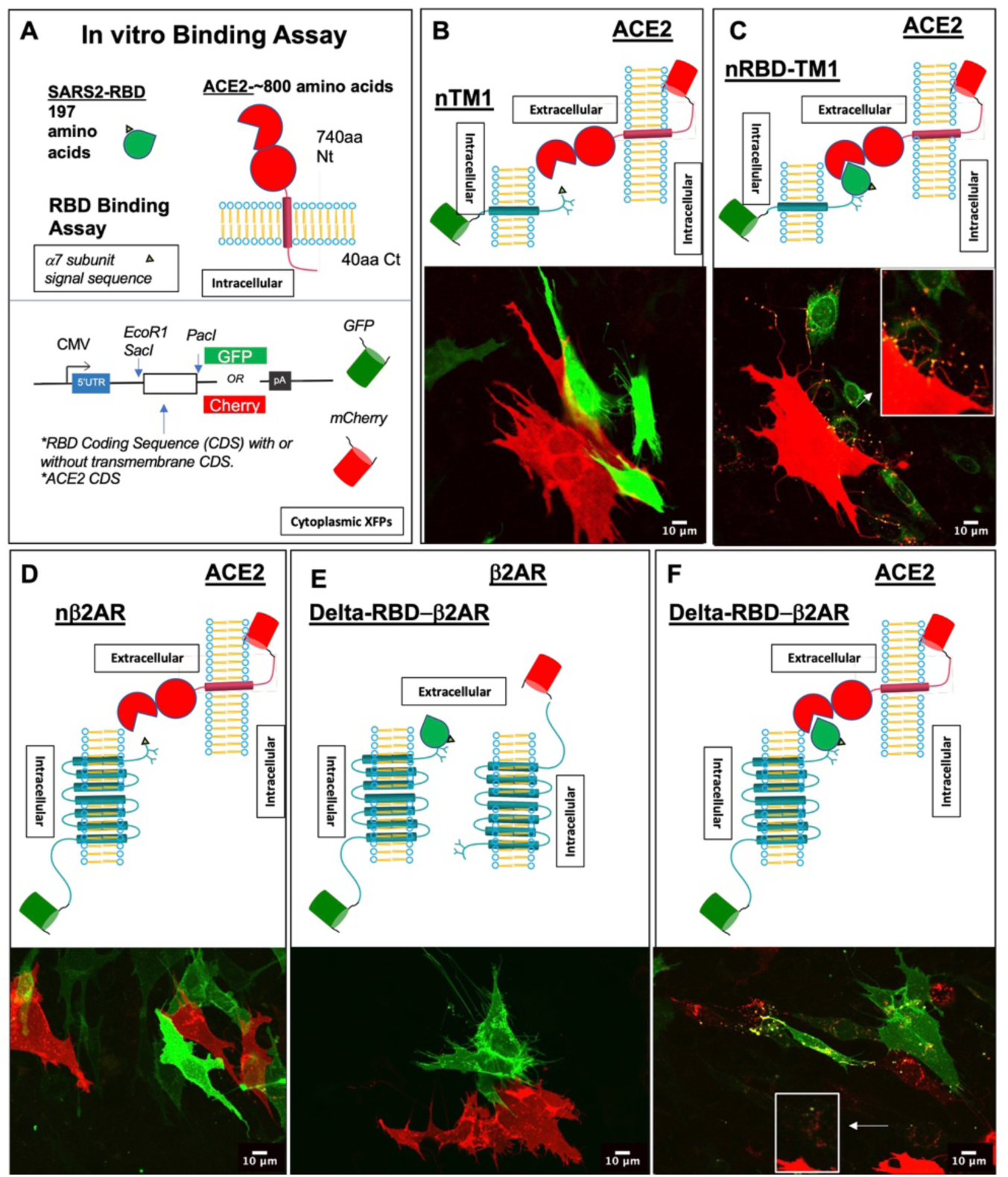
An in vitro substitute for SARS2 virus binding to ACE2-expressing cells. **A.** Experimental system to identify a compound that mitigates SARS2-RBD and ACE2 binding in cell lines. Expression vectors with Cherry and GFP contain EcoR1-PacI restriction sites, whereby cloning of SARS2-RBD variants or ACE2 mutants will generate in frame fusion proteins. **B.-F.** Confocal images of separate electroporations into OP6 cells and co-plated overnight (O/N). All RBD constructs contain the nicotinic receptor α7 subunit (n) signal sequence at their N-termini **B.** Expression and co-plating of the first 60amino acids of β2AR (nTM1) to GFP and human ACE2 fused to Cherry. No double fluorescent cells observed: red (Cherry) and green (GFP). **C.** SARS2-RBD-β2AR-TM1 (nRBD-TM1) is fused to GFP and co-plated with ACE2-Cherry expressing cells. Red filopodia (20) are seen with green fluorescent on its distal points (see inset, magnification). **D.** Control co-plating of nβ2AR-GFP and ACE2-Cherry expressing cells show no double fluorescence. **E.** Control co-plating of the Delta variant, Delta-RBD-β2AR-GFP and β2AR-Cherry expressing show no double fluorescence. **F.** Co-plating Delta-RBD-β2AR-GFP and ACE2-Cherry expressing cells show dramatic internalization of GFP fluorescently tagged protein and no obvious morphology to the ACE2-Cherry expressing cells. Inset shows filopodia of ACE2-Cherry cells with GFP. Scale bars at 10μm.

G-Protein Coupled Receptor (GPCR) trafficking to the plasma membrane (PM) has been studied by my lab for the last decade, where expression vectors with GPCRs fused to fluorescent proteins are transfected in an olfactory neuronal cell line (OP6) that is rich in filopodia extensions (20-22). I previously showed that the presence of fluorescence in the filopodia accurately correlated with PM expression (20-22). OP6 cells are grown at 33°C, which is coincidentally similar to the temperature at which the nasal epithelium would be infected by SARS2 in ambient air (31). I co-opted our in vitro expression system to study RBD and ACE2 interactions. Unless otherwise stated RBD refers to the parental SARS2 Wuhan isolate. Analysis was initiated with a Cherry fusion to the carboxy terminus (Ct) of human ACE2 and a super-folding variant of GFP (sfGFP) fused to the Ct of the secreted RBD. Expression of ACE2-Cherry is readily observed in the filopodia **(Figure 1B-D and F).** In contrast, the secreted-SARS2(sRBD)-sfGFP (s = hemagglutinin-cleavable signal) was detected only within the cell (not shown). When separate transfections were co-plated and cultured overnight, on occasion an ACE2-Cherry expressing cell could be seen with green particles, suggesting uptake of Secreted-SARS2-sfGFP protein (not shown); The membrane of the ACE2-Cherry cells was not labeled with sfGFP. These initial studies show that secretion through the membrane of sRBD-sfGFP may not be occurring.

In previous studies of GPCR tagging, the nicotinic receptor α7 subunit (nrα7) signal peptide (n) was shown to be effective in targeting a functional GFP added to the extracellular portion of a GPCR (nGFP-GPCR) (32, 33). Thus, I tested the nrα7 signal sequence along with RBD to create Secreted-SARS2-nRBD-GFP and Secreted-Delta-nRBD-GFP (delta SARS2 variant). Because the delta variant was becoming dominant at the time of these experiments, it was chosen for comparative purposes to the RBD (Wuhan) form. Again, no membrane GFP fluorescence could be observed on ACE2-Cherry expressing cells when co-plated after transfection. But it appeared that there may be some granules of GFP fluorescence within ACE2-Cherry cells after 12-18hr incubation. The interpretation of these results were either the sRBD-sfGFP or nRBD-GFP proteins were not secreted or if they were secreted, then perhaps they were rapidly internalized and degraded by the ACE2-Cherry expressing cells. Overall, these results suggested a reconsideration of this first approach.

The ACE2-Cherry fusion was successfully expressed at the PM but was unsuccessful at binding sRBD or nRBD-GFP particles. To prevent secretion of RBD protein, the nrα7-SARS2-RBD (nRBD) was tethered to the extracellular portion of the single transmembrane domain (TM1) of the mouse β2AR (the first 60 residues) that is also Ct fused to GFP. This generated nRBD-TM1-GFP and a control vector nTM1-GFP. Both fusions were expressed in cells and co-cultured separately with ACE2-Cherry expressing cells. As expected, nTM1-GFP trafficked to the PM (34) but did not interact or transfer any fluorescence to the ACE2-expressing cells **(Figure 1B).** In contrast, the nRBD-TM1-GFP fluorescence was found on the tips of the filopodia expressing ACE2-Cherry **(Figure 1C and inset)**; It should be noted that ACE2-Cherry cells had well-demarcated Cherry fluorescence in the filopodia, and GFP-expressing cells were not necessarily nearby. This result suggested that nRBD-TM1-GFP protein may have been secreted into the extracellular space. OP6 Cells do not express TMPRSS2 based on mass spectroscopy (not shown), and no TMPRSS2 target sequences (Arg-Serine dimers) are found within this nRBD-TM1-GFP construct; Had there been any protein digestion within GFP, then no fluorescence could have been observed.

To ensure that the RBD domain is within the PM, nRBD was tethered to the full-length, 7TM β2AR; These 7TM proteins cannot be secreted due to their seven hydrophobic domains. Transfection, co-plating and culturing of cells with the control nβ2AR-GFP and the control ACE2-Cherry (**Figure 1D**) or control β2AR-Cherry (**Figure 1E**), revealed fusion proteins on the PM of the cells based on filopodial localization of the fluorescence; No transfer of fluorescence was observed between cells **(Figure 1D, 1E)** (20-22). When nRBD-β2AR-GFP (henceforth SARS2-RBD) expressing cells were co-cultured overnight with ACE2-Cherry expressing cells, robust cell-cell contacts and filopodial binding were observed **(Figure 1F and Inset)**. Moreover, ACE2-Cherry fluorescence was no longer well distributed in the cell, and green-fluorescent particles could be observed within ACE2-Cherry cells as particulates. The Green fluorescent particles taken up by ACE2-Cherry expressing were indeed properly folded β2AR as they were able to bind a β2AR fluorescent antagonist (**Figure S1**, CellAura fluorescent β2antagonist [(±)-propranolol]). I confirmed that Delta-nRBD-β2AR-GFP (*henceforth* Delta-RBD) cells expressing the variant Delta (L452R, T478K)-RBD qualitatively interacted with ACE2-Cherry cells as did the SARS2-RBD form.

Our expertise in olfaction led us to consider that there may be an odor, which can interfere with SARS2-RBD binding to ACE2. With our protein interaction assay in hand, 36 concentrated odor mixtures from the kit “Le Nez Du Café (LNDV)”(35) were tested for their ability to antagonize RBD binding to ACE2. Only one of the mixtures at a 1:500 dilution, *medicinal*, appeared to qualitatively reduce, but not eliminate binding; The degree of interaction could not be quantified by microscopy. Had the odor completely blocked binding, this would have been readily quantifiable. Thus, a quantification procedure to measure the extent of antagonism needed to be set up.

#### The RBD➔ACE2 flow cytometry based binding assay

How strong is the RBD-ACE2 cell-based binding of green fluorescence in or on red fluorescent cells? Would this be observed after trypsinization of co-cultures followed by single cell flow cytometry or FACS analysis? The binding affinity of RBD to ACE2 has been previously calculated at nanomolar equilibrium rates (Kd) (1, 25). I tested the stability of interaction with overnight co-cultures for various GFP fusions against ACE2-Cherry cells, followed by trypsinization, filtering and flow cytometry counts; DAPI positive single cells were quantified for GFP fluorescent positive cells, Cherry fluorescent positive cells, and Cherry/GFP double fluorescent positive cells that were above background. Flow cytometry counts for ACE2-Cherry expressing cells were absent in GFP expression at the gate set for Cherry expression, GFP fusion protein expressing cells were absent in Cherry expression at the gate set for GFP expression, and the double positive window was set above both GFP and Cherry gates. GFP fusions used were: nTM1, nRBD-TM1, nRBD-TM1 plus the odor mixture *medicinal*, β2AR, SARS2-RBD, SARS2-RBD plus the odor mixture *medicinal*, Delta-RBD clone 1, Delta-RBD clone 2. Almost no Cherry/GFP double positive cells were observed for the two controls nTM1 and β2AR **(Figure 2A).** But greater than 60-fold double fluorescent cells were observed in the other four RBD conditions (56.8 ± 5.0-fold) or two RBD incubated with odor (55.9 ± 10.4-fold) **(Figure 2A).**

**Figure 2:**
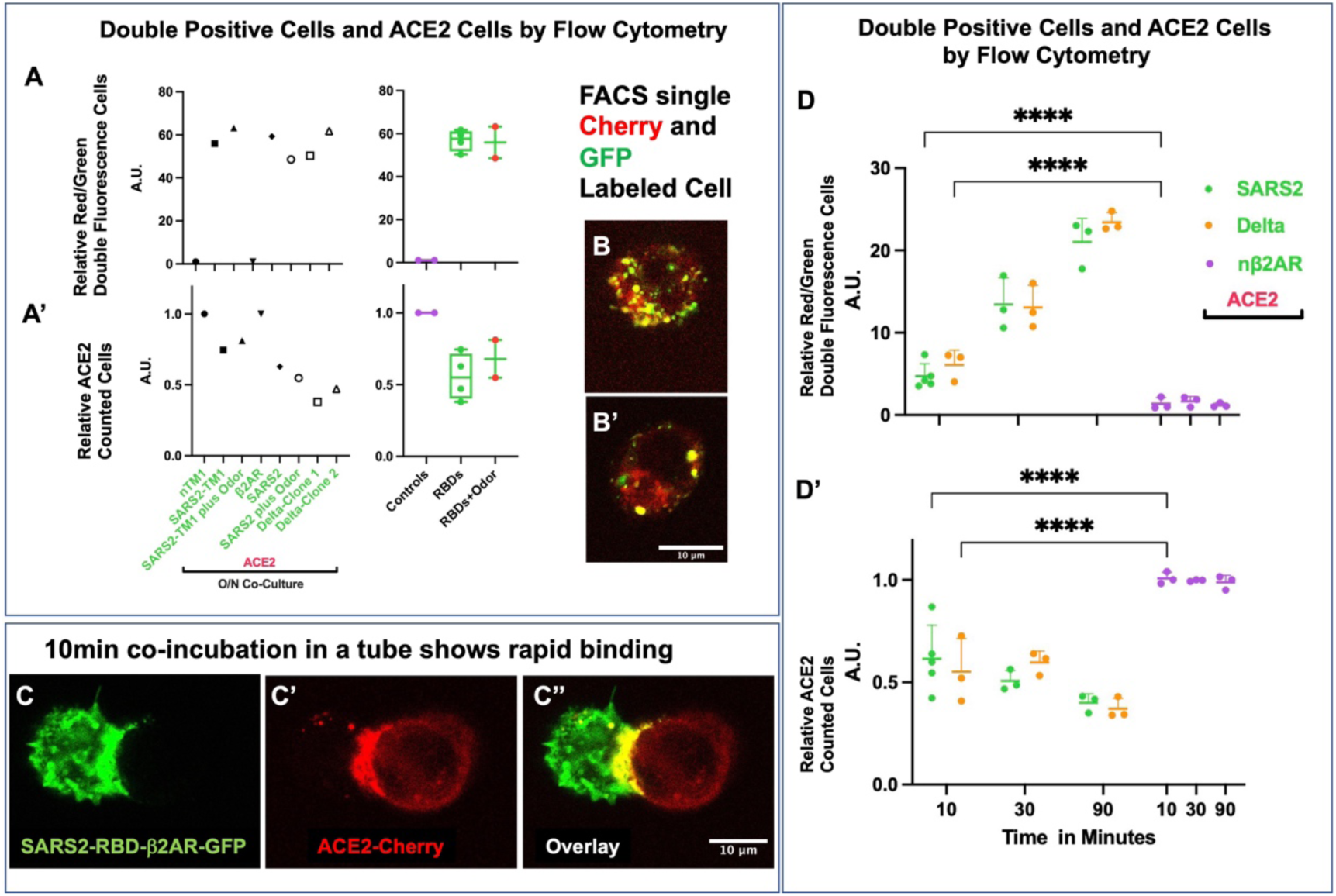
Quantifying RBD-ACE2 interactions by single cell flow cytometry analysis. **A and A’.** Initial experiments aimed a quantifying interaction. RBD-TM1/ RBD-β2AR-GFP fusion constructs and controls co-plated after separate electroporation (EP) and left to interact for 16hrs. nTM1, SARS2-TM1, SARS2-TM1 plus overnight (O/N) LNDV odor mixture 35 (1:500), β2AR, SARS2, SARS2 plus LNDV odor mixture 35 (1:500), and two clones of Delta (1 and 2). In all cases where an RBD was present, double fluorescent cells were observed at 60x WT values, which were in the single digits. Data were performed in triplicate, but when all the WT or RBD containing cells are analyzed in aggregate, then highly statistically significant data is obtained. Values are normalized to controls (Y-axis arbitrary units, A. U.). Surprisingly, ACE2 fluorescence was diminished in all cases RBD was present and again, when cells are analyzed in aggregate, data is highly significant. **B and B’.** Two double fluorescence cells shown by confocal imaging, revealing red cells with green speckles. **C-C’’.** *Dissociated Cell-based Assay* Confocal imaging of a dissociated green cell adhering to a dissociated red cell that are acutely mixed for 10min at 37°C; Many of the cells were found to interact with an overt relocalization of the ACE2-Cherry protein (**C’**). **D and D’** *Dissociate, Plate, Flow cytometry (DPF) Assay*: values in are normalized to controls (Y-axis arbitrary units, A. U.). The speed of the interaction observed in **C**, was recapitulated by layering dissociating EPd test constructs on ACE2-cherry cells in triplicate including for 3 time points: 10min, 30min and 90min, followed by dissociation and flow cytometry counting. ACE2-Cherry cells at all time points are from the same set of EPs were plated O/N on individual 30cm plates. GFP fluorescent cells, Control nβ2AR, SARS2-RBD and Delta-RBD cells were seeded onto sets of 30cm plates in triplicate. After O/N growth, individual 30cm plates of cells were trypsinized and plated on top of ACE2-Cherry cells for the defined time. Like the O/N co-cultures, the trypsinization-co-plating-flow cytometry analysis revealed double fluorescence in 10min as well as a near 50% reduction in ACE2-Cherry expressing cells

These pilot experiments clearly demonstrated that RBD-ACE2 interactions could be quantified and that the odor *medicinal* **did not** mitigate binding. From this flow cytometry experiment, I went on to acquire single Cherry cells, GFP cells, Cherry and GFP double fluorescent cells by FACS and microscopically analyzed each population: Green cells largely contained GFP fluorescence and only rare fragments of ACE2-Cherry fluorescence could be observed, Cherry only cells did have some green fluorescence and the double fluorescent cells showed red cells with robust green fluorescence **(Figure 2B, 2B’).** Flow cytometry experiments strongly argued that pieces of green-fluorescent membrane were being pulled from Delta-nRBD-β2AR-GFP expressing cells.

#### The RBD➔ACE2 in situ single cell binding assay

The robustness of the RBD-ACE2 interaction suggested that the observed binding may occur in a short period of time. To test this idea, trypsinized Delta-nRBD-β2AR-GFP expressing cells and ACE2-Cherry expressing cells were co-incubated in a tube at 37°C for 10 minutes, forming doublets of green and red fluorescent cells **(Figure 2C-C’’’)**. Within these doublets, the distribution of Cherry fluorescence (ACE2 protein) is no longer uniform and found at the edge of the cell, juxtaposed to the GFP-expressing cell.

#### Time course for RBD➔ACE2 interactions

Flow cytometry analysis of double fluorescent cells proved to be a robust, quantifiable method to determine the degree of interactions between RBD-expressing cells and ACE2-expressing cells. The speed of the interactions between these cells, led to a different version of the assay. Now, transfected cells are grown separately overnight, followed by trypsinizing the “test constructs” and plating them onto ACE2-expressing cells. A time course experiment was performed using triplicate platings of test constructs at 10, 30, or 90 minutes (min). Following this incubation, cells on target plate were washed, trypsinized, and quantified by flow cytometry analysis of single cells [*Dissociate-Plate-Flow Cytometry (DPF) assay*]. As expected, nβ2AR expressing cells did not interact with ACE2-expressing cells at any timepoint forming very few GFP/Cherry double fluorescent cells **(Figure 2D).** But, for both nRBD-β2AR (SARS2-RBD) and Delta-nRBD-β2AR (Delta-RBD) expressing cells, 4.7 and 6.1-fold increases (± 1.5 and ± 1.8, respectively) over background in GFP/Cherry double fluorescent cells were observed in 10minutes (min), respectively; 13.4 and 13.1-fold increases (± 3.2 and ±2.7, respectively) in 30min, respectively; and 21.0 and 22.9-fold increases (± 2.8 and ±1.2, respectively) in 90min, respectively **(Figure 2D).** Cell seeding is generally a slow process that takes hours, so the ∼5-fold increase in double fluorescent cells at 10min was remarkable. When quantifying the ACE2-cherry fluorescent cells above background, a noticeable decline in the number red cells at 10 min was observed reaching ∼3-fold fewer numbers by the 90 min timepoint, a ∼60% reduction ±5% **(Figure 2D’).** A 40%±16% reduction of ACE2 cell counts was observed in the pilot experiment as well **(Figure 2A’)**. The rapid drop in ACE2 levels suggests either red cells have died in the 90 min period perhaps due to reorganization of membrane proteins, or ACE2 protein is being rapidly degraded through RBD binding. A decrease in ACE2-Cherry fluorescence was observed in both Cherry only and GFP/Cherry double fluorescent cells incubated with the RBD expressing cells suggesting that ACE2-Cherry fluorescence is being reduced on a per cell level. Utilizing this new assay, it was confirmed that neither Secreted-SARS2 nor Secreted-Delta gave rise to GFP/Cherry double fluorescent cells **(Figure S2A)**, nor reductions in ACE2-Cherry cell counts **(Figure S2A’)**. Several control experiments further validated the assay: 1) Reducing the number of Delta-RBD cells added to ACE2-expressing cells did not alter the effect; 2) Delta-RBD cells did not interact with β2AR-Cherry cells; and most importantly the Delta-RBD cells did not cause the ACE2-expressing cells to be dislodged from the plate as GFP/Cherry doublets were not observed in the supernatant after a 90min co-culture **(Figure S2A and S2A’)**. Finally, the supernatant from Delta-RBD cells grown overnight did not produce any GFP/Cherry doublets nor a decrease in ACE2-Cherry fluorescence **(Figure S2B and S2B’)**. A direct test of the secreted particle possibility utilized purified His tagged-RBD protein coupled to an anti-His-488 antibody and incubated for 90min with ACE2-Cherry expressing cells. No impact could be observed on the downregulation of ACE2-Cherry fluorescence despite the formation of antibody labeling of RBD on ACE2-expressing cells (**Figure S3A and S3A’**). In sum, there was no evidence of sudden cell death nor secreted particles from the cells expressing a tethered RBD. Based on these observations, it’s unlikely that the non-membrane-bound Spike S1 fragment would lead to downregulation of ACE2-Cherry.

#### Does cell adhesion cause downregulation of membrane proteins?

Thus far, the data show specificity between nRBD-β2AR-GFP expressing cells and ACE2-Cherry expressing cells. Yet, nonspecific interactions and ACE2-Cherry protein could be due to β2AR interaction or through the effects from an unidentified process of cell-cell adhesion. As a control Venus or Cherry fusions of the cell adhesion molecule Cadherin 2 (Cdh2) were separately transfected and co-cultured overnight in OP6 cells producing robust cell-cell contacts compared to Delta-RBD and Cdh2 **(Figure S2B)**. Interaction of Cdh2 was observed as double fluorescent cells, however the red fluorescence of CHD2-Cherry was not decreased (**Figure S2B** and **S2B’**). To rule out the RBD-ACE2 interactions were phenocopying a cell adhesion phenomenon, a series of controls using ACE2-Cherry, β2AR-Cherry, Cdh2-Cherry cells were co-incubated for 90min with β2AR-GFP, Delta-nRBD-β2AR-GFP, or Cdh2-GFP cells, followed by flow cytometry analysis; Only Delta-nRBD-β2AR-GFP and ACE2-Cherry cells led to double fluorescent cells and decreases in Cherry fluorescence of ACE2-Cherry (**Figures S2C, S2C’**).

Double fluorescence cells between ACE2-Cherry and Delta-RBD GFP cells was more substantial than the adherence of Cdh2-GFP + Cdh2-Cherry cells in 90min co-cultures **(Figure S2C and S2C’).** Here it can be firmly concluded that the observed cell dynamics are derived from RBD and ACE2 binding.

#### RBD➔ACE2 membrane reorganization

We have previously shown that plasma membrane dynamics can be readily monitored by an array of palmitoylated fluorescent proteins [20-residue sequence (aka *gap*) from Zebrafish gap43] (20). To determine if the OP6 cells were undergoing general membrane changes from the RBD-ACE2 binding, ACE2-Cherry and gapTeal were cotransfected (Co-EP) and Delta-nRBD-β2AR-Venus cells were added and imaged after 30min **(Figure S4A-D’’’)**. Reorganization of the ACE2-Cherry fluorescence and Delta-nRBD-β2AR-Venus was readily observed, but not that of gapTeal **(Figure S4A, A’)**. Quantification of fluorescent pixels showed co-accumulation of Venus and Cherry fluorescence at the plasma membrane, but not Teal **(Figure S4B)**. Unfortunately, quantification of these results by single cell flow cytometry is not possible because Teal and Venus fluorescence are not separable by our available flow cytometry machines. But flow cytometry analysis of single cells can be performed with GFP, Cherry and Farred fluorescent proteins. To circumvent this issue, the assay was reorganized. Now ACE2-GFP and gapFarred [codon optimized AFP (20)] were Co-EPd. Separately transfected Delta-nRBD-β2AR-Cherry, Cdh2-Cherry or nβ2AR-Cherry were followed by trypsinization, co-plating and imaging within 30min over time **(Figure 3A-C’’’)**. Quantification of fluorescent pixels again showed co-accumulation of GFP and Cherry fluorescence at the plasma membrane, but not Farred **(Figure 3D)**. It is notable that the effects on membrane reorganization of ACE2 protein was independent of Ct fluorescent protein fusion: GFP fluorescence also appeared weaker suggesting it was being degraded. Flow cytometry analysis on single cells and confirmed the specific interaction of Delta-nRBD-β2AR-Cherry with ACE2-GFP showing 37.8 ±6.1-fold more double positive cells than controls and a 47±4.3% reduction of ACE2-GFP fluorescence; Control proteins nβ2AR-Cherry and Cdh2-Cherry did not associate with ACE2-GFP nor cause its downregulation **(Figure 3E and 3 E’)**. Furthermore, gapFarred fluorescence was not reduced to the same extent as GFP fluorescence **(Figure 3E’)**.

**Figure 3:**
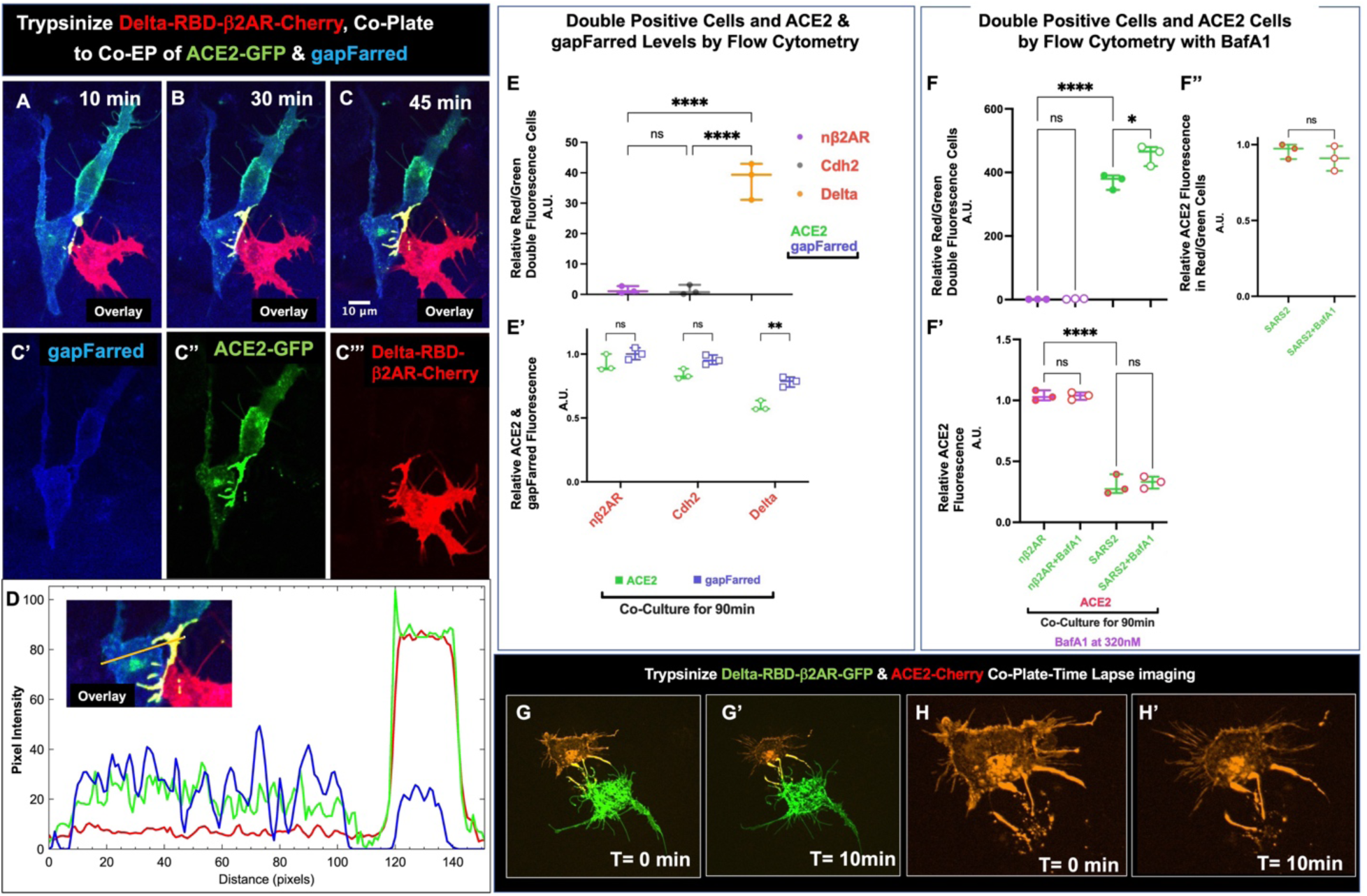
RBD-Cherry affects ACE2-GFP, but not gapFarred (see also Figure S4) **A-C.** Confocal images for Co-EP of ACE2-GFP and gapFarred. After 16hrs Delta-RBD-Cherry cells are trypsinized and plated onto ACE2/gapFarred cells. **A-C’’’**. In less than 10min after plating, aggregation of both Delta-RBD and ACE2 fusion proteins is observed, but not gapFarred (a M71Q mutation in Cherry was utilized to reduce detection in. Farred spectrum). **D**. Histogram of fluorescence intensity changes in **C-C’’’** also reveals aggregation of Delta-RBD and ACE2 fusion proteins, but not the membrane localized protein gapFarred. **E and E’** *DPF Assay*: values in are normalized to controls (Y-axis arbitrary units, A. U.). Flow cytometry analysis of the Delta-RBD plated cells onto double fluorescent ACE2-GFP/gapFarred cells for 90min reveals double fluorescent cells (Red/Green) and a reduction of ACE2 fluorescence (Green) relative to two controls, nβ2AR and Cdh2. gapFarred fluorescence did show reduced expression in the context of Delta-RBD, but not to the same extent. **F. F’.** Spinning disk confocal imaging of O/N EPd ACE2-Cherry and Delta-RBD- nβ2AR-GFP cells coplated for 10min, which reveals altered morphology of the red fluorescent cells, but not the green, fluorescent cells (**see also Movies S1A-C**).

The assay developed here to study RBD/ACE2 interactions were not developed to study the previously shown of ACE2-internalization and loss. Here, SARS2-RBD interactions with ACE2 occur without the presence of TMPRSS2 nor S2’ fusion protein to facilitate receptor-mediated endocytosis. As such, lysosomal cathepsins would likely be required for endocytic entry and loss of ACE2 as is found for SARS2 entry in TMPRSS2 deficient cells (36-38). There are many mechanisms that could be involved in endocytosis and eventual reduction of ACE2 fluorescence in addition to Clathrin-mediated processes (39, 40). To rule in/out two of the possibilities, it was determined if the proteasome inhibitor MG-132 or the lysosome inhibitor bafilomycin A1 (BafA1) could interfere with ACE2 internalization. Neither MG-132 (**Data File S2**) nor BafA1 (**Figures 3F’-F’’**) affected ACE2-cherry fluorescence in the context of SARS2-RBD, but more double positive cells were observed with the BafA1 treatment (**Figure 3F**) suggesting an increased availability (less loss) of ACE2-Cherry for binding to SARS2-RBD. Interactions between SARS2-RBD and ACE2-Cherry expressing cells over 90min in the presence of lysosomal inhibition did not prevent ACE2-Cherry loss nor inhibit the formation of doubled positive cells. Perhaps lysosomal inhibition of ACE2-Cherry loss would be observed in an earlier time point. It should be noted that ACE2-expressing cells counts are derived from expression levels above background (**see Figure 2D’)**, ACE2 fluorescence levels above background have also been calculated **(see Figures 3E’, 3F’-F’’, and S3A’, S3B’’, S3C’’)**. The ACE2 internalization effects were even more profound after overnight co-plating **(Figures S4C-C’’’)** compared to controls **(Figures S4D-D’’’)**, which showed no internalization of ACE2. It should be noted that internalization of ACE2-Cherry did not lead to cell death. Finally, spinning disk imaging of the cell-based interactions with Delta-nRBD-β2AR-GFP and ACE2-Cherry cells were performed to show real-time dynamics; ACE2-Cherry reveal drastic reorganization, but not Delta-nRBD-β2AR-GFP cells within 10 minutes of imaging performed immediately after co-plating **(Figures 3G-G’, 3H-H’, and Movies S1A-C)**. In sum, RBD specifically reorganizes and degrades ACE2-fluorescent proteins.

#### Meta-Analysis of 197 residue SARS2-RBD domain

This new cell-based RBD-ACE2 interaction assay provided a novel framework to define critical RBD binding residues. Previously, the SARS1-RBD (residues 320-515) defined a 74% identical region for SARS2 (residues 333-529) **(Figure 4)**. A subset of residues that provide structural integrity and binding potential were identified by reviewing the crystallographic data (1, 28), deep mutational screening (DMS) (25, 41), in silico predicted interactions (42) and the known properties of amino acids in proteins.

**Figure 4:**
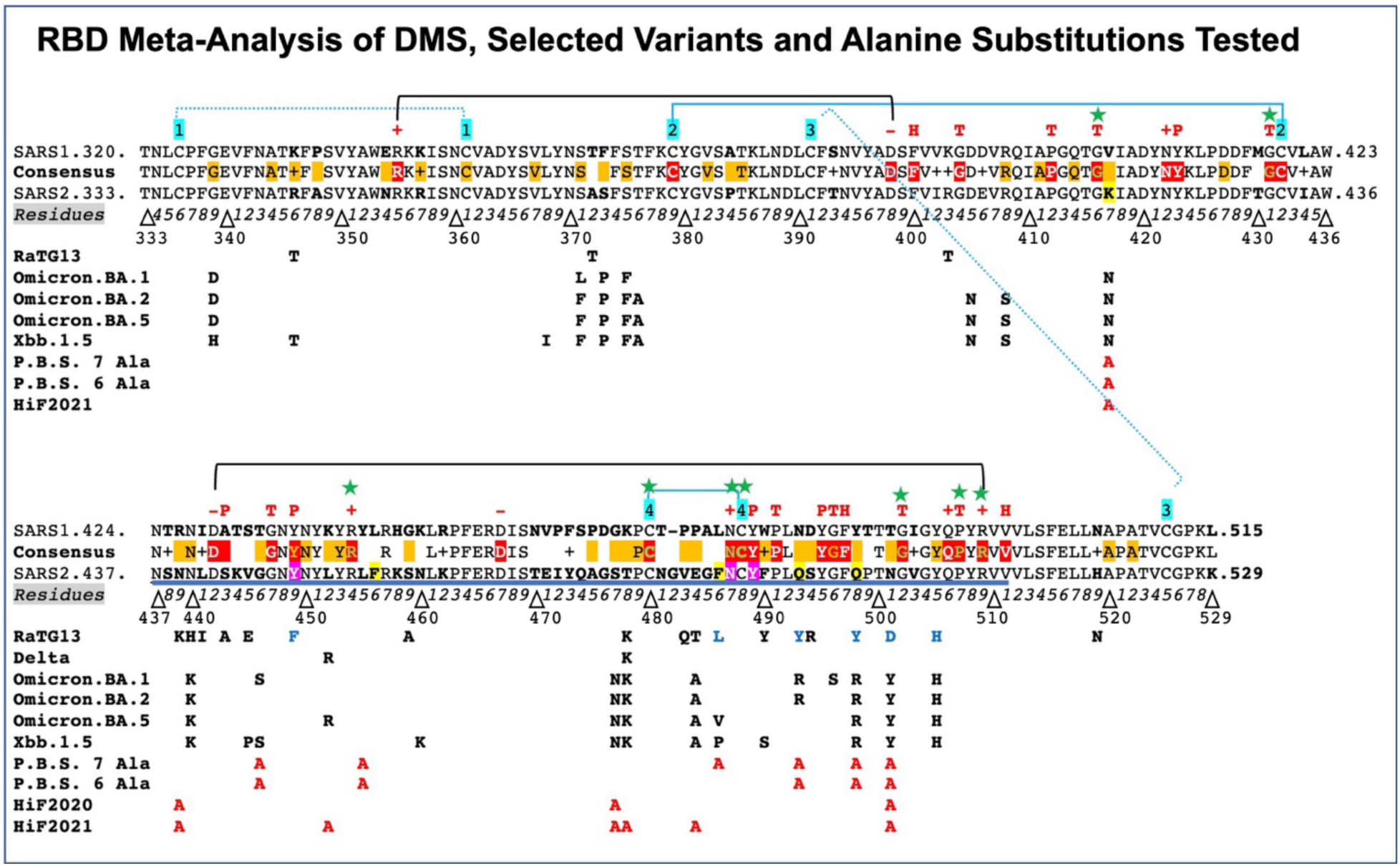
RBD Meta-Analysis of Deep Mutational Screening, Identified Variants, Mutations Tested. **Alignment of SARS1 and SARS2 reveals 74% amino acid identity (145 residues) over the 197 residues of the Receptor Binding Domain (RBD). SARS2 Residue numbers from 333 to 529 are shown.** Consensus residues are shown between SARS1 and SARS2: The + denoting similar amino acids (15 residues). Eight conserved Cysteine residues denoted by blue numbers and brackets with their proposed linkages: 1-1 (SARS2,336-361), 2-2 (SARS2,379-432), 3-3 (SARS2,391-525), 4-4 (SARS2,480-488) (99). Meta-analysis of Deep Mutational Screening of RBD reveals 29 residues (Red highlighted residues in the consensus) that are retained in 70% of the mutations and thus deemed immutable (25, 41). Orange highlighted residues or shaded regions in the consensus denote where mutations are found in SARS2 at a frequency greater than 0.03% or found in the omicron variant (http://cov-glue-viz.cvr.gla.ac.uk/mutations.php through 2021)(43); two residues are labelled with orange lettering as they reside among the highly conserved red highlighted residues, **Y449** and **G496**). In contrast to the variant residues, 9 red highlighted residues with green lettering, also denoted with green stars (G416, G431, R454, C480, N487, C488, G502, P507, R509) were deemed absolutely necessary, which include one pair of Cysteine residues C480-C488, three Glycines and one Proline that promote Beta turns and two Arginines that likely contribute to RBD structural integrity through interactions with conserved Aspartic Acid residues: black brackets (R355-D398 @2.8Å; D442-R509 @3.1Å; R454-D467 @3.4Å)(100) (100)**. Most of the red highlighted residues promote structural integrity of the RBD domain:** Red highlighted Glycines and Prolines are denoted with a red T, charged residues are denoted with red + or -, and hydrophobic residues are denoted with red H and polar Tyrosine/Serine with a red P. **Receptor Binding Motif (RBM):** N437-R510 is underlined in SARS2. Amino acid differences are bolded with 5 Yellow highlighted residues (K417, F456, F486, Q493, Q498, N501) in SARS2 have been modeled by density function calculation to be the key residues that SARS2 tightly bind to ACE2 compared to SARS. Three residues Y449, N487 and Y489, highlighted in purple for SARS2 are predicted to be critical for strong association with ACE2. **RBD Variants Analyzed:** RaTG13-RBD (blue residues predicted to be relevant for binding) (54), Delta variant-RBD **B.1.617.2**, Omicron variant-RBD **B.1.1.529 (OM.BA.1)**, Omicron variant-RBD **B.1.1.529.2 (OM.BA.2), OM.BA.5,** and **XBB1.5.** **Multiple Alanine Mutants analyzed to look at critical residues for RBM-ACE2 interaction: Predicted Binding Sites (P.B.S.):** K417A, G446A, L455A, F486A, Q493A, Q498A, N501A, alters 4 of the modeled residues (bold), plus two additional residues implicated (28, 42). **P.B.S. 6Ala:** returns F486 within Disulfide Loop 4 (480-488). **HiFrequency variations in 2020 SARS2 variants:** N439A, S477A, N501A **HiFrequency variations in 2021 SARS2 variants**: K417A, N439A, L452A, S477A, T478A, E484A, N501A

DMS by two groups led to the identification of clones that can still bind ACE2. Both groups produced very similar data, but one group also controlled for RBD stability revealing 29 residues that were not mutated in 70% of the clones and thus necessary for binding **(Figure 4, red highlights within SARS1-SARS2-RBD consensus sequence)**. Ten of the 29 residues were distributed and not implicated in the folding of the RBD and thus candidates for ACE2 binding: F400, N422, Y423, S442, N487, Y489, Y495, F497, Q506, H511. Nine of the 29 residues were retained in 90% of the DMS clones **(Figure 4, green lettering and green starred)** and would appear to be critical for function including R454, N487, G502, P507 and R509 as well as the four Cysteine (Cys) residues that form disulfide bridges in Loop 2 and Loop 4. The greatest concentration of immutable residues lay between N487 and V511, which is consistent with three residues described in the literature predicted to affect binding when mutated (Y449, N487, Y489, **Figure 4** purple highlighted)(43).

As an apparent contradiction, residues predicted in silico to be critical for binding **(Figure 4, yellow highlights)** do not overlap any of the 29 relatively immutable residues from DMS clones: K417, F456, F486, Q493, Q498 (42). Among various variants produced in first two years of the pandemic only two residues (Y449, G496) are amongst the 29 residues. Finally, a set of three residues that encompass C488 from disulfide Loop 4: N487, C488, and Y489 are well conserved in all known betacoronaviruses that bind ACE2 **(Figure S5)**. Taken together, most interacting residues appear distributed, except for a stretch of residues in proximity to C480 and C488.

It is known that the alphacoronavirus H-CoV-NL63 has a 136 residue RBD domain that also binds ACE2 (NL63-RBD). This domain has only eleven residues conserved with SARS2-RBD, but four of them are colinear with the RBD disulfide Loop 4 region **(Figure S5 and S6; residues C-FPL)** (44). The NL63-RBD homology region is not among the residues predicted to interface with ACE2 **(Figure S6; RBM1-3)**, but it is interesting to note that the N578Y mutation that makes a five-residue match to the SARS2-disulfide Loop 4 region (CYFPL) increases its binding potential to ACE2 **(Figure S5)**(44). By contrast, the SARS2 Spike homologue, RpYN06, is missing the region between C480 and C488 that comprises disulfide Loop 4 and fails to bind ACE2 **(Figure S7)**(45-47). Thus, a strong argument can be made for critical ACE2 binding residues reside within the RBD-disulfide Loop 4 region including the pair of Cys residues.

#### RBD and disulfide Loop 4 region domain mutagenesis

To better understand the functionality of residues within the RBD domain, an array of RBD variants and mutants including disulfide Loop 4 substitutions were tested to characterize RBD binding to ACE2. The SARS1, SARS2 and NL63 RBDs have been shown to bind the ACE2 protein with Kds in the low nM range **(Figure S5)** (1, 44, 48). SARS2 variants have not been compared yet but are likely in the same range. One RBD, RaTG13 binds with a Kd in the μM range (1000-fold less) (**Figure S5)** (26). All four orthologues and five SARS2 variants (Alpha, Delta, Delta-Plus, Omicron-BA.1, Omicron-BA.2) were tested in our DPF assay **(Figures 4, 5, S8)**. No differences were observed in their ability bind and reduce ACE2-Cherry expression, except for NL63, which produced fewer double positive cells but more severe loss of ACE2-Cherry expressing cells **(Figures 5A, 5A’ and S8B’)**. Alanine substitutions were used to simultaneously convert all 3 high frequency mutants (>.03%) found in year 2020 (HiF2020) and all 7 residues found in year 2020 and 2021 (HiF2021). None of these mutations altered binding and reduction of ACE2-Cherry expression shown with SARS2-RBD **(Figure 5A, S8B, B’**). Next, seven Predicted Binding Sites (P.B.S.; K417, G446, L455, F486, Q493, Q498, N501) between RBD and ACE2 were converted into alanine as a group (P.B.S. 7Ala-RBD-β2AR-GFP). The P.B.S 7Ala-RBD trafficked to the plasma membrane but was not observed to bind ACE2-Cherry cells **(Figures 5A, 5B, S9A, S9A’, S10B)**. This mutant was the first that completely blocked binding **(Figures 5)**, a 6.1±3.1-fold double positive rate compared to 3.0±1.8-fold in controls. One of the alanine substitutions was within the disulfide Loop 4, F486, might have been predicted to be critical for RBD-ACE2 interactions. Indeed, the P.B.S. 6Ala mutant with the A486F revertant could now bind to and reduce ACE2-cherry fluorescent cells **(Figures 5, S9B, S10C)**, 85.6±10-fold double positives, almost as well as SARS-RBD **(Figure S9C and S10A)** with 101.2±13.0-fold double positives. Other single reversions were not performed, but due to the distributed nature of binding; It is plausible that several might also rescue binding. Among the 7 residues encompassing the P.B.S., it is likely that no one residue is critical for binding except F486. Yet, this residue could be heavily mutated in the DMS experiments, suggesting that it does not form bonds that are necessary and sufficient of RBD binding. Taken together, DMS mutagenesis along with the vast number of residue differences between RaTG13/SARS1 and SARS2 along with the multiple alanine mutations suggested that critical residues for binding are distributed throughout the RBD.

**Figure 5:**
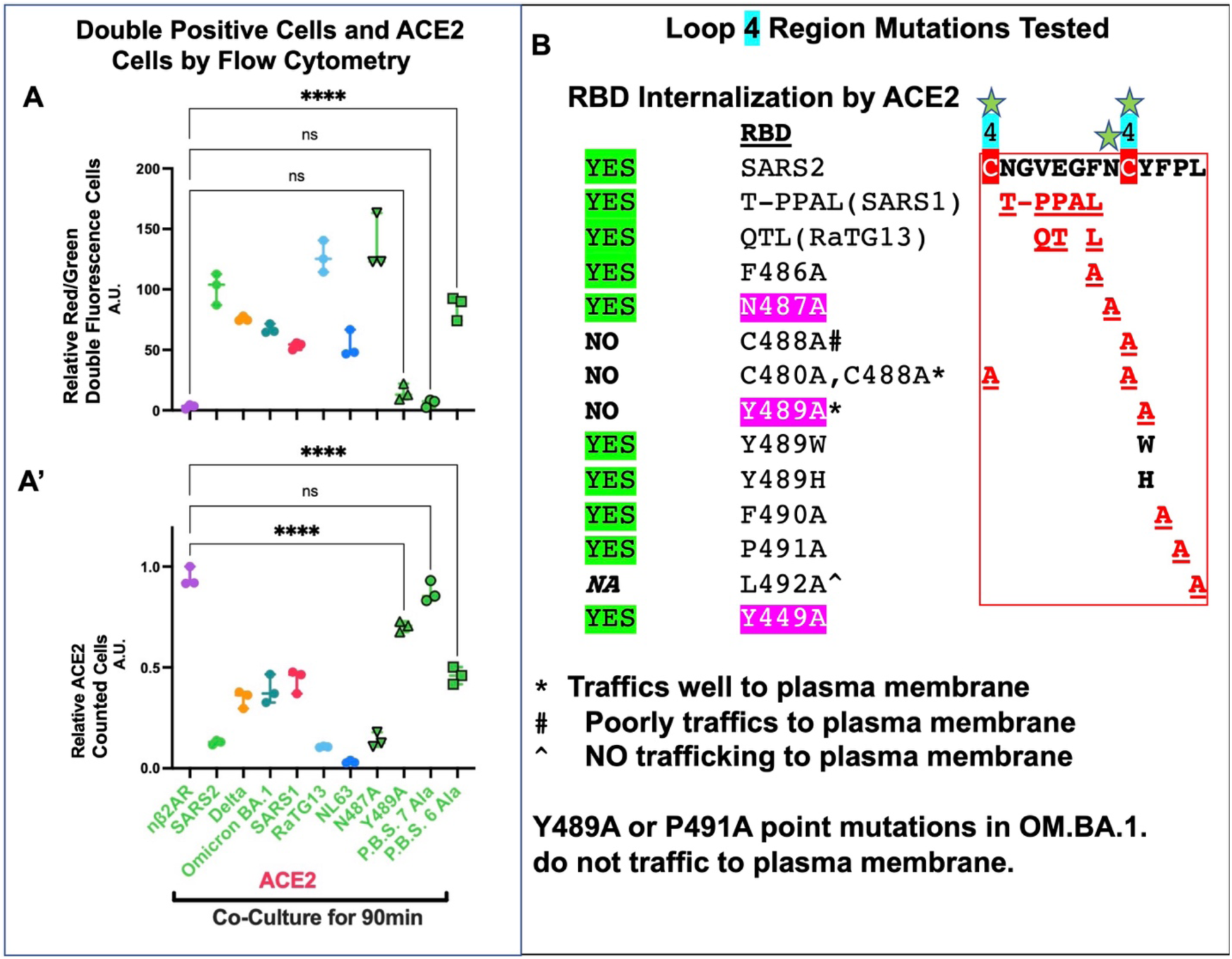
RBD Variants, Orthologues and Loop 4 mutants. **A, A’. RBD Variants and Orthologues:** *DPF assay*: values are normalized to controls (Y-axis arbitrary units, A. U.). Three Variants of Concern were tested for their ability to bind and be internalized by ACE2: Delta, OM.BA.1 and OM.BA.2 (not shown) and two RBD orthologues SARS1 and RaTG13 tested. All 5 of these RBDs were internalized by ACE2 in an indistinguishable manner, despite the measured binding of RaTG13 being 1000-fold less than SARS1 and SARS2. Four multiple alanine mutants were also tested (HiF2020, HiF2021, P.B.S. 7Ala, P.B.S. 6Ala). Neither the three-alanine mutation in HiF2020 nor the seven-alanine mutation in HiF2021 had any effect on RBD internalization by ACE2-expressing cells. In contrast, the seven-alanine mutation in P.B.S. did abolish RBD binding. When F486A was reverted (A486F), RBD internalization by ACE2-expressing cells was readily observed. The divergent RBD orthologue in H-CoV-NL63 could function in the in vitro plate binding assay. NL63-RBD (136 residues) is known to bind to similar domains on ACE2 through three non-contiguous RBM regions (**See Figures S5, S6**). Indeed NL63-RBD was very efficient at being internalized by ACE2-expressing cells. Single cell flow cytometry binding assay scoring for reduction of ACE2 levels is inverse proportional to the formation of Red/Green cells. NL63-RBD was more robust at downregulating ACE2 than all other RBDs tested, which also accounts for the reduced number of Red/Green single cells. **B. Loop 4 mutants:** Meta-Analysis from DMS suggested that key Loop 4 Cysteines, N487 and Y489 were critical for RBM interaction with ACE2. Focusing in on Loop 4, residue substitutions in SARS2 that match SARS1 and RaTG13 are characterizing. In addition, alanine scanning substitutions between F486 to L492 are characterized including Y449A for comparison. In O/N co-plating of constructs or by *DPF assay*, nearly all RBD orthologues, Loop 4 substitutions, and alanine substitutions were internalized by ACE2-expressing cells except for three: C488A and L492A, neither of which showed significant trafficking to the plasma membrane (PM) and Y489A, which robustly trafficked to PM, bound, but was not internalized by ACE2-expressing cells **(Figures S8, S9)**. The single cell flow cytometry binding assay results parallel the qualitative in vitro plate binding assay showing absence of Red/Green cells for P.B.S. mutant and near absence for Y489A. The P.B.S.+F486 nicely shows rescue. Based on cross species conservation, both Y489A and P491A were made in the OM.BA.1 variant, but unlike SARS2, both were unable to traffic to the PM **(Figure S10)**.

If RBD binding to ACE2 were distributed, then how are nanomolar Kds accomplished? Cross species sequence comparisons suggest disulfide Loop 4 may be key to nucleate ACE2 interactions. It is known that a small tripeptide domain like Arg-Gly-Asp (RGD) can be enough to achieve high affinity binding to another protein (49). Thus, a small motif within the RBD domain may be all that is necessary, but no such 100% conserved three residue region exists amongst RBD domains and mutants. Yet, the disulfide Loop 4 represented an interesting region as the disulfide bond itself may play a critical role in recognition of the ACE2 protein **(Figure 5B)**. In the context of SARS2-RBD, small regional substitutions between C480 and 488 were generated: SARS1 481-486 and RaTG13 483-486 substitutions as well as scanning alanine mutants for residues 486 through 492 **(Figures 5B and S8B**). Neither SARS1 and RaTG13 Loop 4 substitutions nor alanine mutants within this region showed any differences in binding and internalization **(Figure 5)**. Based on the P.B.S. 6Ala-RBD, the F486A alone might have disrupted binding, but it did not **(Figures 5)**, again pointing to the distributed nature of RBD binding to ACE2.

Alanine scanning of neighboring residues Y489, F490, P491, L492 and the Y449 residue predicted to be important for binding revealed four interesting aspects of RBD binding to ACE2. First, only Y489A was able to nearly abolish binding (14.7±6.6-fold double positives) and reduced the loss of ACE2-Cherry fluorescence cells (30%±2.6%), while still being well expressed at the plasma membrane (**Figures 5, S8, S9, S10**). In contrast, the Y449A residue predicted to decrease binding of RBD had no effect. Second, C488A did not traffic well to the plasma membrane, but even when it did, binding to ACE2-Cherry cells was not readily observed. It seemed likely that the C488A mutation that poorly traffics RBD to the plasma membrane (PM) may be due to the C480 residue forming a disulfide bond with one of the other 6 Cys residues **(Figure 4).** To mirror the loss of both Cys residues found in RpYN06 **(Figure S7),** the double mutant C480A, C488A-RBD was generated. This mutation traffics very well to the PM, on occasion binds ACE2-Cherry expressing cells, but does not become internalized **(Figures 5, S9)**. Third, neither N487A, F490A nor P491A-RBDs affected PM trafficking nor ACE2 binding/degradation properties of RBD. Finally, mutational analysis was expanded to the Omicron (OM).BA.1 variant for Y489A and P491A mutants. But neither mutant could traffic to the PM suggesting that OM.BA.1 had a different confirmational landscape to that of SARS2 or any previous variant **(Figures 5 and S10)**. The failure of Y489A-OM.BA.1-RBD mutation to allow PM trafficking suggests a disruption of C480-C488 disulfide bond formation and further points to an altered Loop 4 confirmation in the Y489A-SARS2-RBD mutation. Overall, the data strongly implicate the formation of the disulfide Loop 4 bond as a critical feature of RBD-ACE2 interactions.

#### Blocking RBD binding using a disulfide reducing agent

The analysis RBDs from NL63 and betacoronaviruses such as SARS1, SARS2 and its variants define a continuum of RBD residues necessary for allowing its internalization by ACE2. These data argue that it would be difficult to find an airborne molecule, which would specifically disrupt contacts between the changing landscape of RBD and ACE2 interactions. But the disulfide Loop 4 analysis suggested that critical Cysteine residues involved in disulfide bridges were common to all RBD domains could be a targeted to mitigate infection. That said, the volatile reducing agent Dithiothreitol (DTT) was able to disrupt SARS2 pseudotype virus infections on ACE2-expressing cells in the millimolar range (29, 50). DTT has a strong characteristic odor with a modest vapor pressure of 1.28 x10^-4^ mm Hg at 25C. A derivative of DTT, dithiobutylamine (DTBA) was identified in 2012 that had similar properties, but more resistant to changes in pKA and reduces disulfide bonds in small proteins more quickly (51). The RBD-ACE2 solution binding assay (**Figure 2C-C’’’)** was utilized to assess GFP-Cherry in concentration series (0mM, 0.5mM, and 2mM) for DTT and DTBA. SARS2-RBD-β2AR-GFP or OM.BA.1-RBD-β2AR-GFP and ACE2-Cherry GFP/Cherry doublets were readily observed in the 0mM DTT and DTBA conditions for both SARS2 and OM.BA.1-RBD (**Figure 6A**). In the 2mM DTT condition it was difficult to find SARS2-RBD or OM.BA.1-RBD bound to ACE2 (Figure **6B** **and inset**). In contrast, RBD interactions were not observed in 2mM DTBA. The effect of reducing agents on SARS2 and OM.BA.1-RBDs in the solution binding assay mirrored what was already tested using pseudotype viruses, showing that binding assay strategy could be set up to screen for other volatile compounds for their ability to disrupt disulfide bonds. Finally, the NL63-RBD with a different set of disulfide bonds was quantified in the presence of 2mM DTT and 2mM DTBA in the plate binding assay with ACE2-Cherry. 2mM DTBA completely removed the ACE2-Cherry cells from the tissue culture plate, but the 2mM DTT did not, and these interactions were quantified by flow cytometry; the 2mM DTT prevented double fluorescent cells from forming and prevented downregulation of ACE2-Cherry fluorescence (**Figure 6C and 6C’**). Thus, a limited exposure of disulfide bonds in RBDs using a volatile disulfide reducing agents could provide a new strategy to mitigate viral infections.

**Figure 6:**
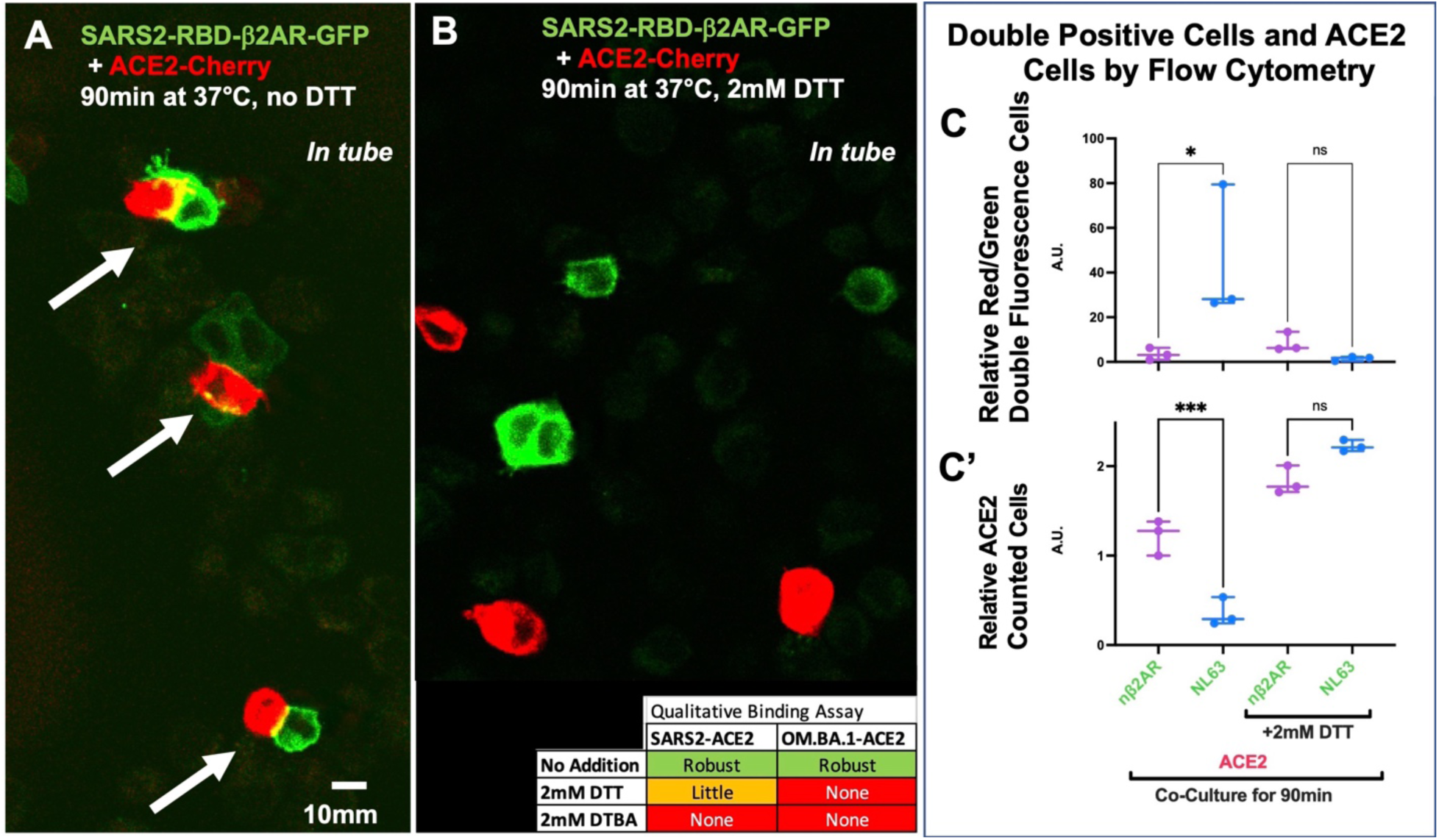
Disulfide Bond Reducing Agents Block RBD-ACE2 Association. **A, B.** *Dissociated Cell-based Assay*: SARS2-RBD (Green) or ACE2-Cherry (Red) by low magnification confocal microscopy which affords visualization of 10-15 Green and Red cells. Qualitative analysis of binding was performed after 90min incubation at 37°C for SARS2-RBD. Red/Green doublets (white arrows) were readily observed in at least 3 fields of view. Addition of 2mM DTT limits this interaction where doublets are rarely found. In contrast 2mM DTBA abolished the visualization of doublets (see inset). Both 2mM DTT and 2mM DTBA abolished doublet visualization for OM.BA.1-RBD as well. **C and C’** *DPF assay*: values are normalized to controls (Y-axis arbitrary units, A. U.). NL63-RBD binding to ACE2 was tested by 90min incubation. NL63-RBD shows both increased numbers of Red/Green single cells, but also a decrease in ACE2 expression. By contrast addition of 2mM DTT abolished Red/Green single cells and ACE2 expression levels were not reduced.

### ACE2 Mutants

#### ACE2 Amino terminal domain mutagenesis

SARS2-RBD is predicted to bind many residues on ACE2 through mostly hydrogen bonding (1, 28). These interaction domains can be separated into Nt proximal region: residues 19-45; Nt distal region: residues 79, 82-83, Internal region 1: residue 330, and Internal region 2: residues 353-357. Both RaTG13 and SARS1 RBDs are predicted to bind the same three regions with some variation in specific residues **(Figure 7A)**. The ACE2 Nt proximal region is poorly conserved (<50% homology) across 408 species from residues 19-27 but becomes increasingly more conserved from residues 28-45 (>50% homology)(52). By contrast, the ACE2 Nt distal region retains conservation only in residue Y83 (>50%) and the Internal region residues of 330 and 353-357 are also highly conserved (>50%) (**Figure 7A, B**). In comparison, NL63-RBD has reduced contacts with the divergent Nt proximal region, no contacts with the divergent Nt distal region, and expanded contacts with the conserved Internal region 1 (27). The regional overlap in binding to ACE2 for NL63-RBD and SARS2-RBD (SARS1 and RaTG13 as well) is remarkable despite the near absence of protein homology. The disulfide Loop 4 region of SARS2 is strongly implicated to bind only the ACE2 Nt proximal and distal regions (**Figure 7B)**. Mapping studies show that both N487 and Y489 are in involved in coordinating binding at both the Nt proximal and distal regions (**Figure 7B**), suggesting that disruption of this contacts might abolish binding (1).

**Figure 7:**
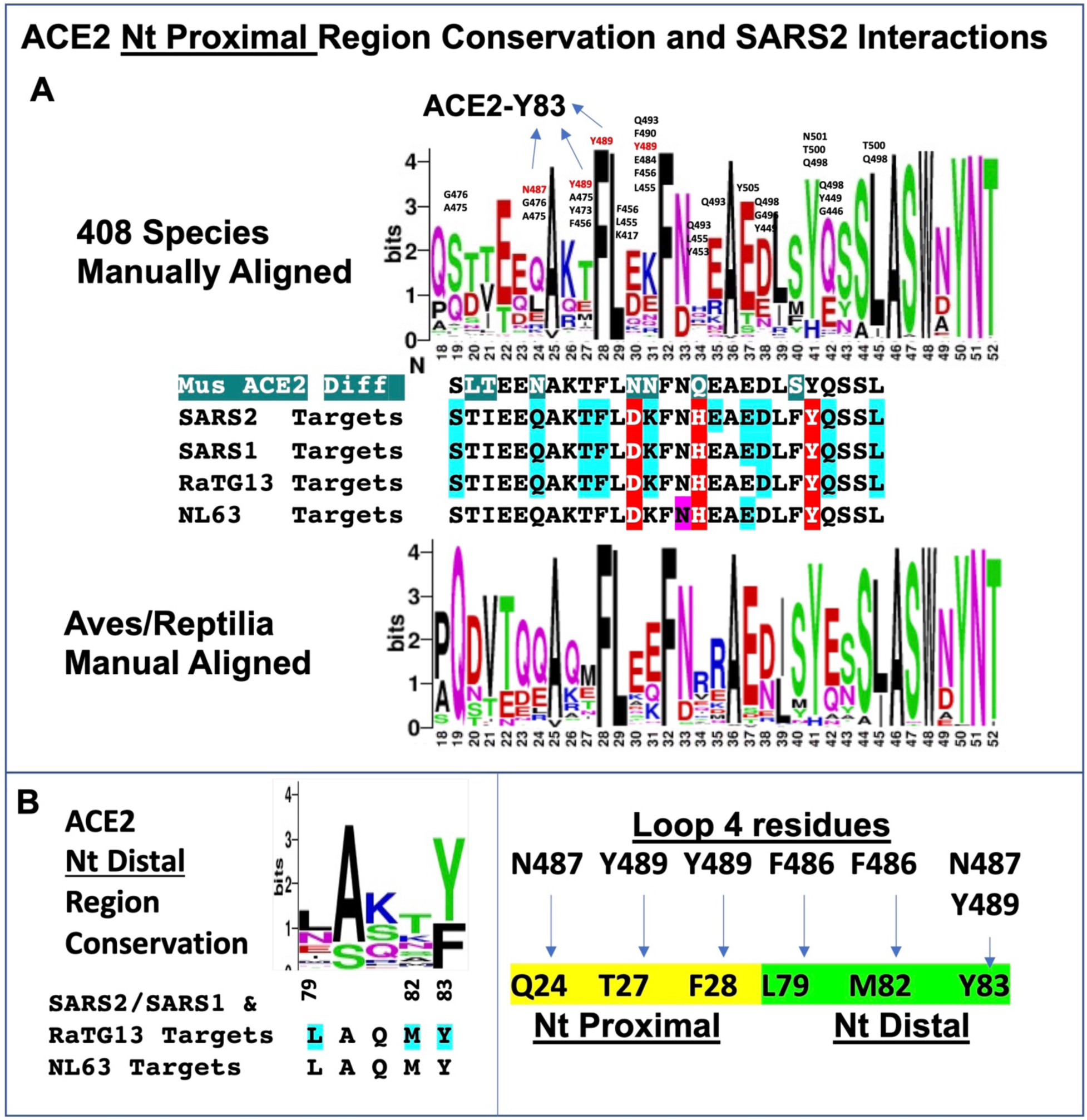
Conservation of ACE2 N-terminus and Predicted SARS2 Contact Sites. **A. ACE2 Nt. Proximal Region**. Logo plot of 408 ACE2 species (52) manually aligned starting from residue 18, the last residue before the signal peptide is spliced. ACE2 conserved residues between 22-41 appear to have a periodicity consistent with forming an alpha-helix whereby they would nominally be on the same face. SARS2 point of contacts are predominantly to the non-conserved ACE2 residues 19, 24, 27, 30, 31, 34, 35, 37, 38 and 42; conserved ACE2 residues F28, Y41, and L45. SARS2 contacting residues annotated within Logo plot. SARS2 N487 and Y489 residues in red also interact with ACE2-residue 83 (**See B**.). Logo plot of Aves/Reptilla is shown, which has a divergent amino terminus. Between Logo Plots reveals comparisons for SARS2-RBD, SARS1-RBD, RaTG13-RBD, and NL63-RBD predicted binding sites to human ACE2 (Residues: blue highlighted occur in ^3^/_4_ RBDs, red highlighted occur in ^4^/_4_ RBDs, purple highlight is NL63 specific). **B. ACE2 Nt Distal Region**. Predicted binding sites of human ACE2 for SARS2, SARS2, RaTG13 and NL63 (blue highlighted occur in ^3^/_4_ RBDs). SARS2 residue F486 interacts with L79 and M82 whereas, N487 and Y489 serve to coordinate ACE2 protein, Nt amino region and Y83. F486, N487, Y489 are all near disulfide Loop

Utilizing our co-plating in vitro assay and guided by the co-crystal studies of SARS2-RBD and ACE2, a set of eight mutations (Group I) targeting key residues in the Nt domain with either alanine or alteration in charge character **(Figure 8A)**. Another three mutations targeted multiple residues that would coordinate ACE2 Proximal and Distal regions by N487/Y489 residues. These multiple alanine substitutions at key ACE2 residues: Q24A, Y83A; T27A, F28A, K31A, Y83A; or L79A, M82A, Y83A (LMY➔A). None of the eight sets of point mutations had any effect on blocking ACE2 Nt contacts with SARS2-RBD. These results seem at odds with the co-crystal data, but not with the alanine-substituted SARS2-RBD mutants that also failed to disrupt binding and internalization.

**Figure 8:**
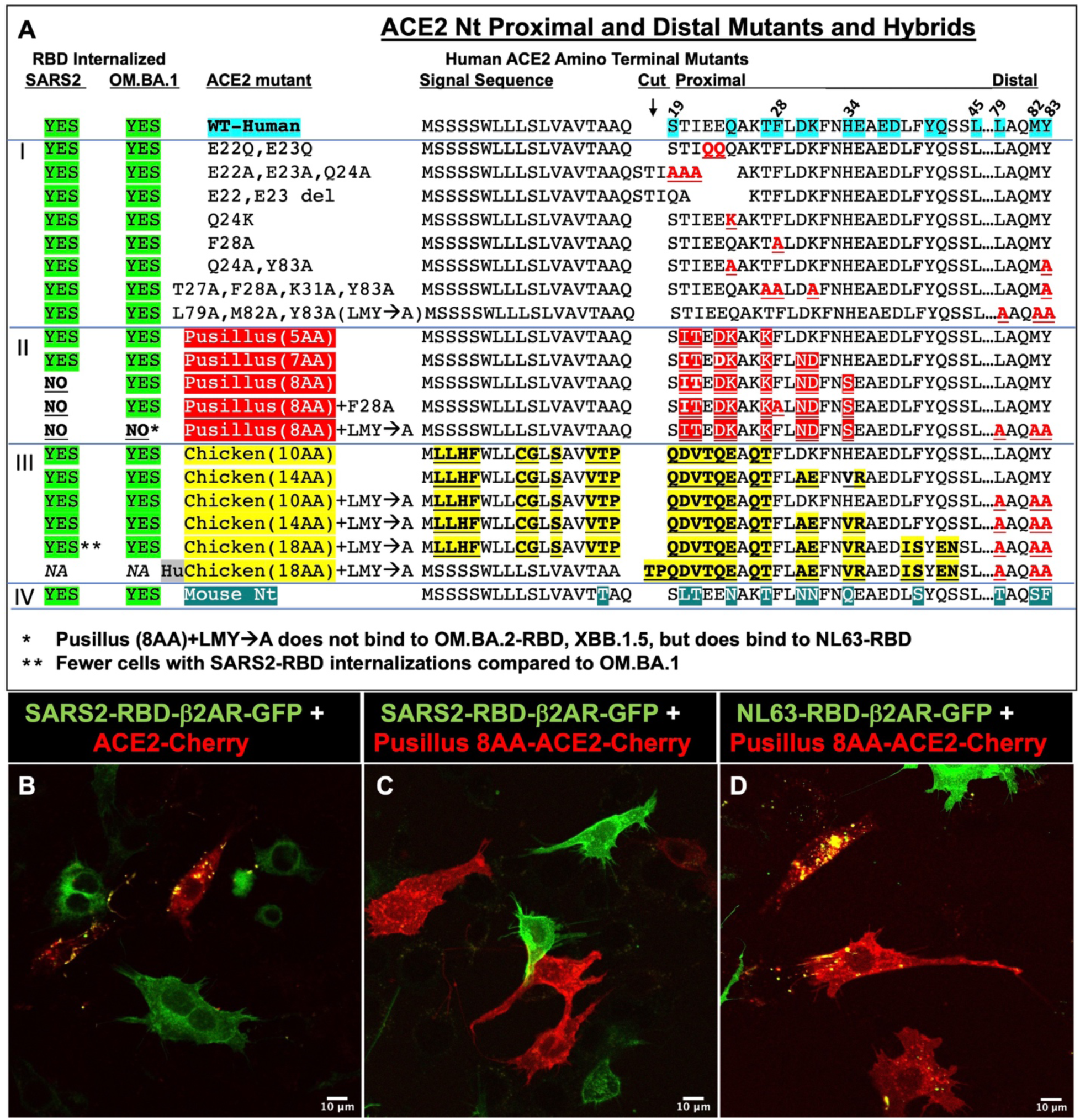
Amino Terminal human ACE2 Mutants. **A.** Twenty variants of human ACE2 (**Group I, II, III, IV**) were generated with amino-terminal mutations in the first 83 amino acids of the predicted protein. Only one mutant, HuChicken(18AA), did not show red fluorescence at the plasma membrane (PM). When RBD (green) internalization occurred, most of the red cells (ACE2 mutant) had internalized green fluorescence. To score for the absence of binding (as was performed for RBD mutants), at least three touching Red/Green cells had to be observed with no green-RBD internalizations (see Figure 3). **Group I mutants**: Based on crystallographic studies showing residues in the WT (wildtype) ACE2 (turquoise) that interact with SARS2-RBD. Mutations carrying T27A, F28A, K31A, Y83A is based on 4 residues that increase the affinity of RBD to ACE2 (25). No block of Internalization for green fluorescence for red fluorescence cells were observed in all eight variants for both SARS2 and OM.BA.1-RBD expressing cells. **Group II mutants**: Based on *R. pusillus* ACE2 not binding to SARS2-RBD (53, 54). Internalization of SARS2-RBD was observed for mutants containing the 8 amino acid substitutions, but they did not affect OM.BA.1-RBD internalization. Addition of F28A (a key residue in Y489 interaction) had no effect on OM.BA.1-RBD internalization. Only the addition of L79A, M82A and Y83A was able to abolish OM.BA.1-RBD internalization. **Group III mutants**: Based on Chicken ACE2 not binding to SARS2-RBD (53-55). In contrast *R. pusillus* mutants, none of these six mutants were able to abolish Internalization of green fluorescence in red fluorescence cells. It should be noted that NL63-RBD was internalized by both Pusillus 8AA and Pusillus 8AA+LMY➔A (**see Figure 9**). **Group IV mutant**: Nt Residues 1-94 are replaced with Mouse Nt residues, which have no effect on binding/internalization of SARS2-RBD or OM.BA.1-RBD. **B-D.** Utilizing LSM510, representative confocal images for co-plated RBD variants and ACE2-Cherry cells. **B.** Internalization of SARS2-RBD byACE2-Cherry cells, **C**. No internalization of SARS2-RBD by Pus8AA-ACE2-Cherry cells, **D**. NL63-RBD still shows internalization by Pus8AA-ACE2-Cherry cells.

It has been shown that ACE2 from three species *R. pusillus*, Chicken, and Turkey either poorly or do not bind SARS1 or SARS2-RBD domains (53-55). This lack of binding is believed to be a distributed consequence of residues along the entirety of the ACE2 protein. But sequence analysis showed that Aves/Reptilla (**Figure 7A**) were more variable than the entire set of 408 ACE2 aligned residues. This divergence set forth a hybrid ACE2 Nt approach substituting the first 43 residues: five mutants based on *R. pusillus* (Group II), and six mutants based on Chicken (Group III).

*R. pusillus* ACE2 has only nine differences with the human ACE2 protein between Q19 and L45, in the Nt proximal region where SARS2-RBD is predicted to bind. Three hybrids referred to as Pusillus (Pus) and number of amino acids (AA) were generated with different number of substitutions (5 amino acids)-Pus5AA, Pus7AA and Pus8AA and tested with SARS2-RBD. Neither the Pus5AA nor Pus7AA disrupted binding/internalization. But the Pus8AA completely disrupted/internalization **(Figures 8A-C, S11)**, which is consistent with the idea that Y489A and the Loop 4 region are important for SARS2-RBD binding to Nt ACE2. All variants and mutants of SARS2-RBD failed to bind or be internalized by Pus8AA: Delta, HiFrequency2020 3Ala (3 Alanine substitutions in the most common changes observed 2020), HiFrequency2021 7Ala (7Alanine substitutions in the most common changes observed in 2021), Predicted Binding Site (P.B.S.)-7Ala (7 Alanine substitutions from in silico prediction programs)(42), P.B.S. 6 Ala (one fewer substitution) as well as SARS1, and RaTG13 **(Figure S11)**. Surprisingly, RBD-OM.BA.1 and OM.BA.2 (**Figures 8A, 9A-F, S11)** were unaffected, as well as NL63-RBD (**Figure 8D**); all three RBDs were still bound and internalized by the Pus8AA substitution. This might have been expected for NL63 due to its paucity of contacts with the ACE2 Nt proximal region (**Figure 7A)**.

F28 is strongly conserved in ACE2 and is predicted to interact with the critical SARS2 residue Y489, but an F28A mutation did not block SARS2-RBD internalization. To extend these observations, F28A was added to the ACE2 Pus8AA hybrid to determine if OM.BA.1 or BA.2 would be blocked from binding/internalization, but no decrease in binding was observed (**Figure 8A)**. These observations conclude that there is a differential reliance on the ACE Nt proximal region for SARS2 versus OM.BA.1, OM.BA.2 and NL63-RBDs, suggesting multiple modes of ACE2 binding/internalization. Subsequent RBD variants OM.BA.4/BA.5 (both with L452R, F486V and R493Q relative to OM.BA.2) do not bind to the ACE2 Pus8AA Nt mutant, suggesting this newer variant had interactions more like that of SARS2-RBD. Finally, data from ACE2 deep mutational screening (DMS) (25, 41) show that SARS2-RBD-H34S does not block interactions, yet the wild type H34 residue in ACE2 Pus7AA hybrid did rescue SARS2-RBD binding. Taken together Pusillus AA hybrids might change conformation of the Nt region or disrupt specific residue interactions between SARS-RBD and ACE2. AlphaFold analysis of ACE2 Pus8AA hybrid did not reveal change to the **helix** (residues 20-52)-**turn**-**helix** (residues 55-81) structure of the Nt proximal and distal regions, respectively (**Figure S12**). However, the amino terminus did adopt novel conformations that perhaps disrupt its contacts with the RBD Loop 4 region. By contrast Pus7AA maintains the H34 residue, which is predicted to interact with SARS2 residues Q493, L455, and Y453. But this interaction is unlikely to be critical as chicken14AA containing H34V does not block binding nor does H34S (25). Taken together, the Pus8AA hybrid likely changes the conformation and contacts of the Nt proximal region, leading to a destabilized interaction between SARS-RBD and ACE2.

My proposed model of disulfide Loop 4 interactions suggested that it coordinates its binding between the ACE2 Nt proximal and distal Regions (ACE2 residues 79-83). Based on the ACE2 Pus8AA mutation, SARS2 may primarily utilize the ACE2 Nt proximal region and conversely, OM.BA.1 and OM.BA.2 RBDs might primarily utilize the Nt distal region. None of the Group I mutations analyzed, including the Nt distal region mutant LMY➔A altered OM.BA.1 binding/internalization **(Figure 8A)**. The RBD domains of OM.BA.1 and OM.BA.2 contain 16 mutations that differ from SARS2, which perhaps caused them to rely on both the Nt proximal and distal regions. To test this idea, the ACE2 Pus8AA hybrid was combined with the LMY➔A mutant. Pus8AA + LMY➔A was finally able to block OM.BA.1 and OM.BA.2 RBDs from being bound and internalized (**Figures 8A, 9G-I, S11**), confirming the hypothesis; the NL63-RBD, which does not bind to the ACE2 Nt distal regions, was unaffected as expected (**Figure 9J-L**). These Group II ACE2 mutants show that SARS2, OM.BA.1 and NL63 RBDs have differential contact points to promote their binding/internalization

**Figure 9:**
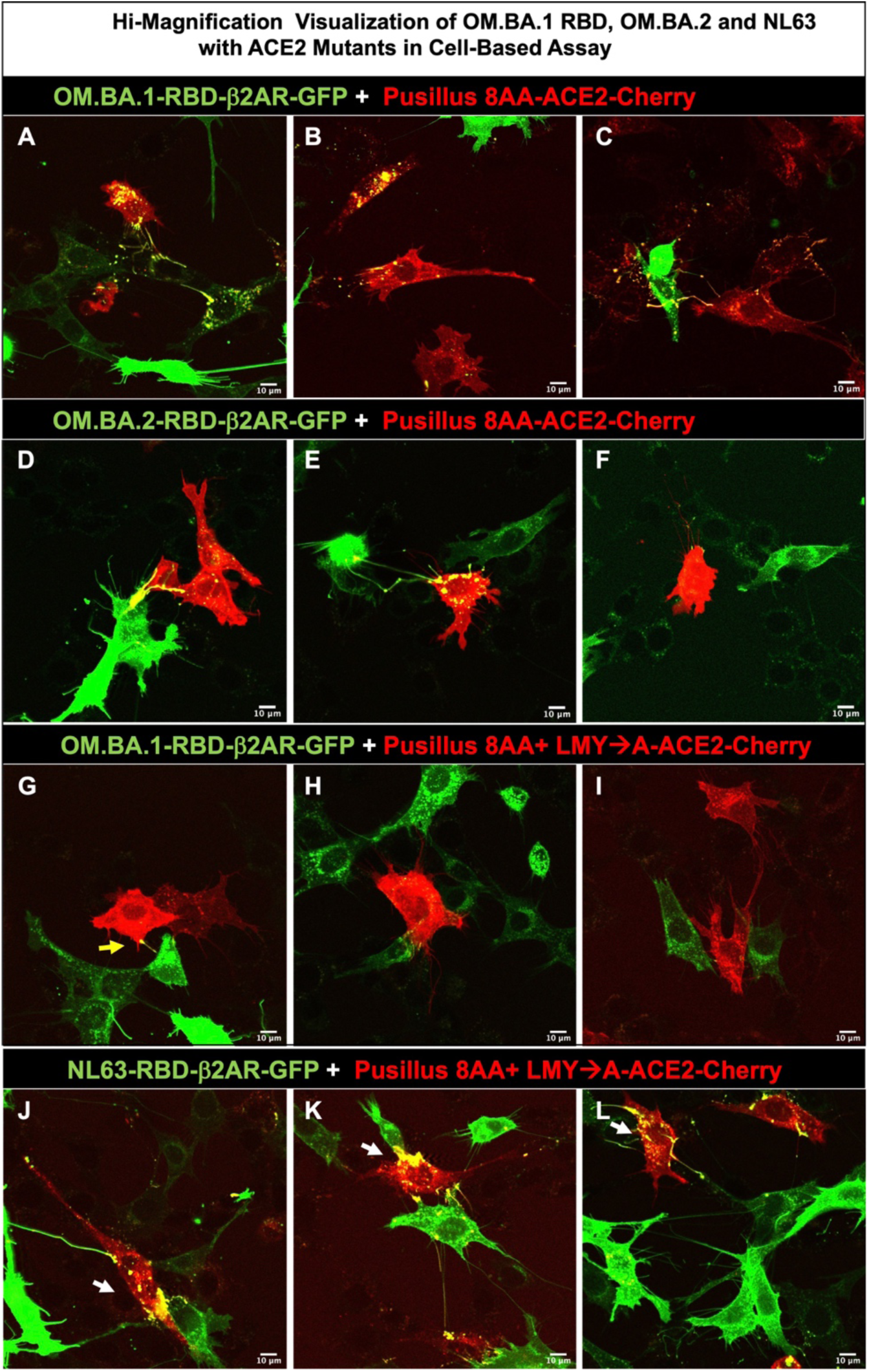
Visualization of OMBA.1 RBD mutants and NL63-RBD in Cell-Based assay with ACE2 mutants.-I. Utilizing LSM510, confocal images for co-plated RBD variants and mutants with ACE2-Cherry cells. **A-C**. Three representative images showing OM.BA.1-RBD protein readily internalized into Pus8AA-ACE2 expressing cells. **D-F**. Three representative images showing OM.BA.2-RBD protein readily internalized into Pus8AA-ACE2 expressing cells. **G-I.** Three representative images showing OM.BA.1-RBD protein **not** internalized by Pus8AA+LMY➔A-ACE2-expressing cells. **J-L.** In contrast, three representative images showing NL63-RBD protein readily internalized by Pus8AA+LMY➔A-ACE2-expressing cells.

In contrast to the Group II hybrids, Chicken ACE2 mutants (Group III) containing both the Nt proximal and distal region substitutions did not block SARS2, OMBA.2 nor NL63 RBDs from binding and internalization. Group III data suggest that some combination ACE2 Pus8AA residues can block SARS2-RBD binding by a robust change in contact residues (**Figures 7A, 7B, 8A**). The totality of data for OM.BA.1 and OM.BA.2 RBD binding point to a conformational shift in their RBDs that alters their usage for ACE2 Nt Proximal and Distal regions. Consistent with this conclusion, both Y489A and P491A are properly trafficked to the plasma membrane for the SARS2-RBD, whereas analogous mutations in OM.BA1-RBD fail to traffic to the plasma membrane likely due to a change in conformation leading to failure in the formation of the correct cysteine bonds (56).

Mice cannot be infected by SARS2, which is why mutant mice strains have been generated with the human ACE2 CDS replacing the mouse ACE2 gene (57). A third hybrid (Group IV) with the mouse (m)ACE Nt proximal and distal was made with 25 of the first 94 residues altered (**Figures 8A, S13**). This Group IV hybrid contained mutations at similar residues to the Pus8AA as well as a LMY➔TSF mutation. Yet, no alteration in binding or internalization was observed for SARS2, OM.BA.1 and NL63 RBDs. Thus, substantial changes in the ACE2 Nt proximal and distal regions suggest a lack of specificity for RBD binding. Interactions were quantified for four ACE2 substitutions by our Dissociation-Plate-Flow Cytometry *(DPF) assay*: Pus8AA, Pus8AA +LMY➔A, and mACE2 Nt 1-94 with RBDs from SARS2, OM.BA.1, NL63 and a newer variant XBB.1.5 (**Figure 10A-A’**).

**Figure 10:**
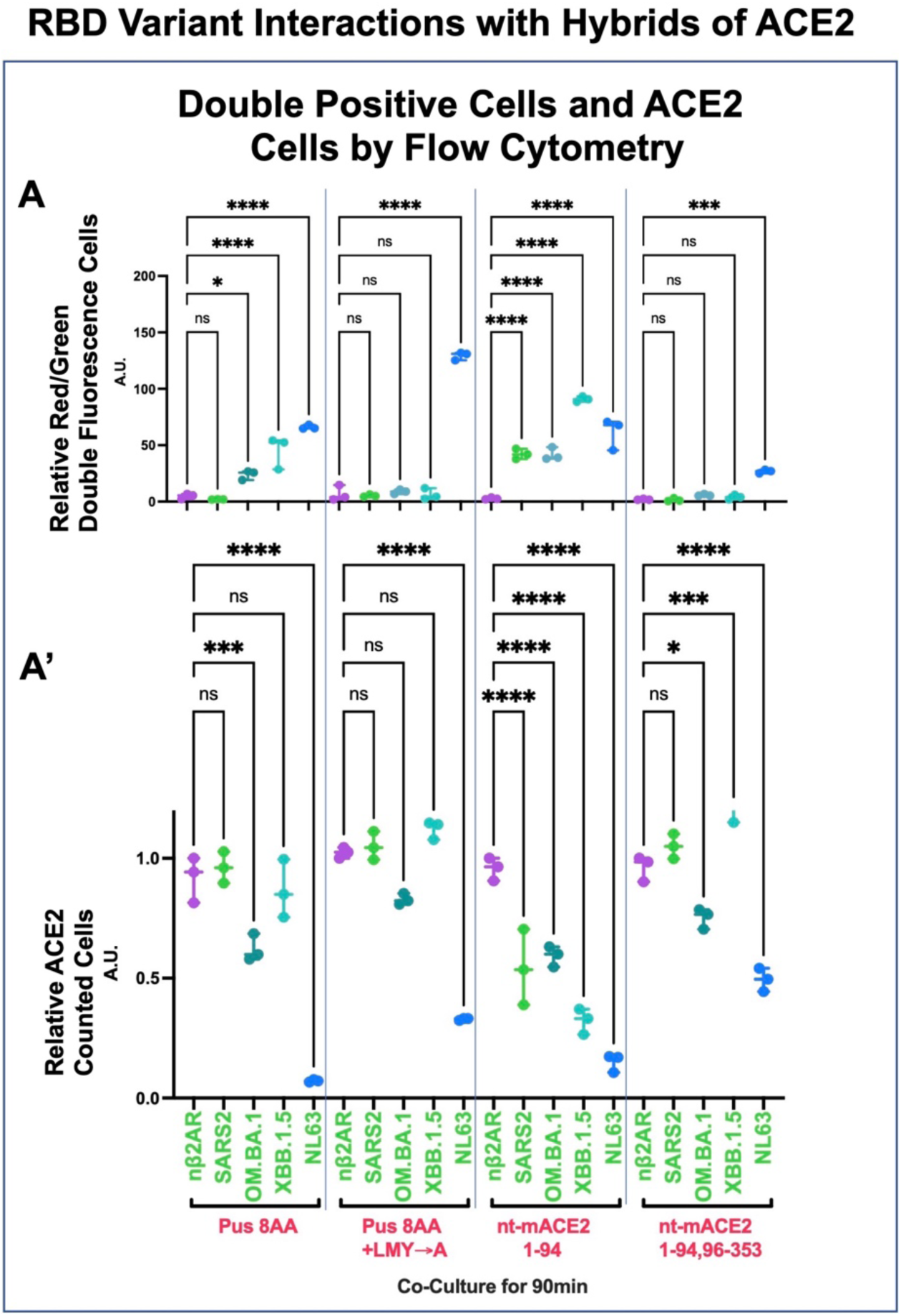
RBD Variants interactions with Mutated ACE2. **A.** Results from visual inspection for Pus8AA, Pus8AA+LMY➔A and Nt-Mouse residues 1-94 (Figure 8) were further quantified by the *DPF assay* for nβ2AR, SARS2-RBD, OM.BA.1-RBD, XBB.1.5-RBD and NL63-RBD. In addition, a near complete swap of human residues 1-353 with equivalent region of mouse residues. NL63-RBD show double fluorescence in all four ACE2 mutants and a reduction of ACE2 fluorescence, revealing all ACE2 constructs have the capacity to function. SARS2 variants show differences (**see Figure 9 and Data File S2**). Values in **A and A’** are normalized to controls (Y-axis arbitrary units, A. U.). *For Pus8AA*: only SARS2-RBD shows no binding and hence no double fluorescence (1.92±0.17). *For Pus8AA+LMY➔A*: SARS2-RBD, OM.BA.1-RBD, show no binding (5.23±0.99) and (8.75±1.94), respectively. *For Nt-Mouse residues 1-94*: All variants show binding/double fluorescence (58.22±28.20). *For Nt-Mouse 1-94,96-353:* All variants show **no** binding/double fluorescence (3.71±2.14). **A’**. The ACE2 levels showed variability for XBB.1.5-RBD, as it does not downregulate Pus8AA (a loss of 13.3%±12.1%) as was previously shown for ACE2 but does downregulate the Nt-Mouse residues 1-94 ACE2 chimera (a loss of 67.8%±5.3%). To date, XBB.1.5 is the only variant where binding **can be** decorrelated from an ACE2 decrease in level of expression. GraphPad Prism 9.3.1 used to generate **A and A’**.

Indeed, flow cytometry quantification of double positives for SARS2, OM.BA.1, NL63 mimicked the co-plating assay results with XBB.1.5 showing binding and internalization like that of OM.BA.1 in all four Nt ACE2 hybrids (**Figure 10A, Supplemental Data File S2**). In contrast to OM.BA.1, XBB.1.5 did not cause a change in fluorescence to Pus8AA or Pus8AA +LMY➔A-Cherry cells. OM.BA.1, despite a lack of robust binding to Pus8AA +LMY➔A (8.75±1.94), still showed modest reduction to its fluorescence (17.2%±2.2%) (**Figure 10A’**). Finally, a fourth mutant mACE 1-357 was tested, which abolished binding for RBDs SARS2 (1.45±1.37), OM.BA.1 (5.71±0.89), and XBB.1.5 (3.98±2.08), but not for NL63 (26.7±1.60) (**Figure 10A**). An interaction could be observed for OM.BA.1 by the 24.8%±0.04% loss of ACE2 fluorescence (**Figure 10A’**). These data are consistent with NL63 having a greater set of contacts with residues 321-330 of ACE2 (**Figure 11A**).

**Figure 11:**
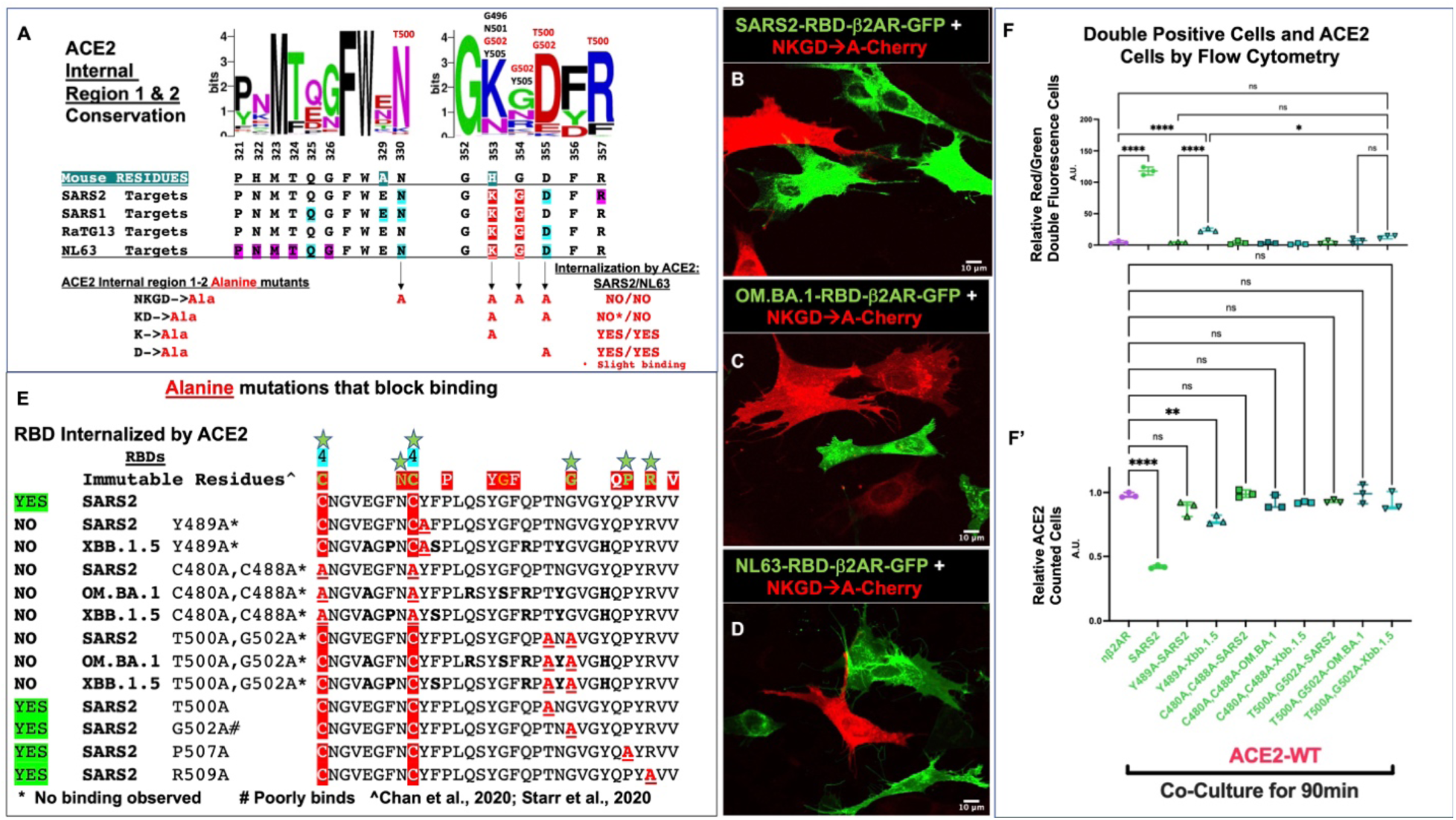
Internal human ACE2 mutants and RBD interaction mutants. **A.** ACE2 Internal Region. Predicted binding sites of human ACE2 for SARS2, SARS2, RaTG13 and NL63 (Residues: blue highlighted occur in ^3^/_4_ RBDs, red highlighted occur in ^4^/_4_ RBDs, purple highlight is SARS2 specific). ACE2 mutants tested based on conserved predicted binding sites: 1) N330A, K353A, G354A, D355A (NKGD➔A); 2) K353A, D355A (KD➔A); 3) K353A; 4) D355A. Both SARS2 and NL63 RBDs were tested in two ways: cotransfection or separate transfections followed by overnight co-plating (**see Supplemental Figure S14**); Co-transfection of RBD-β2AR-GFP and ACE2-Cherry, which could interact, typically internalized membrane distribution of both proteins. Mutant NKGD➔A, which poorly traffics to the plasma membrane, shows no binding to either SARS2 or NL63 RBDs, KD➔A shows no binding to NL63 RBD, but some binding to SARS2-RBD without internalization. By contrast, the single K353A and D355A mutants both revealed binding and internalization of both NL63 and SARS2 RBDs. **B-D.** Utilizing LSM510, representative confocal images for co-plated RBD variants and mutated NKGD➔A ACE2-Cherry cells. Neither **B.** SARS2-RBD, **C**. OM.BA.1-RBD, nor **D**. NL63-RBD show internalization into mutant ACE2-Cherry cells. **E.** Alignment of different SARS2/OM.BA.1/XBB.1.5 mutants and that were tested for block binding (**see Supplemental Figure S13**); the region consists of residues 480-511 from SARS2. Bolded residues differ from SARS2. Bolded Red and underlined residues are alanine mutations. Mutants tested were Y489A and the double mutant C480A, C488A on SARS2, OM.BA.1, XBB.1.5 that previously showed no binding. In addition, the T500A, G502A double mutant, which is predicted to bind to KGD region of human ACE2. Visual observations were noted and quantified. **F and F’.** *DPF assay* quantification for mutants that did not predict binding by visual inspection. Only XBB.1.5-Y489A-RBD showed slight binding as some double positive Red/Green cells could be observed (23.67±2.87) and some loss of ACE2-Cherry fluorescence is shown (21.7%±3.4%). Values in **F and F’** are normalized to controls (Y-axis arbitrary units, A. U.), **see Data File S2**. GraphPad Prism 9.3.1 used to generate **F and F’**.

#### Co-transfection of membrane tethered RBDs and ACE2

In vitro co-plating, while simple, requires visual inspection of RBD-expressing cell touching an ACE2-expressing cell. This assay is robust but can be difficult to quantify weak binding without internalization. The *DPF assay* is very quantitative, but more labor-intensive. To more quickly rule out weak interactions, a co-transfection and plating assay was set up to see if protein interactions could be observed within the same cells, as is reminiscent of the interactions published decades ago for other cell surface receptors and ligands, the SEVENLESS-BOSS (58) proteins as well as the NOTCH-DELTA (59) proteins. A priori, it was not known if membrane-bound RBD and ACE2 proteins could interact if co-expressed. Here, it was expected that interactions between expressed proteins would be at a higher probability compared to those between cells. Co-transfection between nβ2AR-GFP (n=signal peptide) with ACE2-Cherry in the same cells and imaged after 16 hrs reveals robust membrane trafficking for both proteins in the same cells (**Figures S14A-A’’**). By contrast, co-transfection of SARS2-RBD with ACE2-Cherry shows only robust SARS2-RBD at the membrane and filopodia with ACE2-Cherry internalized (**Figures S14B-B’’)**. Internalization of ACE2 by RBD through intra- or intercellular interactions may be clathrin dependent (40);. This final piece of evidence firmly concludes that ACE2-Cherry is subordinate to the effects of the SARS2-RBD protein and does not cause cell death. This co-transfection assay confirms and extend the results observed on ACE2-Cherry membrane localization: Y489A (**Figure S14C-C’’**) and P.B.S 7Ala (**Figure S14D-D’’**) do not affect ACE2 internalization, C480A, C488A (**Figures S14E-E’’**) had little effect; and P.B.S. 6Ala (**Figures S14G-G’’**) did not block internalization.

Consistent with co-plating experiments, co-transfection of SARS2 and OM.BA.5 RBDs with Pus8AA hybrid ACE2 did not show binding/internalization (**Figures S15A-A’’, S15B-B’’**) whereas co-transfection with OM.BA.1 RBD did show internalization (**Figures S15C-C’’**). Surprisingly, OM.BA.1 RBD did internalize Pus8AA +LMY➔A (**Figures S15D-D’’**), despite it not forming double fluorescent cells by *DPF assay* (**Figure 10**), but perhaps consistent with the loss of fluorescence (**Figure 10**). Co-transfection proved an effective way for simultaneously assessing the membrane trafficking of ACE2 and RBD mutants as well as determining if proteins could interact. It is likely that co-transfection reveals less stable contacts between RBD and ACE2, which still allow for the internalization of the ACE2-Cherry protein (compare **Figures S14E-E’** to co-plating in and DPF assay results in **Figures 11F-F’**). It should be noted that co-transfection results likely mimics SARS2 viral production within ACE2-expressing cells.

#### ACE2 Internal region mutagenesis

Interpretation of the data thus far suggests the ACE2 Nt proximal region is necessary for SARS2-RBD binding/internalization and both ACE2 Nt proximal and distal regions are necessary for OM.BA.1 and BA.2-RBDs; Neither region is necessary for NL63-RBD binding/internalization. SARS2-, SARS1-, RaTG13- and NL63-ACE2 crystal structures map residues to two additional Internal regions of ACE2: residues 321-326 (PNMTQG); 330 (N) and 353-356 (KGDF), with NL63 contacting all these residues, with SARS2, SARS1 and RaTG13 contacting different subsets (**Figure 11A**). All four of these RBDs contact ACE2 conserved residues K353 and G354 (**Figure 11A**). G354 contact is not likely as important as the D355 contacts found in three RBDs (*not* SARS1). NL63-RBD being unaffected by the Pus8AA+LMY➔A mutant nor mACE2 1-357 suggests that this Internal region is key to triggering binding and internalization of all RBDs. It was previously shown that ACE2 mutants K353A and D353A nearly abolish all SARS and NL63 infections (48, 60, 61). Several alanine substitutions tested this region for interfering with SARS2 or NL63 binding/internalization: quadruple mutant-N330A, K353A, G354A, D355A, the double mutant-K353A and D355A, and individual mutations in K353A and D355A (**Figure 11A**). The NKGD➔A and KD➔A mutants blocked both SARS2 and NL63-RBD binding and internalization (**Figures 11A-D, S15E-E’’, S15F-F’’**). But the individual K353A and D355A mutations had no effect (**Figure 11A**). The internal region mutants point to a common binding site on ACE2 that all RBDs utilize for binding/internalization.

Crystal studies of RBD with ACE2 suggest K353-G354-D355 in ACE2 interacts predominantly with T500 and G502 in the SARS2-RBD, a region distinct from the Loop 4 region (1, 28). G502, P507, and R509 all represented immutable residues from DMS studies suggesting they could all be key residues for ACE2 interaction. Co-transfections were carried out for this region of SARS2-RBD: T500A, G502A, P507A, P509A and T500A, G502A double mutant (**Figures 11E, S14H’-H’’ through S14L-L’’**). Only the double mutant T500A, G502A was able to completely abolish ACE2 binding (**Figures S14J-J’’**). These data argue that two regions of SARS2-RBD work together to coordinate binding with ACE2: T500 and G502 coordinate the internal region of ACE2 while Y489 and the Loop 4 disulfide bond (C480-C488) coordinate the Nt distal and proximal residues. Perhaps the effect of these mutations is not universal across variants as the Y489A mutation in OM.BA.1 disrupted membrane trafficking, likely due to a disruption in RBD folding (**Figure 11E**). The *DPF assay* was once again utilized to compare and quantify these three types of mutations in the SARS2, XBB.1.5 and OM.BA.1 RBDs (**Figures 11F, F’, S14E-E’, S14F-F’**); Both C480A, C488A (3.04±1.04) and T500A, G502A (7.87±4.80) disrupted binding/internalization (double positives) of ACE2 across all variants (**Figures 11E, 11F-F’**). The sum of all RBD mutations produced suggest binding is localized to a stretch of residues between Y489-G502, which is also where blocking antibodies target (28).

#### Can RBD binding to ACE2 be antagonized by KGD containing peptides?

A common binding site on ACE2 for both NL63 and SARS2-RBDs suggested that a competitive antagonist based on the Internal region 2 amino acid sequence could be derived. Six peptides were generated and tested for functional antagonism in the solution binding assay at micromolar concentrations (**Figure S16A**). None of the peptides interfered with RBD-ACE2 binding for both NL63 and SARS2-RBDs. However, one of the peptides (GKGD) caused red fluorescence of ACE2-Cherry cells to surround SARS2-RBD expressing cells (**Figure S16A**). Because the tripeptide Lys-Gly-Asp (KGD) is like the high affinity Arg-Gly-Asp (RGD) tripeptide for fibronectin (49), two RGD containing peptides for blocking activity were also tested at micromolar concentrations, but neither had any impact. Next, the GKGD peptide alone was tested in the *DPF assay* with a preincubation of 100μM GKGD peptide, followed by flow cytometry analysis for both SARS2 and NL63-RBDs (**Figures S16B-B’**); Again, no mitigation of SARS2-ACE2 binding was observed. Finally, purified His-tagged SARS2-RBD (60ng, ∼250 residues) was added to ACE2 cells with or without preincubation with 49μg of the 4-residue GKDG peptide in the *DPF assay*; Despite a ∼50,000x molar excess of peptide, no decrease in RBD binding was observed (**Figures S16C-C’’**). Higher than 100μM peptide concentrations were not tested due to rampant nonspecific effects at mM presentation of compounds. Taken together, these data suggest that small KGD-containing peptides do not bind RBDs with sufficient affinity to block ACE2 binding.

## Discussion

### RBD Mutants

#### Spike S1 subunit-RBD interactions with ACE2 are critical for cellular internalization

Respiratory coronaviruses like SARS1, SARS2 and NL63 utilize their spike protein to bind to ACE2 on the surface of cells that line the respiratory epithelium(44, 60). Mapping studies from SARS1 identified the minimum protein sequence necessary to bind to ACE2, a 193 amino acid region-the receptor binding domain (RBD), which is also found in SARS2 (1, 23, 41). Binding studies with SARS1 and SARS2 RBDs show roughly the same high affinity binding to ACE2, which does not correlate to viral transmission rates (62, 63). One of the key differences in viral transmission rates is the addition of 12 nucleotides within the Spike protein, creating a four-amino acid RRAR-furin cleavage target after residue 684, allowing for pre-activation(64, 65), splitting the protein into an Nt-S1 and Ct-S2 subunits (66). The liberation of residues 1-684 (S1), containing the RBD from 333-529, likely provides easier access for binding to ACE2 and for TMPRSS2 to cleave at the S2 subunit and reveal the S2’ Fusion Peptide (67). It is thought that the RBD association with ACE2 allows proximity access to the plasma membrane located TMPRSS2; The S2’ Fusion Peptide subsequently drives membrane fusion events for SARS1, SARS2 and NL63. The work here suggests that the RBD association with ACE2 may also take part in the viral transmission and/or internalization process. In this regard the natural propensity of ACE2 to be internalize makes it an attractive target for coronaviruses but is unlikely the only target as RpYN06 which does not bind ACE2, must be internalized by a distinct membrane target.

#### Binding efficiencies and homologies between RBD across species

There are 9 betacoronavirus and 1 alphacoronavirus with measured Kd of binding between RBD and human ACE2. SARS1 and SARS2 have been measured at 58nM and 16nM respectively(62). High affinity is a trademark of most of these coronaviruses except for RaTG13 and RsYN04, which bind in the micromolar range (26, 45). The only alpha coronavirus that binds to ACE2, does so at 17nM (48), nearly identical to that of the SARS2 RBD. H-CoV-NL63 is a common human “cold virus” and has not caused a worldwide pandemic, which again argues that affinity by itself does not correlate with virulence (68). SARS1& SARS2-RBDs completely lack substantial NL63-RBD amino acid homology or common tertiary structures but co-crystal studies with ACE2 reveal similar contact sites within its Nt and Internal regions (27). This convergence on ACE2 contacts suggests that only certain regions can provide high affinity interactions and/or mediate its internalization, both of which may be critical to viral transmission.

#### An in vitro RBD-ACE2 binding assay

It has recently been shown that SARS2 infection of the olfactory system is a critical step for infection to prosper (69). These results argue that propagation of the preliminary infection allows for both asymptomatic dissemination of the SARS2 virus as well as symptomatic consequences of deeper respiratory infections within ACE2/TMPRSS2 expressing cells that line the respiratory tract. If a compound can be identified to block the SARS2-Spike S1 subunit binding to ACE2 in the olfactory epithelium, then it stands to reason viral transmission rates could be reduced. The in vitro system set up here robustly traffics RBD domains and ACE2 to the plasma membrane. The filopodia that exhibit RBD-β2AR-GFP fluorescence are 1μm or less, which is like the average size of a betacoronavirus particle, 0.1 μm (70-72). Thus, filopodial membrane interactions with the plasma membrane (PM) of the ACE2-Cherry expressing cells may show similarities to interactions between viruses that infect new cells. A 1000-fold range of RBDs with published affinities were tested, yet no differences were observed in the assay, consistent with pseudotype virus assays (53-55). Overall, the in vitro assay may represent a higher throughput strategy to study the molecular details of RBD binding to ACE2.

#### ACE2-Cherry protein relocalizes upon binding to RBD-β2AR

The RBD portion of spike proteins function by bringing its respective coronavirus to the PM where enzymes like TMPRSS2 cleave the spike protein liberating a S2 Fusion Peptide whose function is to fuse viral and cellular membranes and enable endosome-independent entry (66). One of the surprising findings here is that ACE2 by itself promotes internalization of RBD tethered to β2AR. In addition, rapid loss of ACE2-Cherry fluorescence at the PM (<10min) without formation of red/green doublet cells (**Figures 2D’**) suggests that reversable RBD binding is sufficient to trigger ACE2-Cherry loss. The mechanism of ACE2-Cherry loss is not known, but targeted degradation seems likely as other membrane localized proteins are not being lost (**Figures 3, S4**). It has been published that the SARS2 S protein also causes the loss of ACE2 expression in HEK-293A cells over the course of several hours and that this process can be reduced by the lysosomal inhibitor BafA1. Here I show membrane levels of ACE2-Cherry in OP6 cells over the 90minute interaction period with SARS2-RBD are still reduced (**Figure 3F’**), but the number of double positive cells did increase suggesting that ACE2 proteins were perhaps more stabilized at the membrane (**Figure 3F**). In this context, NL63-RBD double positives also increased, but just below statistical significance (**Data File S2**).

#### Meta-Analysis and mutational analysis of RBD reveal critical residues around disulfide Loop 4

The realization of a functional RBD-ACE2 interaction assay led to a meta-analysis of RBD mutations that were either derived from natural selection during infection (Variants) or DMS analysis by two laboratories (25, 41). Variants typically contain two or more mutations. By contrast DMS mutations are single substitutions of all 19 amino acids at each of the 197 residues of RBD, leading to an analysis of nearly 4000 mutants. DMS shows the distributed nature of RBD binding to ACE2. The mapped SARS2-RBM with ACE2 constitutes a 74 amino acid region of which DMS meta-analysis suggests 7 of these residues (R454, C480, N487, C488, G502, P507, R509) to be necessary for binding to ACE2. Six of these residues are predicted to have critical roles in the RBD double beta barrel configuration: Positively charged residues R454 and R509 likely connect to D667 and D442, respectively and neither are predicted to bind ACE2; residues C480 and C488 form the Loop 4 disulfide bond; and finally alpha helical breaking residues G502 and P507 that likely promote formation of the R509-D442 bridge. This leaves only N487 that would be necessary for binding and indeed is predicted to make contacts with both ACE2 Nt proximal and distal residues Q24 and Y83, respectively. But the N487A mutation in RBD and both Q24A and Y83A mutations in ACE2 also did not affect binding or internalization. Thus, none of these 7 residues would appear to confer the specificity of RBD-ACE2 binding.

Two additional analyses were performed looking at the most abundant mutations at multiple positions in the RBD in years 2020 (HiF2020-3 residues) and 2021 (HiF2021-7residues). Alanine mutants at all seven positions K417, N439, L452, S477, T478, E484, and N501 had no effect on the binding or internalization of ACE2. A second set of seven alanine substitutions of in silico predicted binding sites (P.B.S.): K417A, G446A, L455A, F486A, Q493A, Q498A, N501A. This P.B.S. 7Ala-RBD mutant did block binding and internalization. Two of these residues, L455 and F486 are not mutated in OM.BA.1, suggesting that one or both may be critical for binding. Reversion of residue 486 to phenylalanine in the P.B.S. 6Ala-RBD mutant did indeed rescue RBD binding and internalization by ACE2. But the F486A mutation alone in RBD and corresponding mutations in ACE2 at M82A and Y83A did not abolish binding and internalization. Thus, the totality of these alanine mutations, like the meta-analysis of the DMS, point to a distributed binding between RBD and ACE2.

A distributed nature of RBD binding seems antithetical to Kds with nanomolar affinities. DMS analyses of binding max studies reveal several residues G447, N448, Y449, N487, and Y489 that are critical for binding with Y489 being the most critical (43), yet Y449A, N487A did not affect binding and internalization; Y489A did prove critical. Juxtaposed to Y489 is the Loop 4 disulfide bond C480-C488 that is absent in the RBD RpYN06 and does not bind to ACE2(45). A logical conclusion is that Loop 4 may be critically important for RBD binding or “attachment” to the Nt of ACE2. All Loop 4 residues between C480 to C488 were substituted with SARS1 and RaTG13 as well as single alanine substitutions between F486 and L492. From all these swaps it did not appear that any one residue was key to the functionality of Loop 4, but rather supported the idea that formation of Loop 4 itself is critical. By contrast, blocking the formation of Loop 4 with the double alanine substitution of C480 and C488 prevented internalization of RBD by ACE2-expressing cells. Despite this result, it is not possible for Y489 alone to confer non-covalent high affinity binding to ACE2. Perhaps the disulfide Loop 4 is an anchor onto the Nt of ACE2 that was not observed by the crystallographic studies (1). Finally, DMS mutations (25, 41) would have predicted that N487A to be as debilitating as Y489A, which was not observed. Overall, the mutagenesis and meta-analysis failed to identify a contiguous stretch of a few residues that would bind ACE2.

The alphacoronavirus NL63 RBD has a completely different primary sequence and folding compared to RBDs of RaTG13, SARS1, and SARS2 as well as its RBM region is split over three surfaces, yet all binds similar regions on ACE2. But the amino acid identity is limited to just eleven residues with four of them surrounding the SARS2 Loop 4 region: C488, F490, P491 and L492 (C577, F579, P580, L581 in NL63); This region of NL63 RBD is not implicated in any RBM, but a N578Y mutation a generates a five amino acid match to SARS2 from C488-L492 and improves the NL63 RBD binding Kd 2-fold. In addition, the NL63 C577 residue is part of disulfide Loop 3 (**See Figure S6**). Taken together, the residues surrounding the disulfide Loop 4 in SARS2 is likely one of the critical regions for RBD interaction with ACE2. It is also possible that Loop 4 (Loop 3 in NL63) serves no “binding” but rather as a hook onto the ACE2 amino terminus where it interacts. As such, neither SARS2-Y489W-RBD nor SARS2-Y489H-RBD mutations disrupted binding and internalization with ACE2 despite Histidine’s remarkably different properties as a charged polar group. These results suggest that the effect of Y489A is not a change in the direct interaction between it and ACE2, but perhaps a more subtle change to the Loop 4 positioning relative to the ACE2 amino terminus. The Y489A-OM.BA.1-RBD mutation disrupts PM trafficking, which may result from a failure to form the disulfide bridge C480-C488 like results for C488A-RBD.

#### Could odors act to mitigate SARS-CoV-2 binding?

The initial drive to study RBD/ACE2 interactions was to identify an odor that could mitigate infections. But none of the 36 odor mixtures from the “Le Nez du Café” were able to mitigate formation of RBD/ACE2 double positive cells nor reduction of ACE2 expression levels. A more directed approach could focus on the complete set of 4000 “relevant” odors for humans (https://ifrafragrance.org/priorities/ingredients/ifra-transparency-list) or the 230 Key Food Odors, and 800 common human odors (http://www.flavornet.org) that exist in everyday products. A recent study suggested that Honeysuckle extract might mitigate infection (73).

A second approach would be to identify critical di or tri-peptide motifs for antagonism within the 197 residue RBD. But RBD mutagenesis did produce one tangible target: the disulfide bridge between C480 and C488. It is shown here and at other laboratories that DTT and a more pH stable version DTBA can mitigate RBD/ACE2 binding, albeit at high concentrations. Both compounds are volatile and perceived as having a foul smell due to their thiol groups. These odors could be masked by sweet-smelling odors. Additionally, there may be other odors that are more volatile and capable of disrupting the disulfide linkage, which could represent another solution to reduce rates of infections by an inhalation regimen. Like many chemotherapies, a reducing agent lacks specificity and would damage other proteins but could have value in short-term decreases in the viral load. Paxlovid is very good at reducing the SARS2 viral load despite many unwanted side effects (74).

Vaccines provide IgG responses that can limit viral amplifications, but unfortunately, mucosal protection (IgA) in the nasal epithelium wanes more quickly (75). Thus, most individuals are not immune to infection and the subsequent viral amplification within the sustentacular cells that carry ACE2 and TMPRSS2 expression. If inhibitors can block this initial viral insult (7), then extent of disease would likely be mitigated as well.

### ACE2 Mutants

#### The disulfide Loop 4 region interacts with Nt proximal and distal regions of ACE2

The importance of disulfide Loop 4 located residues F486, N487 and Y489 from RBD analysis, would seem to correlate with the Nt region in ACE2 as critical for binding. RBD residues 486-489 bridge Nt proximal region residues Q24, T27, F28 with Nt distal region residues L79, M82, Y83 (LMY) (**See Figure 7B**). But ACE2 mutations (Group I) targeting these residues did not affect RBD binding and internalization. Group II Nt Hybrids with *R. pusillus* sequences abolished SARS2-RBD interactions, but not OM.BA.1. Combining Nt proximal *R. pusillus* residues and Nt distal substitutions LMY➔A did lead to a complete block in binding and internalization by OM.BA.1-RBD. These results suggest that the 15 amino acid differences in OM.BA.1-RBD lead to a new protein conformation, which is consistent with the absence of Y489A protein trafficking in a OM.BA.1-RBD. At first approximation, these data argue that Nt proximal and distal ACE2 regions are necessary for binding and internalization of SARS2 and OM.BA.1-RBDs. But the Nt mACE 1-94 hybrid still providing binding and internalization, suggest a lack of specificity.

#### SARS-CoV-2 and H-CoV-NL63 both require common Internal ACE2 region for binding

The NL63-RBD binding and internalization by ACE2 was unaffected by any of the 20 Group I/II/III/IV Nt proximal or distal mutations. Those results are consistent with the co-crystals showing only a few contacts by NL63-RBD in the Nt proximal and distal domains. By contrast, NL63-RBD has extensive contacts in both Internal regions 1 and 2 whereas SARS2-RBD has more of its contacts to Internal region 2. Previously, it was shown that K353A had substantial impact on both NL63 and SARS2-RBD binding in pseudovirus infections but did not abolish it. Our co-plating and co-transfection assays also did not reveal complete blocks in binding and internalization. While the single substitution D355A did not have an impact, the double mutant KD➔A abolished NL63 binding and blocked internalization of SARS2-RBD. The internal region represents a second ACE2 site, when mutated, abolishes nanomolar SARS2-RBD binding, consistent with the prediction that high affinity SARS2-RBD binding is distributed. The K353A, D355A double mutation is more proximal to the collectrin (neck) region (residues 616-742) of ACE2 involved in its dimerization (76). It will be interesting to determine if internalization of ACE2 is proceeded by RBDs binding to ACE2 dimers, which then reduces the affinity for the neck region dimers leading to ACE2 internalization. Such internalization is capable of pinching off membrane from neighboring cells with membrane-bound RBD proteins. Finally, post-translational modifications within the RBD such as asparagine deamidation in N405, N481 and N501 (77, 78) could alter binding to ACE2, however, OM.BA.1 and XBB.1.5 have Y501 between the critical T500 and G502 residues involved in internal region 2 binding. It is worth noting that NL63-RBD interacts with K353 and D355 of ACE2 with a similar three-residue stretch, S535-P536-G537 (48) vs. T500-N501/Y501-G502, again suggesting a common interaction mechanism.

Based on the RBD mutants identified here, it appears the most critical RBD region for interaction with ACE2 may be further refined to a 23-residue region from C480 to G502, which encompasses the Loop 4 disulfide bond (C480-C488). It is noteworthy that this region represents the highest density of residue contacts specific to the m396 neutralizing antibody against SARS2 (28, 79). The disruption of RBD variant binding with the double mutants C480A, C488A and T500A, G502A suggest this region remains critical for any new variant binding to ACE2. It is possible that a peptide inhibitor against this 23-residue stretch could be the basis of a competitive or allosteric inhibitors. Several SARS2-RBD and ACE2 peptide inhibitors have been successfully developed (80, 81) that can mitigate binding. With the emergence of new variants, some of these inhibitors will likely lose efficacy. Although none of our KGD or RGD containing peptides antagonized binding of RBDs, it remains possible that a cyclized or other derivative peptide could do so in the future (49).

It has been suggested that infections through other RBD receptors could negate the attempts to mitigate RBD-ACE2 interactions to control the spread of SARS2 (82-84). While this is true, ACE2 remains the entry protein for two distinct coronaviruses, suggesting it is the target protein that is best expressed on the target cells for viral propagation. It is worth noting that the early phase of SARS2 infections in the nasal epithelium target cells allow for viral amplification with minimal IgG response. The RGD/KGD motif on the SARS2 spike protein has also been implicated in binding to integrins and RGD/KGD binding sites on ACE2 may be competing for co-receptor usage (85-87). Both possibilities underscore the complexities in the SARS2 infection process. Disrupting RBD binding in the nasal epithelium would mitigate all these possible co-receptor interactions.

A model emerges (**See Figures 12, S17**) whereby the Internal region 2 on ACE2 may be critically important for stable RBD-ACE2 binding/internalizations but require secondary contacts for stabilization. In the case of NL63-RBD, those additional contacts are in the Internal region 1 and for SARS-RBDs, those additional contacts are in the Nt proximal and distal regions. It still may be possible to identify a high affinity agonist to RBD based on the NL63/SARS2 target KGD motif in ACE2 that could mitigate binding. Finally, peptides against both the Nt and Internal region binding sites might work synergistically to inhibit the spread of SARS-CoV-2.

**Figure 12:**
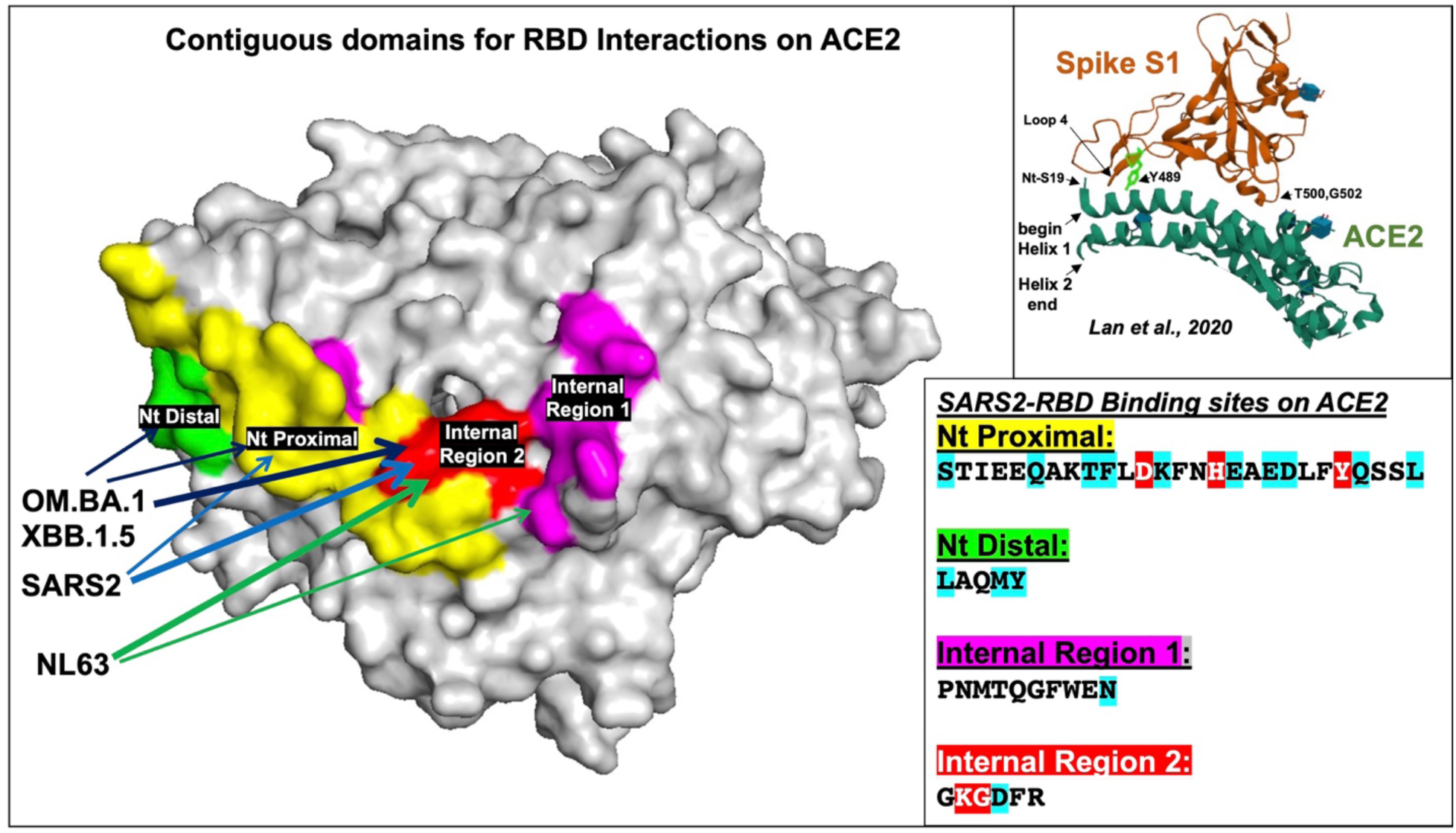
Proposed interpretation of RBD interactions on ACE2 structure. Space filling model of ACE2 structure from pdb_6M0J (27) modified by highlighting interactions for both SARS2 and NL63 RBDs: Nt proximal (yellow), Nt distal (green), and Internal regions 1 (purple) and 2 (red). SARS2-RBD does not bind/internalize if *either* Nt proximal or Internal region 2 (K353, D355) are mutated. OM.BA.1 and OM.BA2-RBDs are not bond/internalized if *either* mutation in both Nt proximal and distal ACE2 or Internal region 2 (K353, D355). NL63-RBD requires Internal region 2 (K353 and D355) for both binding/internalization. **Top Inset** shows crystal structure of Spike S1(burgundy) and RBM of ACE2 (green) modified from Lan et al, 2020. ACE2 residues 19-52 are represented by Helix 1 and 55-81 by Helix 2. Y489 residue shows a main interaction with Helix 1. N487 within Loop 4 interacts with Q24 (not shown). **Bottom Inset** shows **RBD** interactions within the four domains (**see Figures 7, 11**) (residues with blue highlights are predicted binding sites that occur in ¾ of SARS2, SARS1, RaTG13, and NL63-RBDs (26-28); residues with red highlights are predicted binding sites in all 4 RBDs). The RBD interaction data taken together suggest that Internal region 2 binding for SARS1/SARS2/RaTG13 is weak and requires Nt proximal and distal contacts for stabilization (dark blue, light blue arrows), whereas NL63 has additional contacts in the Internal region 1 (green arrows) that supports binding to Internal region 2 (thick arrows). The ability of Y489A to nearly abolish SARS2-RBD binding to ACE2 suggests that none of the domains by themselves have strong binding to ACE2. Overall, the data suggests that Internal region 2 is critical for internalizations of RBDs.

#### Implications

*ACE2 downregulation:* The rapid downregulation by a membrane-bound RBD independent of stable binding suggests that the efficiency of RBD binding and internalization might limit availability of ACE2 protein for future infections. It’s unclear how many full-length Spike proteins that are membrane-bound exist on a virion. On the other hand, the S2-associated S1 particle may mimic the membrane-bound RBD interactions observed. Thus, the loss of ACE2 protein at the plasma membrane before S2 liberation of the Fusion Peptide, might lead to fewer viral particles that can infect target cells. We do not know the rate of *in vivo* ACE2 internalization by RBD and its eventual replacement to the plasma membrane, but this could be studied in the future using mice expressing the human ACE2 protein and membrane-bound RBDs. Reduction in levels of ACE2 will likely affect the Renin-Angiotensin System by increasing Angiotensin II and over activating Angiotensin II receptor type 1 (AT1R), potentially leading to long-term physiological consequences such as Long Covid(30).

The binding assay shown here reveals properties of RBD distinct from other observations that show the SARS2 pseudotype virus causing ACE2 degradation (88-90). It seems plausible that vaccines using RBD domains capable of binding ACE2 may lead to unwanted side effects produced from downregulation of ACE2 in recipients, such as short or long-term alterations in the Renin-Angiotensin System (RAS)(91-93). In addition, a combination of an immune response to the antigen and alterations in RAS may also lead to unwanted side effects (94). In this regard, Pfizer produced several versions of RBD vaccines; if this hypothesis is true, then observed vaccine side effects may have similarities to some of the COVID-19 symptoms. Notably, the full-length Spike protein has fewer side effects than just the RBD alone (95, 96). It is worth exploring the possibility of fewer side effects with vaccines expressing RBD containing mutations such as Y489A, P.B.S. 7Ala, or double mutants C480A-C488A and T500A-G502A that reduce binding to ACE2. The latter two double mutants can work across three variants.

*Internal regions of ACE2:* The published NL63-RBD affinities to ACE2 are similar those of SARS2, but likely utilize internal regions of ACE2 as the key region for its binding and internalization. It is worth noting that NL63-RBD is more robust at reducing ACE2 fluorescence and **Figure 11A’**, which also suggests that the better a protein binds to the internal ACE2 region, the more robust its internalization. The changing nature of SARS2-RBD binding to ACE2 argues that variants could emerge that rely more on the ACE2 internal region for binding like NL63-RBD. It is unclear how COVID-19 disease progression would occur if ACE2 were to be more rapidly internalized.

A nanobody or peptide blocking strategy targeting the KGD region of ACE2 within the nasal epithelium could radically reduce the ability of SARS2-CoV-2 to infect sustentacular cells. Such a peptide might feature the consensus binding residues Ser/Thr-X-Gly found in all SARS2 and NL63 RBDs described here. Mitigating viral amplification in the nasal cavity will likely lead to reduced viral particles, fewer symptoms, and most importantly less asymptomatic spread, which could lead to an improved prognosis for the current and future pandemics.

#### Limitations of the study

The experiments described here are aimed at generating a simple in vitro system for the rapid characterizing the Spike protein-ACE2 interaction domain, specifically the receptor binding domain (RBD). I extend the findings by Starr et al and Chan et al, who performed structure-future relationships between RBD and ACE2 interactions. The focus on the RBD alone is partly because the full-length Spike protein is pre-cleaved, generating S1 and S2 fragments that remain noncovalently associated. There may be value in testing some of these mutations in the context of a pseudovirus. We could test a few mutations in the context of membrane-bound S1 Spike proteins, but the net result would likely be adding more steric hindrance to ACE2 binding as has been shown for furin-mutated Spike protein (64).

I was surprised that the addition of a transmembrane to RBD not only bound to ACE2-Cherry but also reduced fluorescence levels. Loss of ACE2-Cherry fluorescence was not dependent on a β2AR-GFP expressing cells as well. Finally, cell adhesion between differentially labeled CDH2 molecules did not lead to loss of CDH2-Cherry. The observed correlation between the degree of fluorescence loss and the degree of RBD binding is decorrelated when XBB.1.5-RBD is used in the context of an ACE2-Cherry (Figure S3C), ACE2-Cherry (Pus8AA), or when exogenous RBD protein is added. The decorrelation between binding and loss of ACE2 fluorescence suggests that ACE2 may lose its fluorescence due to internalization, and this event is mediated by a disruption of its ability to dimerize.

Because RBDs from both SARS2 and NL63 differentially bind the same spectrum of residues on ACE2, it becomes unclear if these are the only residues necessary and sufficient for RBD internalization. In this regard, residues involved in ACE2 dimerization may indirectly affect loss of ACE2-Cherry fluorescence upon membrane-RBD binding.

## Materials and Methods

### RBD and ACE2 constructs

See details of DNA sequences in **Supplemental Data File S1**. D1794 (is the β2AR with an internal EcoR1 site added (does not alter residue) for rapid cloning of 5’ends. ACE2-Cherry (human ACE2 coding sequence with an internal EcoR1 site mutation (does not alter residues). RBD and ACE2 mutants made by gblocks (IDT) and sequenced after subcloning (see **Supplemental Data File S1**). SARS1 RBD from (23), SARS2 and RaTG13 are equivalent regions. NL63 RBD from NCBI. All RBD variants have been codon optimized.

hemagglutinin-cleavable signal sequence: (MKTIIALSYIFCLVFA) was initially used to test secreted version (25). All membrane tethered Nt RBD constructs used nicotinic receptor α7 subunit signal peptide: (MGLRALMLWLLAATGLVRESLQGEFQRKLYPVATM) (32, 33).

### Cell culture and Imaging

Mouse olfactory placode cells [OP6 (97), a gift from Jane Roskams], were maintained as previously described in Dulbecco’s modified Eagle’s medium (DMEM 1X Gibco) supplemented with 10% fetal bovine serum (FBS Gibco) and 1% penicillin/streptomycin (Pen Strep Millipore) at 33°C. I transiently transfected plasmid DNA constructs using the Amaxa Nucleofector (Lonza) with PBS at 60%-70% confluency according to the manufacturer’s protocol. Transfected cells were allowed to recover for 16-18 hours at 33°C and express the plasmid DNA before imaging.

I have previously characterized β2AR and odorant receptor fluorescent protein fusions (20-22). Cells grown at 33°C and imaged 16hrs post-transfection. Typically, 15ug of DNA was electroporated (EPd) into 0.5 million OP6 cells. Imaging performed LSM510 confocal microscope or Spinning Disk microscope as previously performed (20).

Electroporation of 0.5 million OP6 cells with ACE2 plasmid (Cherry) is plated and grown overnight, and expression was typically found in 1-5% of cells. Electroporation of 0.5 million OP6 cells with an RBD or other plasmids are plated and grown overnight. Expression of these plasmids was typically 2-4 fold better than the ACE2 containing plasmids.

### Analysis of flow cytometry values

Trypsinized and analyzed 100K-250K DAPI (live cells). Green, Red and Green&Red cells. Fluorescent counts acquired. Total green fluorescent cells = (Green&Red Cells + Green cells) divided by DAPI total. Normalize total green cells are based on the experimental control counts. Total red cells= (Red&Green cells + Red cells) divided by DAPI total. Normalized total Red cells are based on the experiment controls for each experiment. Total Red&Green cells were obtained by gating and dividing by Green Cell value and normalized by controls within an experiment, then multiplied by Normalized Green Value.

### RBD/ACE2 Interaction analysis

- **In vitro co-plate binding assay**: RBD and ACE2 DNA constructs were separately EPd into OP6 cells, seeded (co-plated) onto the same 30cm^2^ dishes, and imaged after 16hrs. Internalization of green fluorescence by red cells was obvious whether green cells were in proximity. When internalization was not observed, then images were captured for at least 3 cells where Green/Red cells are touching.
- **In vitro co-EP assay:** 10ug of RBD and 10ug of ACE2 DNA constructs were Co-EPd into OP6 cells, seeded onto 30cm^2^ dishes, and imaged after 1μ6hrs. At least 3 coexpressing cells were imaged for each.
- **Dissociation-Plate-Flow cytometry (*DPF*) assay:** 15ug of RBD and 15ug of ACE2 DNA constructs were separately EPd into OP6 cells and seeded onto different 30cm^2^ dishes. RBD cells were trypsinized and seeded for the desired time onto the 16hrs grown ACE2-expressing cells in 30cm^2^ dishes, then trypsinized, spun down, resuspended, filtered and counted by flow cytometer.
- **In solution binding assay:** 15ug of RBD and 15ug of ACE2 DNA constructs were separately EPd into OP6 cells and seeded onto different 30cm^2^ dishes. Both RBD cells and ACE2-expressing cells were trypsinized, spun down, resuspended in 300μl of media. 50μl of each population were mixed into an Eppendorf tube and put in a 37°C water bath for 10min to 1hr. A 10μl aliquot was pipetted onto coverslip and imaged. Images were taken such that many cells were in the field of view. If doublets formed, then it was apparent throughout the field of view.

Flow cytometry and FACS were performed using a Becton Dickinson Aria II equipped with 5 lasers (BD bioscience). Gates were drawn as determined by negative controls to separate positive and negative populations for each florescent protein. Typically, 300,000 events (yielding about 250,000 DAPI positive cells) were recorded for each flow cytometry analysis, and the data was analyzed using BD FACSDiva 9.0.1. Calculations and graphs were made in GraphPad Prism. Mean and standard deviations were derived in Excel. See **Supplemental Data File S2** for details.

### Peptides and antibodies

Peptides synthesized from GenScript. RGD purchased from R&D systems, #7723/10mg. Cilengitide purchased from Tocris, #5870.

His Tag Alexa Fluor 488-conjugated Antibody Catalog #: IC050G from R&D systems. 2019-nCoV RBD, His tag Cat#40592-V08H from SinoBiologicals

### Live-imaging

OP6 cells transfected with RBD and/or ACE2 constructs imaged on a Zeiss LSM 510 microscope using a Zeiss Plan-APO 25X water immersion objective in culture media for Figures 1, 2, 3, 6, S1, S4, S9 and S10. GFP/Venus expressing cells excited at 488- collected at BP500-545 and Cherry expressing cells at 561 nm-collected at LP575 nm. All images were acquired using the multitracking feature.

### Logo plots and space filling model of ACE2

https://weblogo.berkeley.edu/logo.cgi used to generate Logo plots in Figures 7, 11.

Modifications of space filling model of ACE2 structure from Wu et al, 2020 (pdb_00006m0j) used to generate Figures 12 and S17.

### β2AR antagonist

CellAura fluorescent β2antagonist [(±)-propranolol], HB7820 from Hello Bio https://www.hellobio.com/cellaura-fluorescent-beta2-antagonist-propranolol.html?q=7820 (98).

His Tag Alexa Fluor® 488-conjugated Antibody Catalog #: IC050G from R&D systems. 2019-nCoV RBD, His tag Cat#40592-V08H from SinoBiologicals

### Odors

Le Nez du Café Revelation Kit Odors-36 Scent concentrations from Expresso Parts.

## Acknowledgements

Special thanks to Masayo Omura for providing logistical and intellectual support that helped drive the success of these experiments. Many thanks to Masayo Omura, Ivan Rodriguez, Sergio Bernal-Garcia, and Thomas Bozza for advice on the experimental design and interpretation of results; Mary Slavinsky and Masayo Omura for technical support in plasmid production; Livia Johnson and Diana Bratu for advice on utilizing the spinning disk microscope; and Sergio Bernal -Garcia for help with GraphPad Prism and Fiji. Thanks to Ivan Rodriguez, Vincent Racaniello, Masayo Omura, and Sergio Bernal Garcia for providing feedback on the manuscript. Tomas Baumgartner Manager, Flow Cytometry Core Facility, Weill Cornell Medicine for support on flow cytometry of single cells. GISAID (http://cov-glue-viz.cvr.gla.ac.uk/mutations.php) for SARS-CoV2 database. Many thanks to Delia Tomoiaga, Claude Rose Levine, and Mark Levine for supporting my efforts to design novel experiments that might help tackle the pandemic of our lifetime. This work was supported by Hunter College and NIH grants# NIH R01 NS091439-01 and NIH R01 DC020764-01.

All data generated are found in the supplementary files of this manuscript.

## Supplemental information inventory

### Supplemental Figures and legends

**Figure S1:**
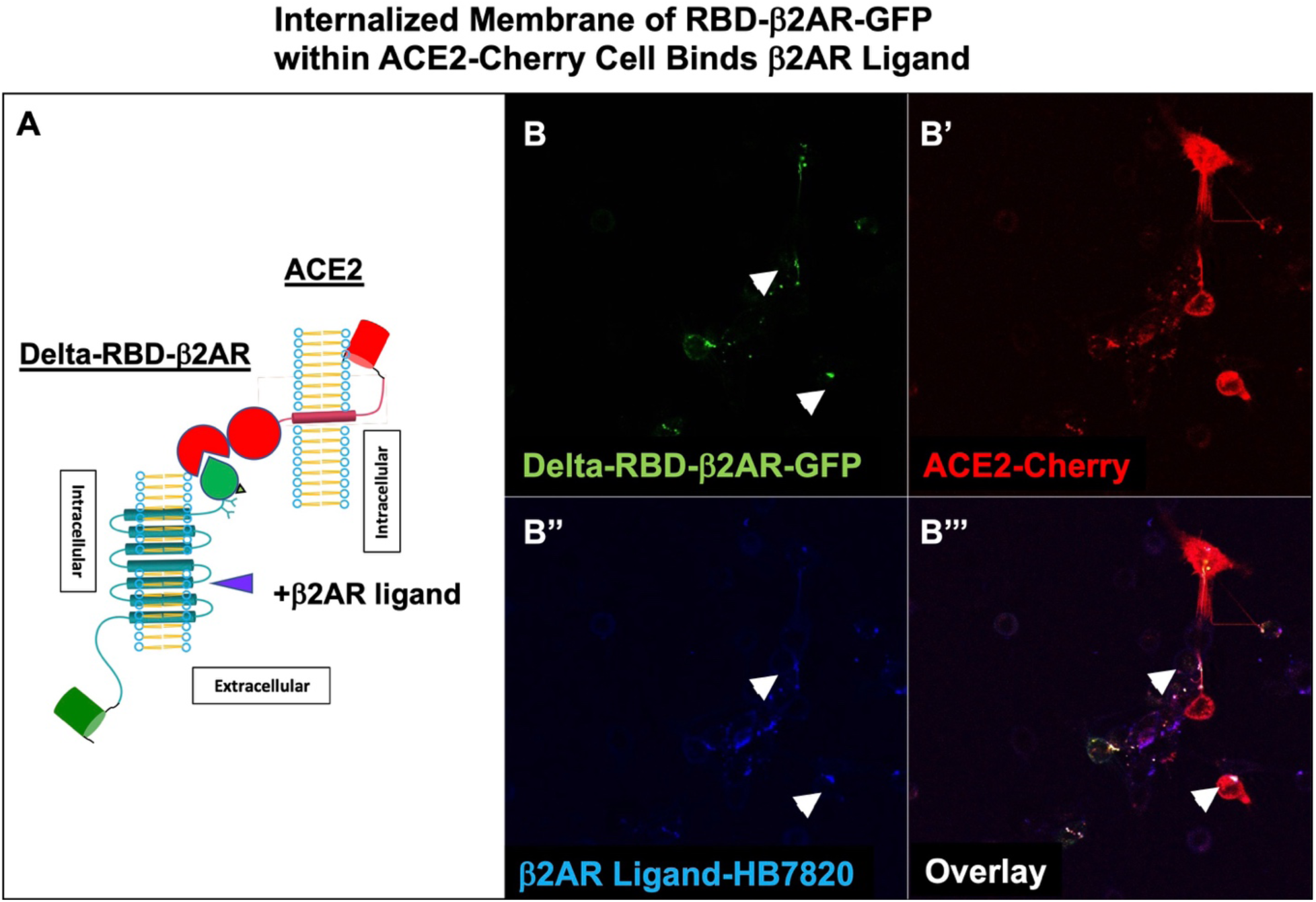
Internalized Membrane of RBD-β2AR-GFP within ACE2-Cherry Cell Binds β2AR Ligand. **A.** Schematic of protein interaction and a β2AR ligand binding**. B-B’’’.** Overnight Delta-RBD-β2AR-GFP co-cultured with ACE2-Cherry cells. β2AR fluorescent ligand HB7820 (70nM, 1:500 dilution of 35μM stock) added to media for 10min and imaged. **B**. Green fluorescence from Delta-RBD-β2AR-GFP. **B’.** ACE2-Cherry expressing cells show Delta-RBD-β2AR-GFP internalized along with ACE2-Cherry. **B’’.** β2AR ligand HB7820 fluorescence. **B’’’.** Overlay shows β2AR ligand HB7820 bound to Delta-RBD-β2AR-GFP within ACE2 expressing cells (white arrowheads).

**Figure S2:**
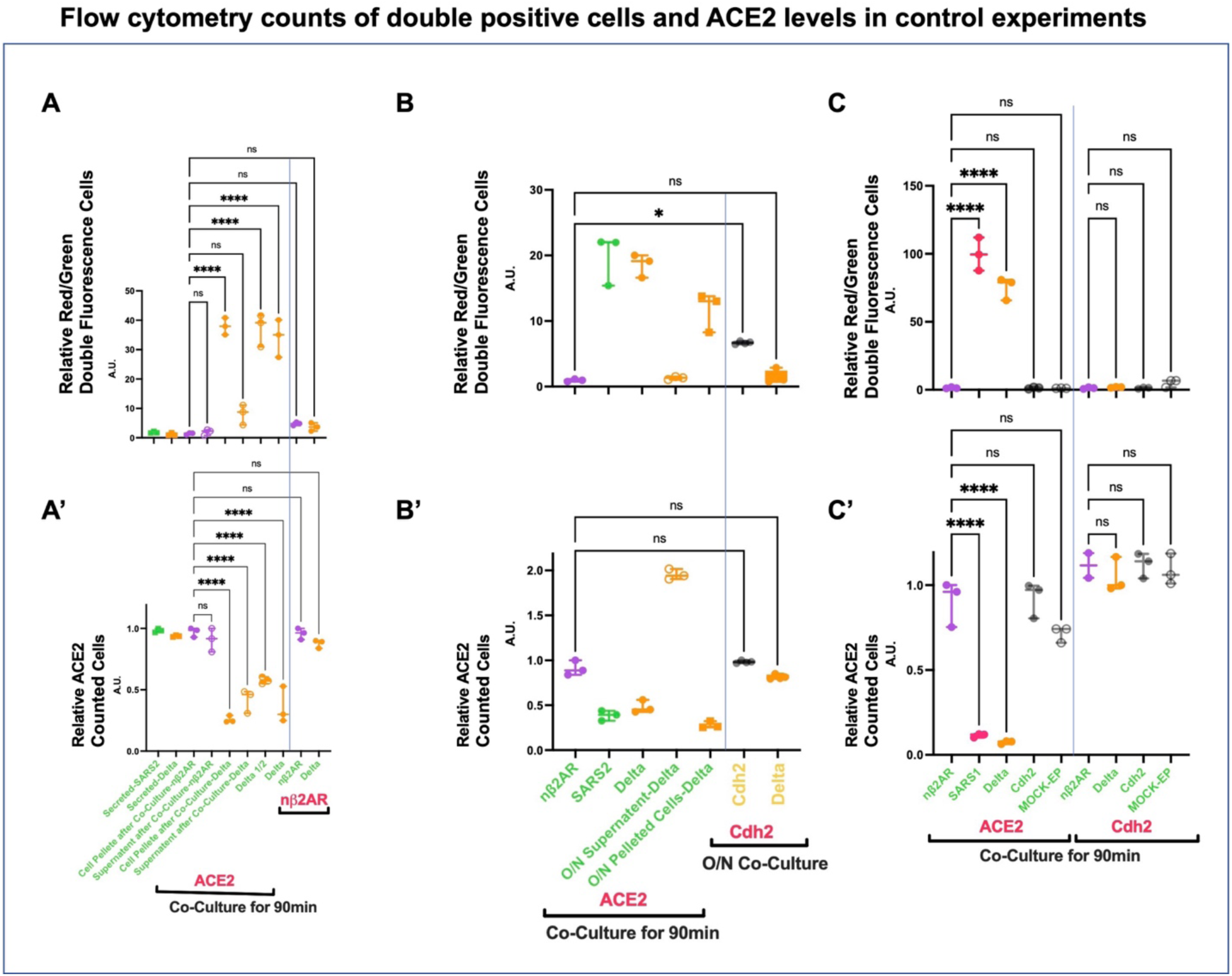
Flow cytometry counts of double-positive cells and ACE2 levels in control experiments. **A-C** *DPF assay*: Values are normalized to controls (Y-axis arbitrary units, A. U.). Overnight (O/N) transfection and flow cytometry counts of Co-plating GFP or Venus expressing cells at 90min with RBD or Cdh2 with ACE2-Cherry, nβ2AR-Cherry, or Cdh2-Cherry expressing cells followed by trypsinization. **A’-C’.** *DPF assay*: Values are normalized to controls (Y-axis arbitrary units, A. U.). relative to controls. Green labeling represents GFP fusion proteins. Red labeling represents Cherry fusion proteins. Yellow labeling represents Venus fusion proteins. **A and A’.** Secreted-SARS2 or Secreted-Delta represents the non-transmembrane RBD expressing cells. No double fluorescent cells (green and red) were observed. Supernatants were collected separately from attached cells for nβ2AR or Delta-RBD-nβ2AR (Delta) expressing cells after co-culturing with ACE2-Cherry for 90min. No double fluorescent cells (green and red) were observed in the supernatant relative to control. Only attached cells showed double fluorescent cells for Delta-RBD-β2AR as expected. **B and B’.** SARS2-RBD-nβ2AR (SARS2) shows indistinguishable results to Delta: more double fluorescent cells relative to controls and loss of ACE2-Cherry fluorescence. Supernatant from overnight (O/N) growth of electroporated Delta cells do not yield double fluorescent cells when plated onto ACE2-Cherry cells but attached cells that were incubated with ACE2-Cherry yielded double fluorescent cells and loss of ACE2-Cherry fluorescence. O/N incubation of Cdh2-Venus with Cdh2-Cherry leads to double fluorescent cells (yellow and red), but not Delta-nβ2AR-Venus with Cdh2-Cherry. No loss of Cdh2-Cherry fluorescence is observed with either incubation. **C and C’.** nβ2AR, SARS1-RBD-nβ2AR (SARS1), Delta, Cdh2, or Mock-EP are co-plated with ACE2-Cherry for 90min. Only SARS1 and Delta show double fluorescence and loss of ACE2-fluorescence. In a separate set of experiments, nβ2AR, Delta, Cdh2 and Mock-EP are co-plated with Cdh2-Cherry for 90min. No double fluorescence for any of these four incubations, nor loss of Cdh2-Cherry fluorescence, was observed. It should be noted that Cdh2-Venus/Cherry double fluorescence was not observed in 90min compared to the O/N co-plating.

**Figure S3:**
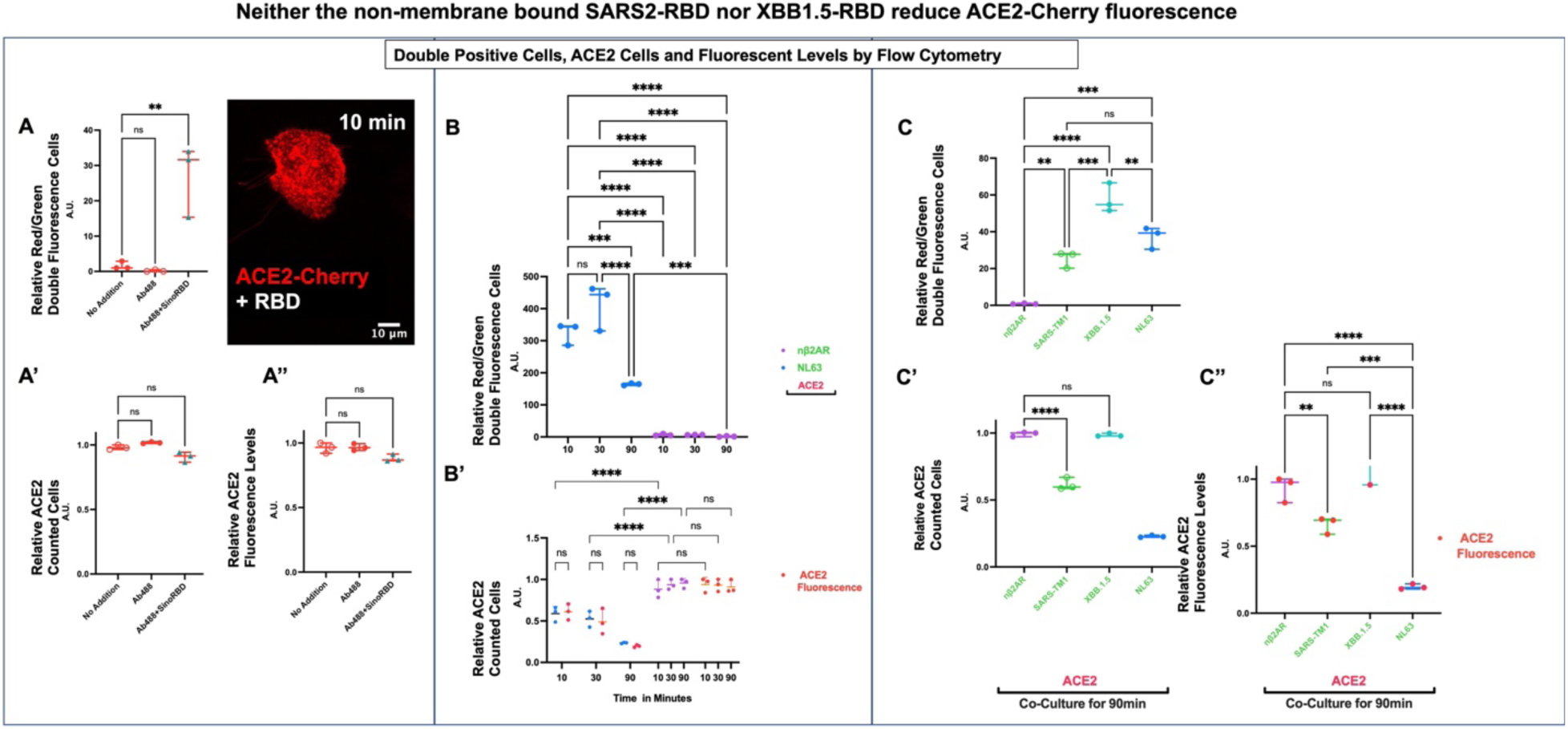
Neither the non-membrane bound SARS2-RBD nor XBB1.5-RBD reduce ACE2-Cherry fluorescence. **A-A’’.** *DPF assay*: values in are normalized to controls (Y-axis arbitrary units, A. U.). This data makes is also shown in **Figure S16**. **A**. Double fluorescence after 90minute incubation of ACE2-Cherry cells with Ab488 (1μg/4mls of Goat-anti-HIS-Tag-Alexa488 Goat antibody [R&D#: IC050G]) and with or without Sino-RBD, purified His tagged-RBD protein (60ng/4mls of 2019-nCoV-HisTag-RBD [Cat#40592-V08H; 234 residues, R319-F541]). **A’** ACE2-Cherry cells with Ab488 or Ab488+Sino-RBD show no changes in fluorescence cells. **A’’** ACE2-Cherry only cells also show no changes in fluorescence values. **Inset,** Shows ACE2-Cherry cell plus 240ng/5mls Sino-RBD. Punctate fluorescence is observed on the cell surface. **B.** Double fluorescence observed for NL63-RBD at 10min, 30min and 90min interaction with ACE2-Cherry plated cells. **B’.** ACE2-Cherry cells show reduced expression overall as well as cells highly expressing cells (red data points) when in contact with NL63 within 10min. Again nβ2AR controls show no difference. **C.** Double fluorescence is observed for SARS2-TM1, XBB.1.5-RBD and NL63-RBD 90min interaction with ACE2-Cherry plated cells. **C’.** ACE2-Cherry cell fluorescence is reduced for SARS2-TM1 and NL63-RBD interactions, but not for XBB.1.5-RBD. **C’’.** ACE2-Cherry cells show reduced expression for highly expressing cells (red data points) in contact with SARS2-TM1 and NL63-RBD interactions, but not for XBB.1.5-RBD.

**Figure S4:**
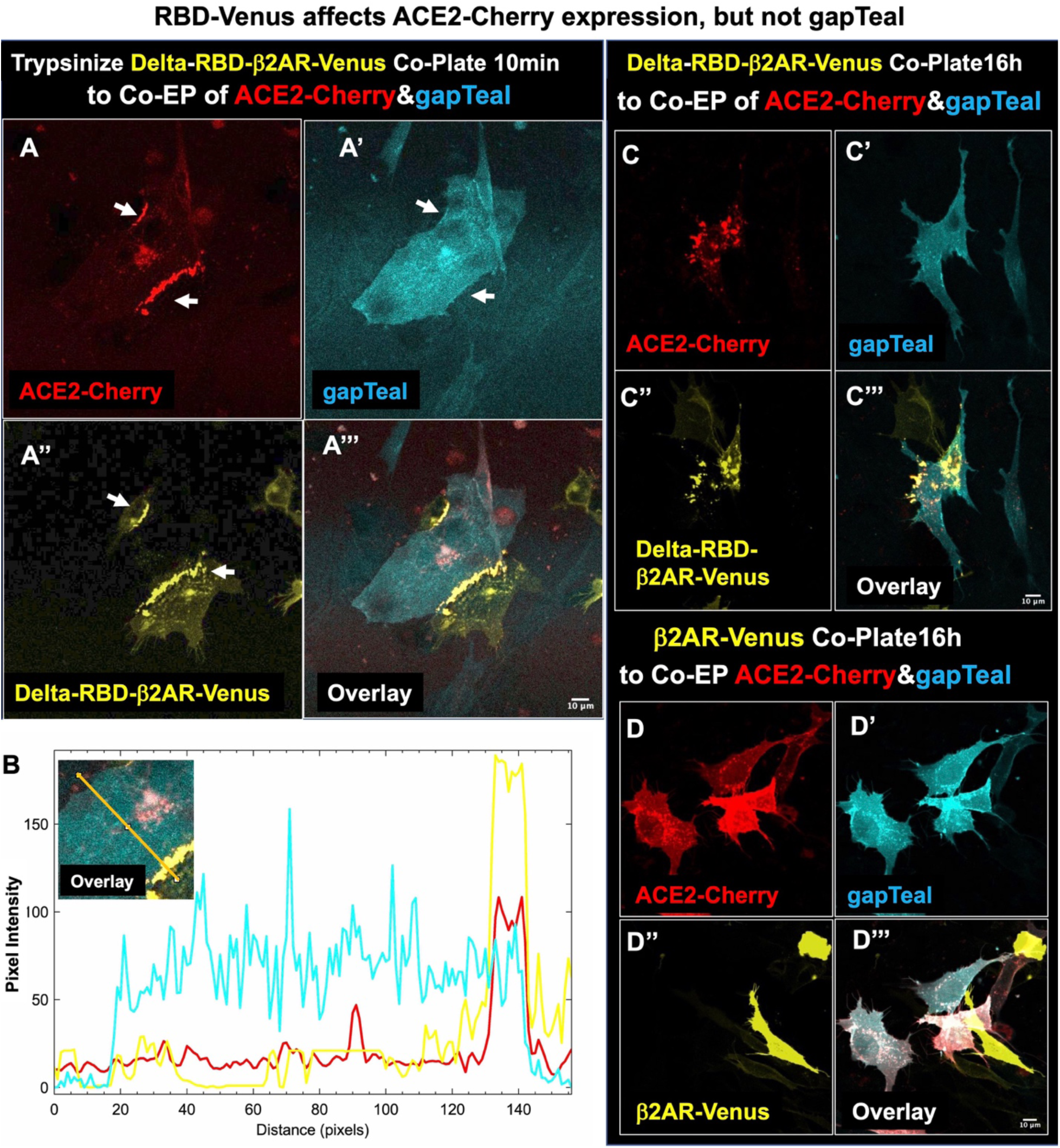
RBD-Venus affects ACE2-Cherry expression, but not gapTeal. **A-A’’’.** Confocal images for Co-EP of ACE2-Cherrry and gapTeal co-plated with Delta-RBD-β2AR-Venus cells from acute trypsinization. **A-A’’’**. In less than 30min, aggregation of both Delta-RBD and ACE2 fusion proteins is observed (white arrows). **B**. Histogram of fluorescence intensity changes also reveals aggregation of Delta-RBD and ACE2 fusion proteins, but not the membrane localized protein gapTeal. **C-C’’’.** 16hr co-plating also shows gapTeal stays membrane-bound despite all the ACE2-Cherry fusion protein found in aggregates. **D-D’’’.** Co-EP of ACE2-Cherrry and gapTeal co-plated O/N with β2AR-Venus cells. No aggregation of β2AR-Venus and ACE2-Cherry protein.

**Figure S5:**
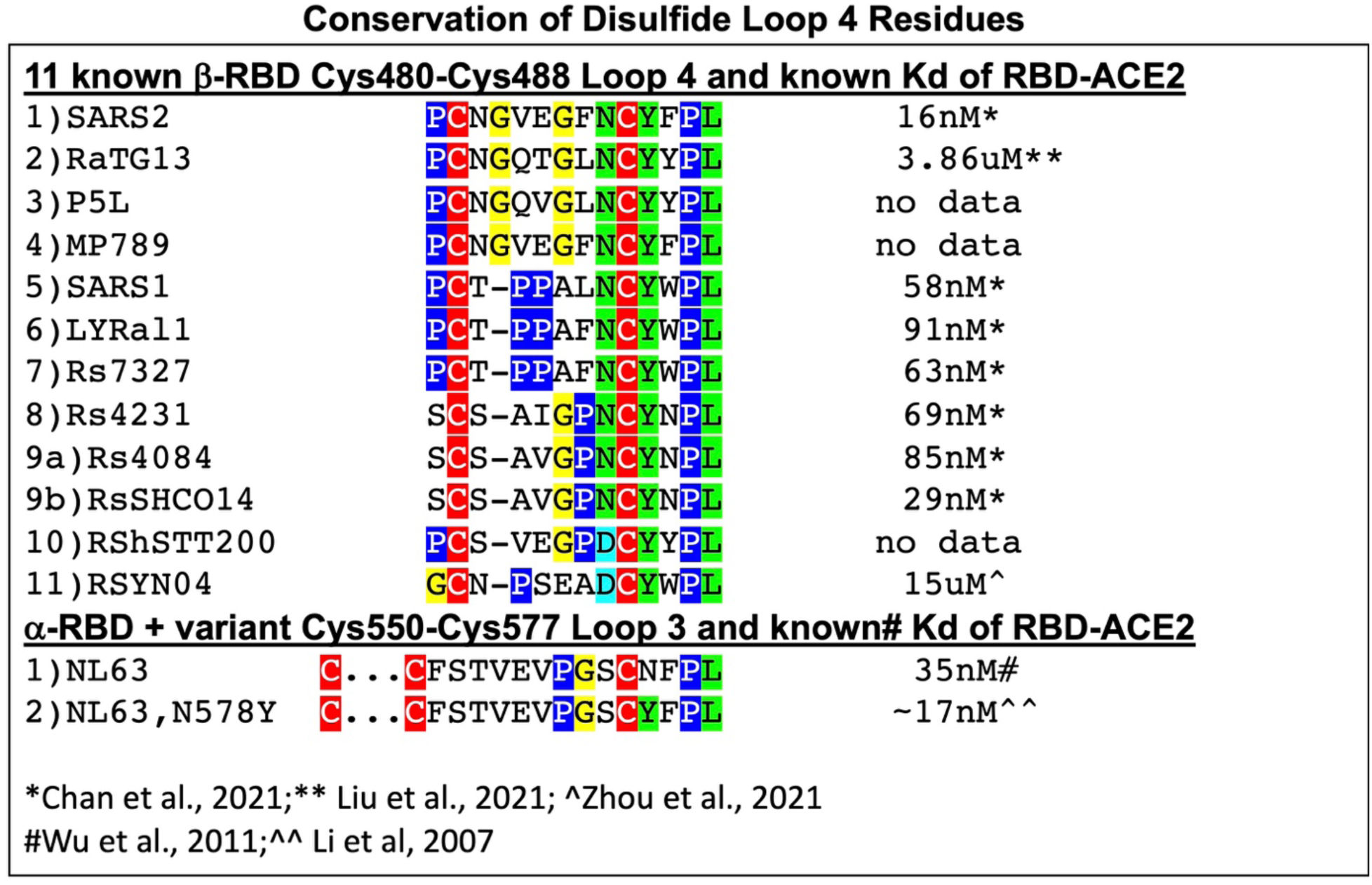
Conservation of Disulfide Loop 4 Residues. A curious finding by Zhou and colleagues found that *Rhinolophus pusillus* virus RpYN06 had strong homology the SARS2 virus but did not contain the disulfide bridge homology between Cysteine-480 to 488 (Loop 4) (**See Figure S7**); The *R. pusillus* RpYN06-RBD did not bind to ACE2 (41). The Loop 4 region that lies within the SARS2-RBM is well conserved among eleven betacoronavirus species, eight of which are known to bind to ACE2 (8, 21, 22, 40, 41, 43, 44). The only homology between the alpha coronavirus NL63-RBD and SARS2-RBD overlaps the Loop 4 region (**See Figure S9**). An N578Y mutation can increase the NL63-RBD binding affinity to ACE2 despite this domain residing outside of the three RBMs. Conserved Cysteines that form disulfide bridges are highlighted in red, Prolines are highlighted in dark blue, Glycines (alpha helical breakers) are highlighted in yellow, and conserved Asparagines, Tyrosines, and Leucines are highlighted in green.

**Figure S6:**
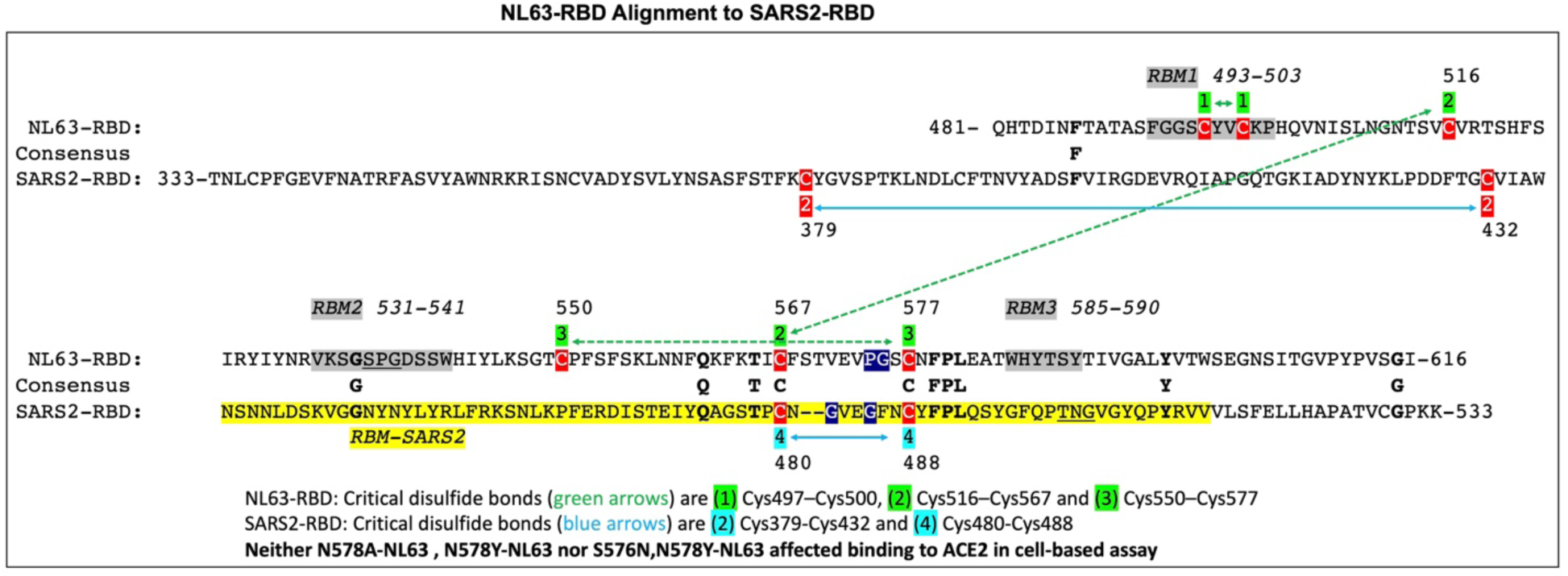
NL63-RBD Alignment to SARS2-RBD. Both NL63-RBD and SARS2-RBD domains both bind and are internalized by ACE2 expressing cells. NL63-RBD contains three RBM domains (grey highlighted regions) and three predicted disulfide bridges (green numbers and arrows), compared to one RBM for SARS2-RBD (yellow highlighted region) and two critical disulfide bridges (blue numbers and arrows). Manual alignment between 136 residue NL63-RBD and 197 residue SARS-RBD. Consensus sequence reveals only 11 amino acid identities. Four of these residues reside in a 5-amino acid stretch that overlaps Loop 4 region within the RBM of SARS2 (40). Loop 4 alignment in Figure S2 identifies conserved Glycines in Prolines between Cysteine residues. These residues (dark blue highlights) are also found in NL63 and near the conserved CxFPL. It is worth noting that the N578Y mutation in NL63 bringing the identity to 5/5 identities CNFPL, doubles the binding affinity to ACE2 (**see Figure S2**). Residues 535-537(SPG) in NL63 and 500-502 (TNG) in SARS2 are underlined and proposed to interact with critical ACE2 residues K353 and D355.

**Figure S7:**
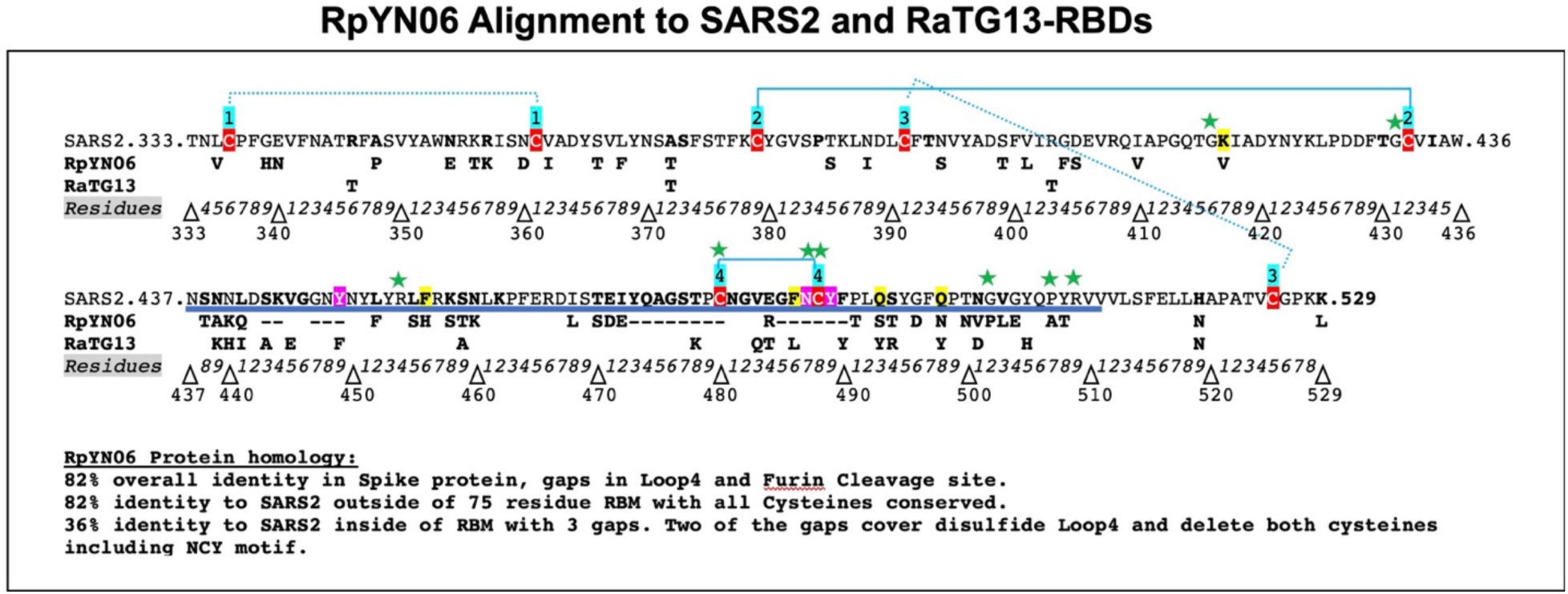
RpYN06 Alignment to SARS2 and RaTG13-RBDs. The RaTG13 betacoronavirus is one of the most highly homologous betacoronaviruses to SARS2 sharing 96% overall identity including the RBM region (residue differences in brown). By contrast, RpYN06 has 95% nucleotide identity over the 30kb colinear genome, but its RBM region has lost nucleotide homology to SARS2 with the best stretch having 22/57 residues conserved (residue differences in blue). It is notable that two of the eight cysteines are not conserved (highlighted in red), and both make up disulfide Loop 4. Loss of RBM homology correlates with loss of ACE2 binding (41, 42)

**Figure S8:**
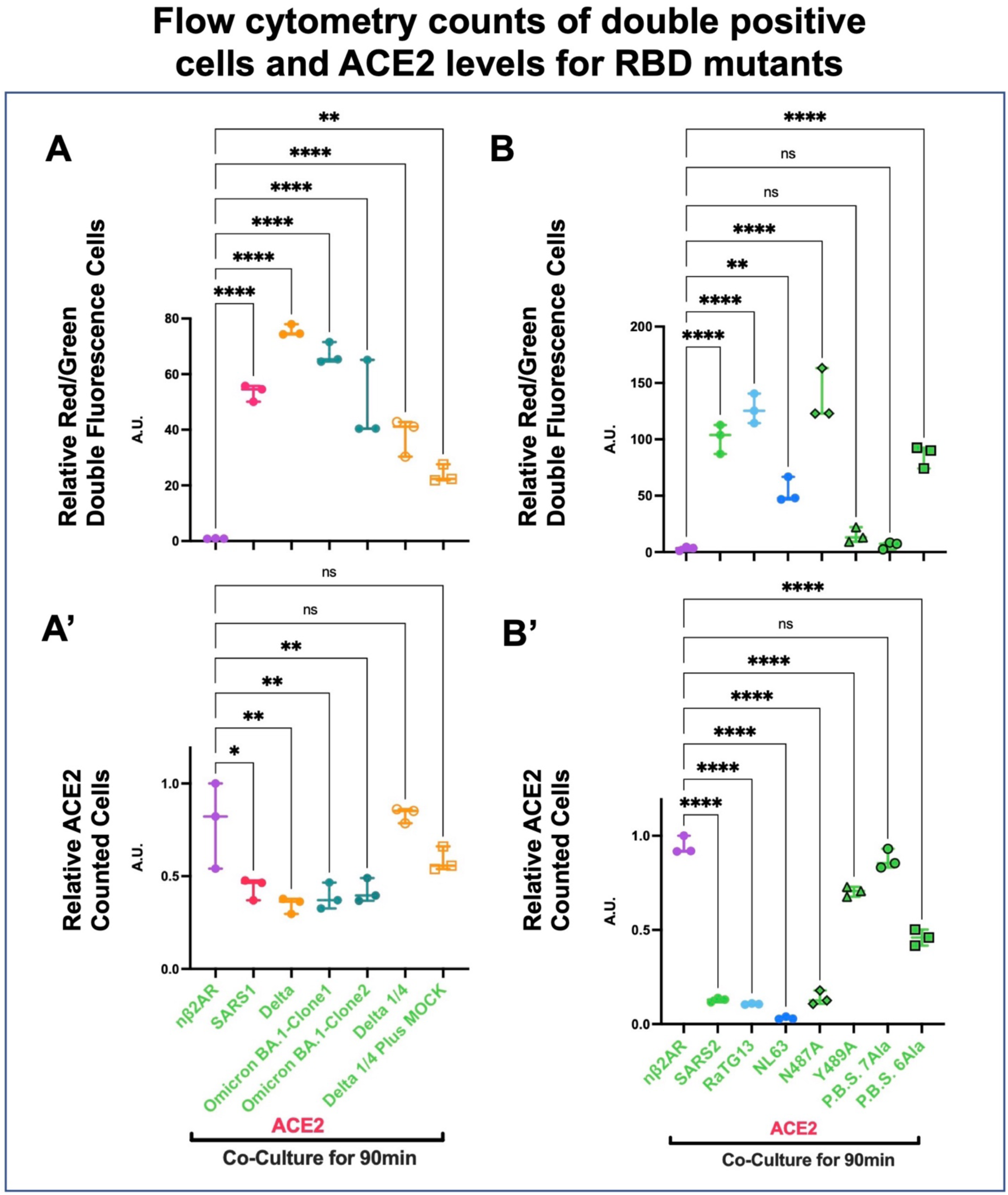
Flow cytometry counts of double positive cells and ACE2 levels for RBD mutants. **A-C and A’-C’.** *DPF assay*: values in are normalized to controls (Y-axis arbitrary units, A. U.). Overnight (O/N) transfection and flow cytometry counts of Co-plating GFP and Cherry expressing cells at either 90min (RBD cells trypsinized and added to ACE2 expressing cells. Green labeling represents GFP fusion proteins. Red labeling represents Cherry fusion proteins. **A-B and A’-B’**. Data shows that double fluorescent cells are found with RBDs of SARS1, SARS2, Delta, Omicron, RaTG13, NL63; Concomitant loss of ACE2 expression was observed. The N487A mutant still robustly bound and ACE2 expression was unperturbed, in contrast to Y489A where nearly all binding is abolished, but still shows reduced expression of ACE2. The 7Ala mutation of the Predicted Binding Site (P.B.S. 7Ala: K417A, G446A, L455A, F486A, Q493A, Q498A, N501A) completely abolished binding and no loss of ACE2 expression was observed. By contrast P.B.S. 6Ala mutation (+F486) generated substantial double positives as well as loss of ACE2 expression.

**Figure S9:**
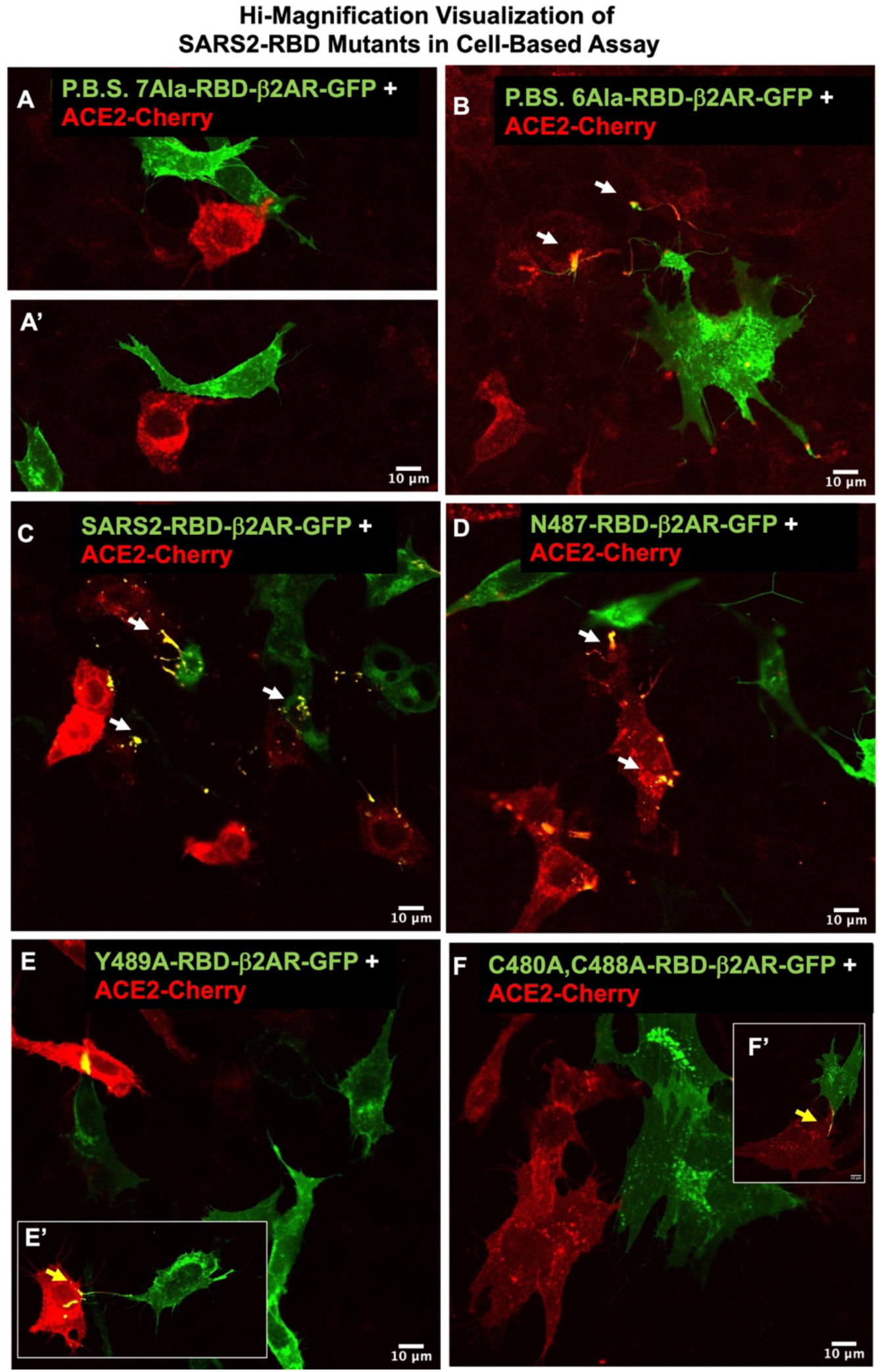
Hi-Magnification Visualization of SARS2-RBD Mutants in Cell-Based Assay. **A-F.** Confocal images for co-plated RBD variants and mutants with ACE2-Cherry after 16hrs. **A and A’.** P.B.S. 7 Ala-RBD expressing cells reveal fluorescent filopodia at the membrane (obvious filopodia), but the protein is not found bound to ACE2-Cherry cells. **B.** P.B.S.6 Ala do bind ACE2-Cherry cells. **C.** SARS2-RBD expressing cells robustly bind ACE2-Cherry cells. **D.** N487A-RBD, **E and E’.** Y489A-RBD and **F and F’.** C480-C488A-RBD with ACE2-Cherry Cells. N487A-RBD interactions with ACE2-Cherry are robust, in contrast to Y489A-RBD and C440-C488A-RBDs, which rarely interact with ACE2-Cherry Cells.

**Figure S10:**
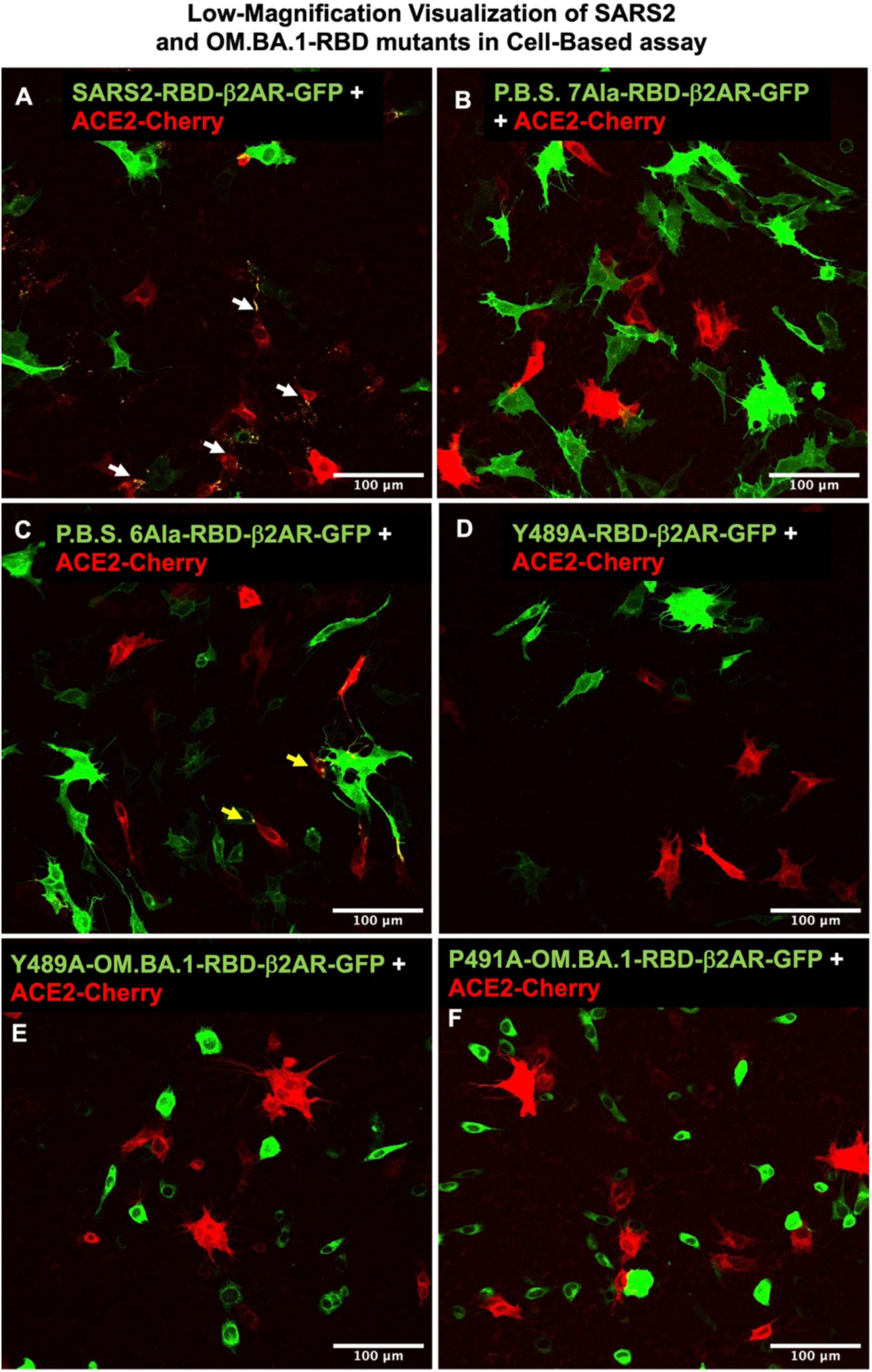
Low-Magnification Visualization of SARS2 and OM.BA.1-RBD mutants in Cell-Based assay. **A-F.** Low magnification confocal images for co-plated RBD variants and mutants with ACE2-Cherry reveals the pervasiveness of RBD-ACE2 interactions/noninteractions, RBD membrane trafficking/non-trafficking. **A.** SARS2-RBD protein is readily internalized by ACE2-Cherry cells and reduces its membrane expression. **B.** P.B.S. 7 Ala-RBD expressing cells are well expressed at the membrane (obvious filopodia), but the protein is not found bound to ACE2-Cherry cells. **C.** P.B.S. 6 Ala Shows binding ACE2-Cherry cells. **D.** Y489A-RBD expressing cells do not show binding to ACE2-Cherry cells. **E.** Y489A-OM.BA.1-RBD and **F.** P491A-OM.BA.1-RBD Show no filipodia and no interactions with ACE2-Cherry Cells.

**Figure S11:**
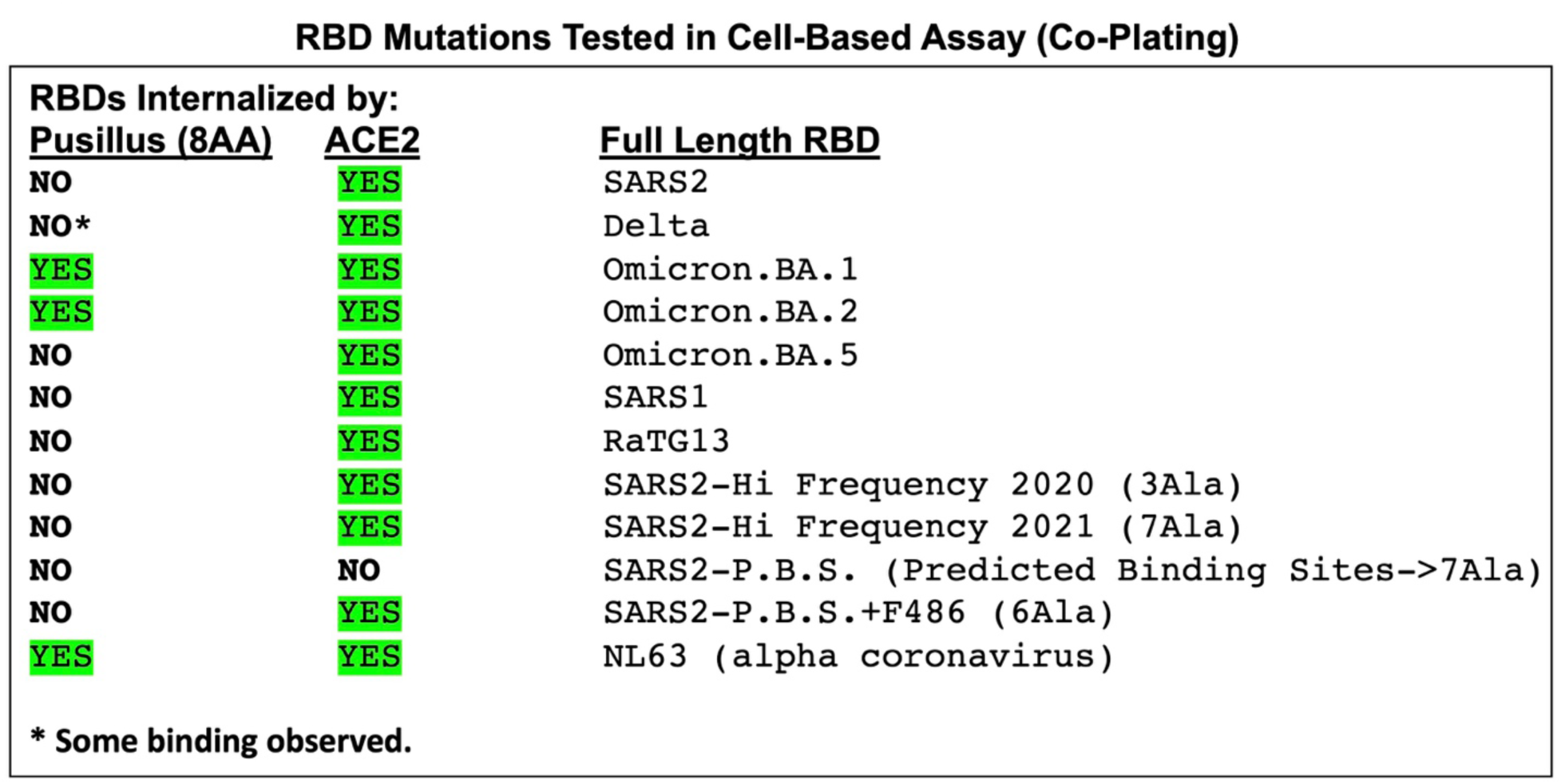
RBD mutations tested with Pus8AA+LMY➔A-ACE2. Co-plating RBD constructs with Pus8AA-ACE2-Cherry cells. A broad set of RBD mutants and variants of SARS2, SARS1, RaTG13 and NL63 are tested by visual inspection **(see also Figure 8)**. Only OM.BA.1-RBD, OM.BA.2-RBD, and NL63-RBD can robustly bind to Pus8AA-ACE2-Cherry. All three SARS2-RBD mutants that could bind to ACE2-Cherry, did not bind to the Pus8AA-ACE2-Cherry.

**Figure S12:**
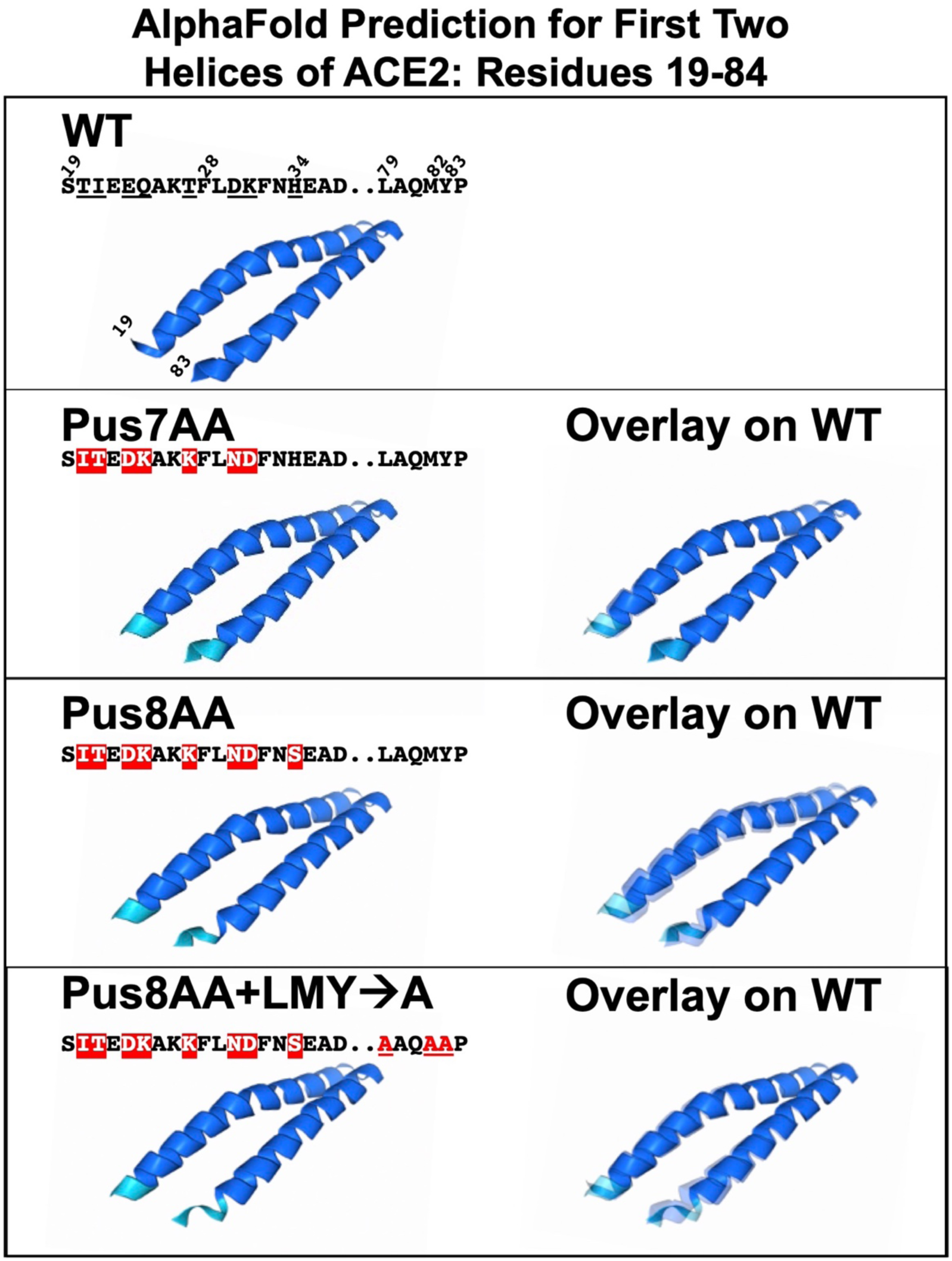
AlphaFold Prediction for the first two helices of ACE2 (residues 19-84) The mature ACE2 protein has its signal sequence cleaved after residue 18. The next 63 residues (19-81) form two helices (helix 1: residues 19-52 and helix 2: residue 55-81). AlphaFoldpredictions (residues 19-84) for WT and Group III mutants: Pus7AA, Pus8AA and Pus8AA+LMY➔A all maintain the two-helix structure. Pus8AA has substitutions in ACE2 residues 20, 21, 23, **24**, **27**, **30**, **31**, **34** (bolded residues are predicted to interact with SARS2-RBD, see also Figure 7). Overlays with WT show only subtle differences in the helices, but more robust changes at the Nt and artificial Ct of the models.

**Figure S13:**
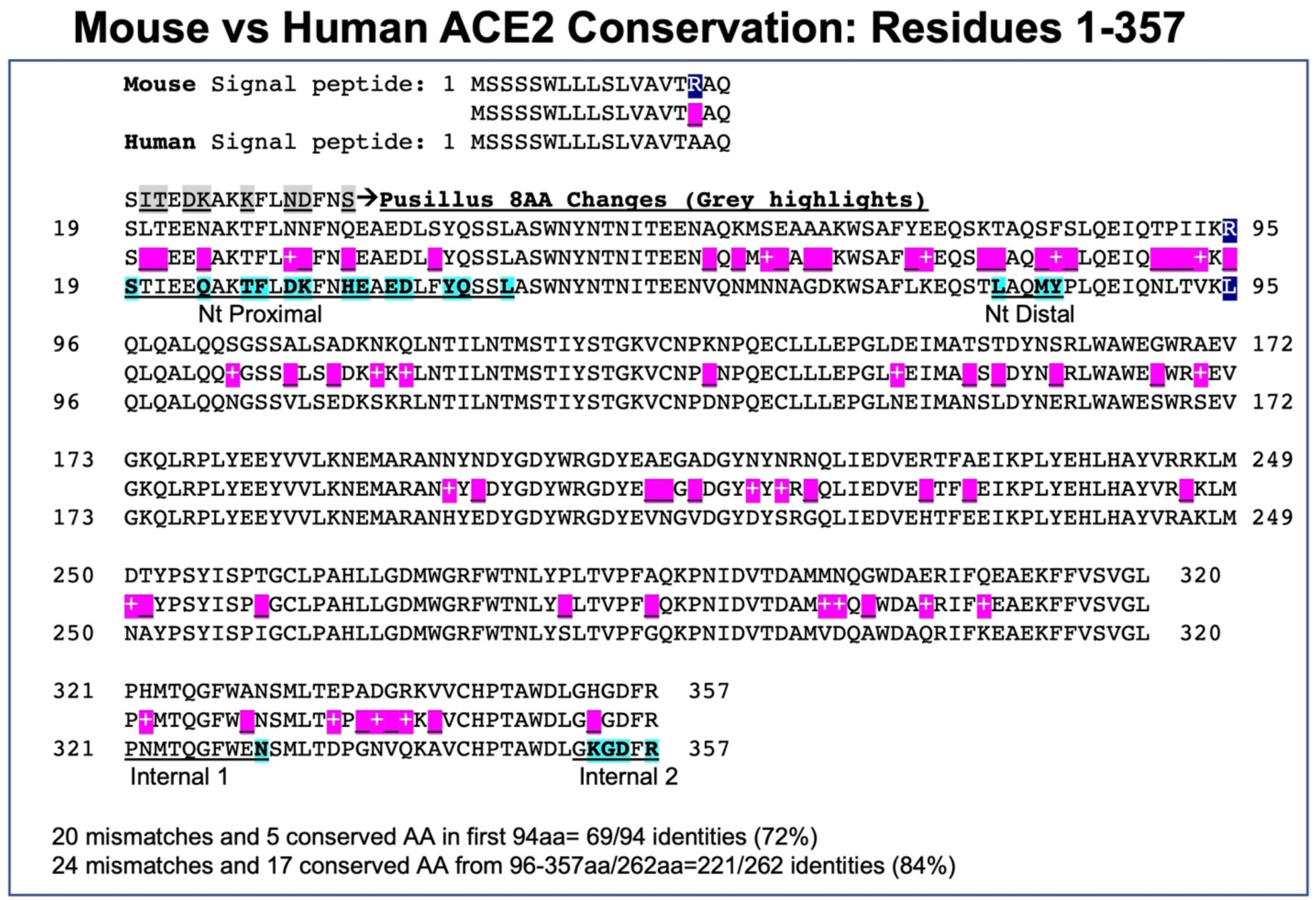
Mouse vs Human ACE2 among first 357 residues. First 357 ACE2 residues compared between mouse and human show 87% identity. The consensus sequence shows 45 mismatches (purple shading) of which 22 are conserved. mACE2 Residue Arg95 has been highlighted (blue shading) and is not contained in the m/hACE2 chimera with residues 1-353 of mA CE2 (**see Figures 8 and 10).** Human residues predicted to interact with SARS2-RBD are bolded and shaded turquoise. Nt-Proximal region (residues 19-45), Nt Distal (residues 79-83), Internal region 1 (residues 321-330) and Internal region 2(residues 352-357) are underlined. Pus8AA chimera (residues 19-34) are shown.

**Figure S14:**
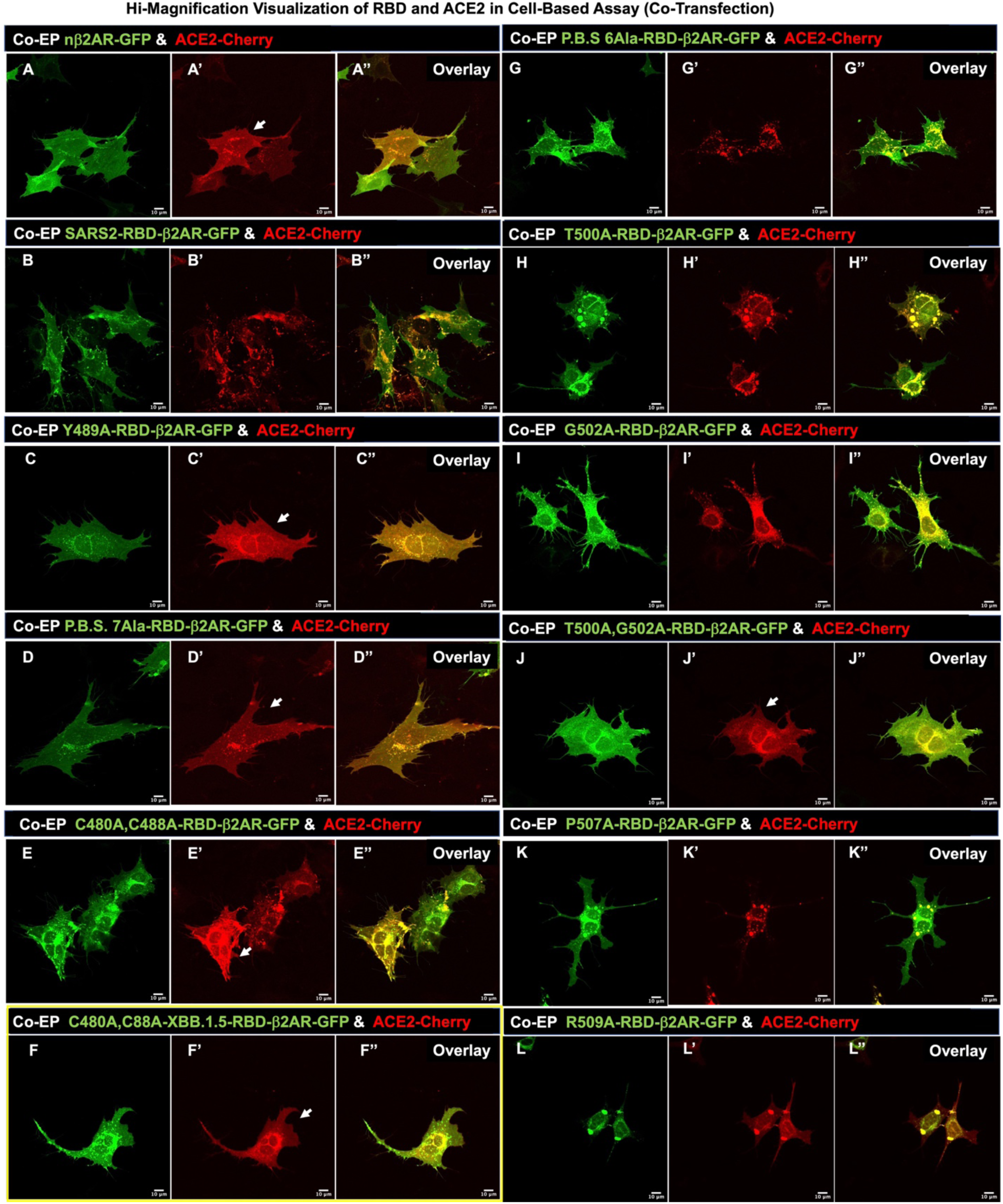
Co-EP with RBD mutations and ACE2. **A-L, A’-L’, A’’-L’’.** Utilizing LSM501, confocal images for Co-EP of GFP-tagged RBD variants and mutants and controls along with ACE2-Cherry, imaged after 16hrs. Green **(A-L)** and Red **(A’-L’)** channels have been split, and overlay (**A’’-L’’**). **A-A’’**. b2AR and ACE2 show uniform membrane expression. **B-B’’.** SARS2-RBD and ACE2 show internalization of all ACE2-Cherry fluorescence. **C-L, C’-L’, C’’-L’’.** Of the four assays (DPF assay; Cell-Dissociation assay; Co-plating assay; Co-EP RBD/ACE2 assay) that were set up, the Co-EP proved to be the most sensitive way to determine any latent binding between RBD and ACE2 proteins. **A’, C’, D’, F’, and J’** show no alteration of membrane ACE2 expression. **B’, G’, H’, K’, L’** show robust internalization of ACE2-Cherry. **E’, I’** show only modest internalization of ACE2-Cherry. It should be noted that C480A, C488A double mutant in XBB.1.5-RBD (**F-F’’**) abolishes ACE2 internalization more robustly in this assay than C480A, C488A double mutant in RBD (**E-E’’**), but not in DPF assay (**Figure 11F’**).

**Figure S15:**
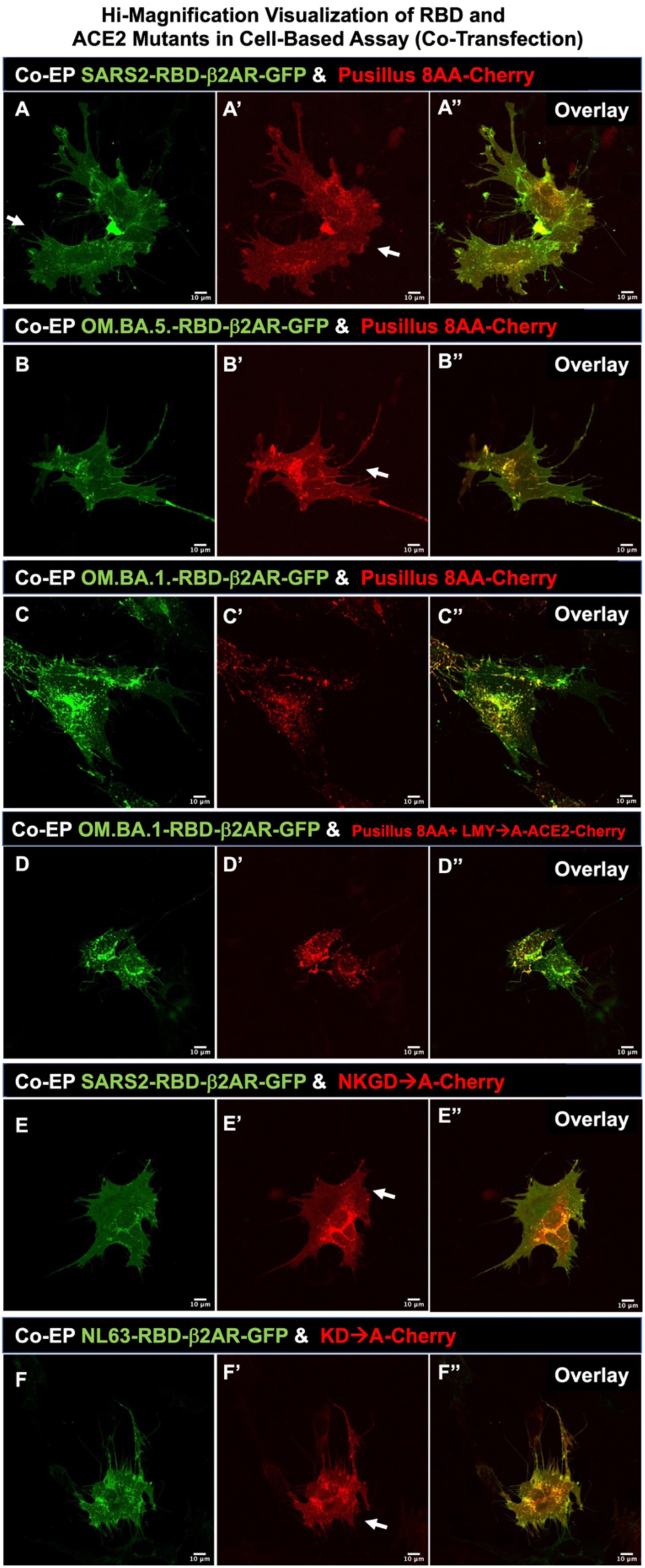
Co-EP with RBD variants and ACE2 mutants. **A-F, A’-F’, A’’-F’’.** Utilizing LSM510, confocal images for Co-EP of GFP-tagged RBD variants and mutants and controls along with ACE2-Cherry, imaged after 16hrs. Green **(A-F)** and Red **(A’-F’)** channels have been split, and overlay (**A’’-F’’**). Omicron.BA.1 clearly shows differential binding to Pusillus 8AA-Cherry compared to SARS2 and OM.BA.5-RBDs. OM.BA.1-RBD shows some internalization of ACE2-Cherry (**C’**), which shows the sensitivity of binding in this assay compared to co-plating assay in **Figure S14 A-F** and DPF assay in Figure 10. Both SARS2 and NL63-RBDs show dependence on Internal Region 2 for binding ACE2-Cherry with no alteration of membrane ACE2 expression (**E’, F’**).

**Figure S16:**
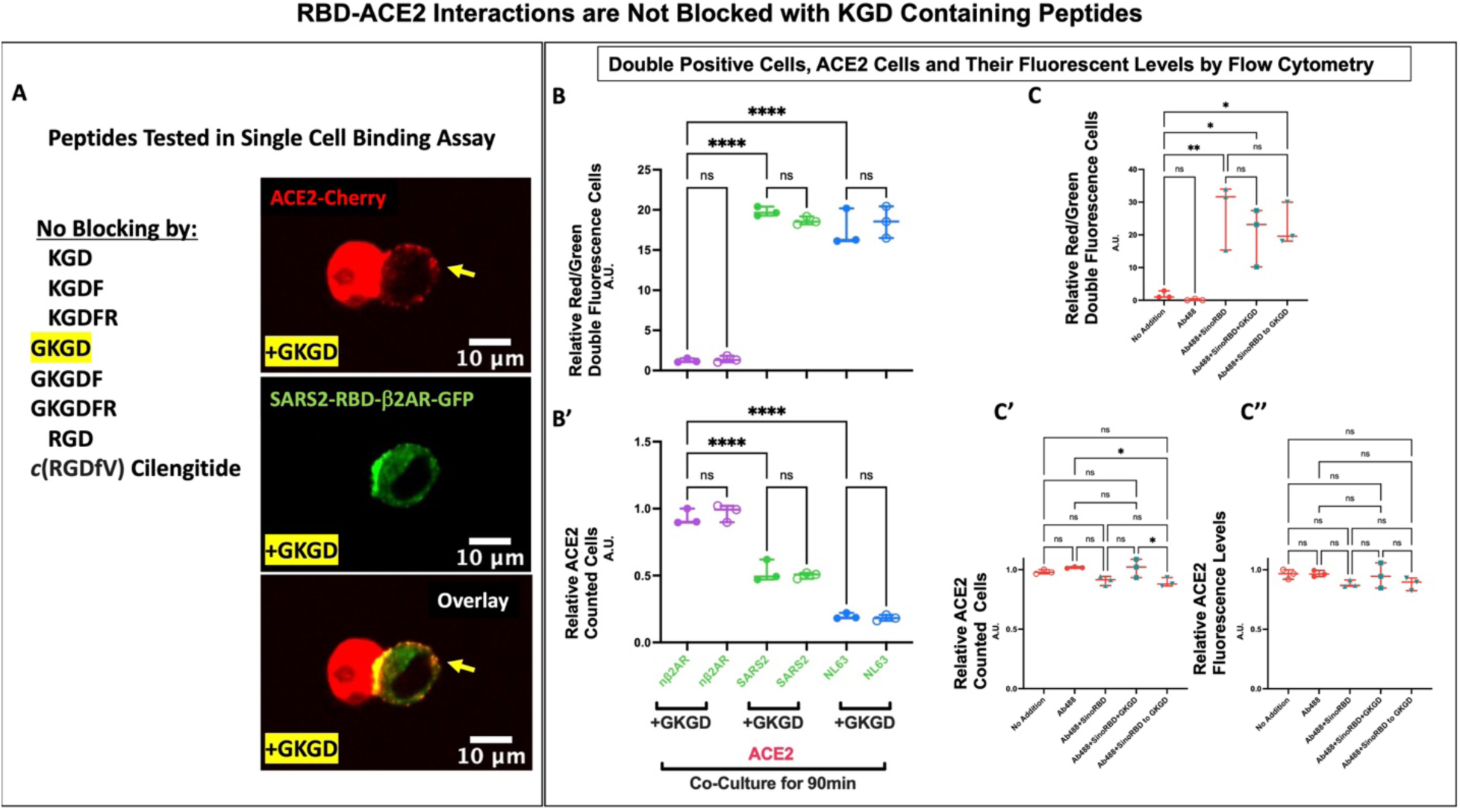
KGD containing peptides do not antagonize RBD binding to ACE2-Cherry. **A**. The necessity of K353-G354-D355 sequence in ACE2 suggested that KGD derivative peptides could block RBD binding. Six peptide versions of KGD and two RGD containing peptides were tested for their ability to block SARS2 or NL63 RBDs from binding ACE2-Cherryin the 30min dissociated cell-based assay. Greater than 100 micromolar concentrations for all 8 peptides did not block binding. Utilizing LSM510, confocal images revealed only one peptide GKGD (yellow highlight) appeared to cause a phenotype that was not previously observed: ACE2 protein being redistributed on the adjoining SARS2-RBD cells. **B and B’**. *DPF assay*: analysis of Co-plating GFP and Cherry expressing cells after 90min. Data shows that double florescent cells are found in both SARS2 and NL63 irrespective of preincubation with 120mM GKGD peptide with subsequent 90min incubation at a 1:20 dilution (6mM GKDG). Concomitant loss of ACE2-Cherry expression was observed in both ACE2 cells in all RBD conditions whether the peptide was present. **C-C’’**. *DPF assay*: **C**. Double fluorescence after 90minute incubation of ACE2-Cherry cells with Ab488 (1mg/4mls of Goat-anti-HIS-Tag-Alexa488 Goat antibody [R&D#: IC050G]) with and without SinoRBD (60ng/4mls of 2019-nCoV-HisTag-RBD [Cat#40592-V08H; 234 residues, R319-F541]). 1000x-GKGD peptide(49mg) 10min pre-incubated with 1mg Ab488 + 60ng SinoRBD in 200mls PBS, then added to 4mls of media or GKDG added to media (49mg/4mls) plus 200mls PBS with 10min preincubated of 1ug Ab488 + 60ng SinoRBD. No reduction of double fluorescence is observed when GKGD peptide is added. **C’** ACE2-Cherry cells with Ab488, Ab488+SinoRBD, or GKGD+Ab488+SinoRBD show no changes in fluorescence. **C’’**. High expressing ACE2-Cherry fluorescence cells also show no changes in fluorescence in all conditions. Values in **B, B’, C, C’and C’’** are normalized to controls (Y-axis arbitrary units, A. U.)-**see Data File S2**. GraphPad Prism 9.3.1 used to generate **B-C’’**.

**Figure S17:**
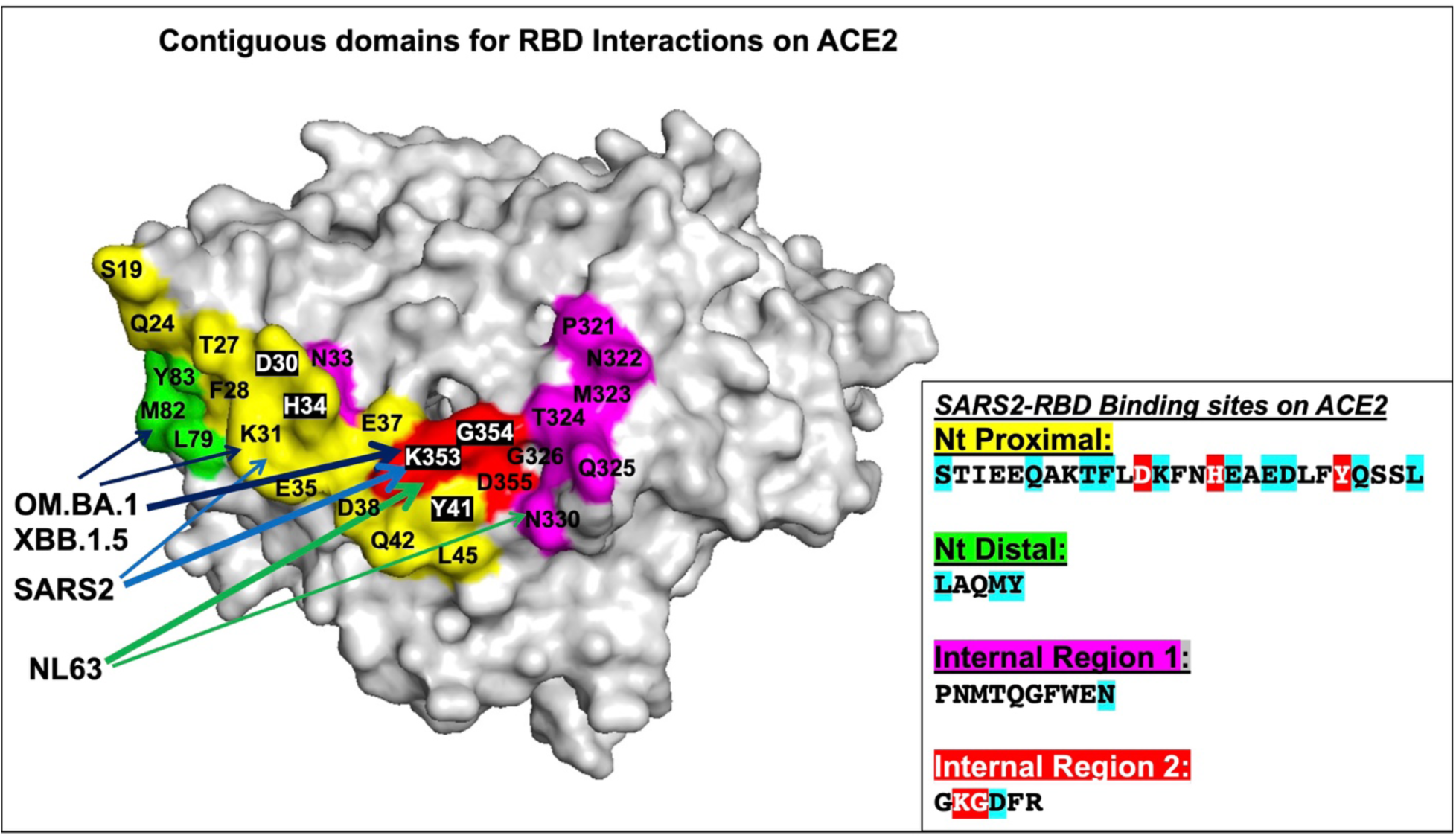
Proposed interpretation of RBD interactions on ACE2 structure with demarcated domains. Space-filling model of ACE2 structure from pdb_6M0J (14) modified by highlighting interactions for both SARS2 and NL63 RBDs: Nt proximal (**yellow residues**), Nt distal (**green residues**), and Internal regions 1 (**purple residues**) and 2 (**red residues**). Purple residue interactions including N33 are nearly from NL63-RBD. SARS2-RBD does not bind/internalize if *either* Nt proximal or Internal region 2 (K353, D355) are mutated. OM.BA.1 and OM.BA2-RBDs are not bond/internalized if *either* mutation in both Nt proximal and distal ACE2 or Internal region 2 (K353, D355). NL63-RBD requires Internal region 2 (K353 and D355) for both binding/internalization. Inset shows **SARS2-RBD** interactions within the four domains (**see Figures 7, 12**) (residues with blue highlights are predicted binding sites that occur in ¾ of SARS2, SARS1, RaTG13, and NL63-RBDs; residues with red highlights are predicted binding sites in all 4 RBDs). The RBD interaction data taken together suggests that Internal region 2 binding for SARS1/SARS2/RaTG13 is weak and requires Nt proximal and distal contacts for stabilization (dark blue, light blue arrows), whereas NL63 has additional contacts in the Internal region 1 (green arrows) that supports binding to Internal region 2 (thick arrows). The ability of SARS2 mutant Y489A to nearly abolish SARS2-RBD binding to ACE2 suggests that none of the domains by themselves have strong binding to ACE2. Overall, the data points to the Internal region 2 as being critical for internalizations of RBDs.

## References

1. Wang Q, Zhang Y, Wu L, Niu S, Song C, Zhang Z, Lu G, Qiao C, Hu Y, Yuen KY, Wang Q, Zhou H, Yan J, Qi J. Structural and Functional Basis of SARS-CoV-2 Entry by Using Human ACE2. Cell. 2020;181(4):894–904 e9. Epub 2020/04/11. doi: 10.1016/j.cell.2020.03.045. PubMed PMID: 32275855; PMCID: PMC7144619.

2. Perlman KMaS. Coronaviruses, Including Severe Acute Respiratory Syndrome (SARS) and Middle East Respiratory Syndrome (MERS). Mandell, Douglas, and Bennett’s Principles and Practice of Infectious Diseases 2015. 2015(Chapter 157):1928–36.e2. Epub 2014 Oct 31. doi: 10.1016/B978-1-4557-4801-3.00157-0; PMCID: PMC7151770.

3. Borah P, Deb PK, Al-Shar’i NA, Dahabiyeh LA, Venugopala KN, Singh V, Shinu P, Hussain S, Deka S, Chandrasekaran B, Jaradat DMM. Perspectives on RNA Vaccine Candidates for COVID-19. Front Mol Biosci. 2021;8:635245. Epub 2021/04/20. doi: 10.3389/fmolb.2021.635245. PubMed PMID: 33869282; PMCID: PMC8044912.

4. Park JW, Lagniton PNP, Liu Y, Xu RH. mRNA vaccines for COVID-19: what, why and how. Int J Biol Sci. 2021;17(6):1446–60. Epub 2021/04/29. doi: 10.7150/ijbs.59233. PubMed PMID: 33907508; PMCID: PMC8071766.

5. Drozdzal S, Rosik J, Lechowicz K, Machaj F, Szostak B, Przybycinski J, Lorzadeh S, Kotfis K, Ghavami S, Los MJ. An update on drugs with therapeutic potential for SARS-CoV-2 (COVID-19) treatment. Drug Resist Updat. 2021;59:100794. Epub 2022/01/08. doi: 10.1016/j.drup.2021.100794. PubMed PMID: 34991982; PMCID: PMC8654464.

6. Killingley B, Mann AJ, Kalinova M, Boyers A, Goonawardane N, Zhou J, Lindsell K, Hare SS, Brown J, Frise R, Smith E, Hopkins C, Noulin N, Londt B, Wilkinson T, Harden S, McShane H, Baillet M, Gilbert A, Jacobs M, Charman C, Mande P, Nguyen-Van-Tam JS, Semple MG, Read RC, Ferguson NM, Openshaw PJ, Rapeport G, Barclay WS, Catchpole AP, Chiu C. Safety, tolerability and viral kinetics during SARS-CoV-2 human challenge in young adults. Nat Med. 2022;28(5):1031–41. Epub 2022/04/02. doi: 10.1038/s41591-022-01780-9. PubMed PMID: 35361992.

7. Shapira T, Monreal IA, Dion SP, Buchholz DW, Imbiakha B, Olmstead AD, Jager M, Desilets A, Gao G, Martins M, Vandal T, Thompson CAH, Chin A, Rees WD, Steiner T, Nabi IR, Marsault E, Sahler J, Diel DG, Van de Walle GR, August A, Whittaker GR, Boudreault PL, Leduc R, Aguilar HC, Jean F. A TMPRSS2 inhibitor acts as a pan-SARS-CoV-2 prophylactic and therapeutic. Nature. 2022;605(7909):340–8. Epub 2022/03/29. doi: 10.1038/s41586-022-04661-w. PubMed PMID: 35344983; PMCID: PMC9095466 US10988505B2) that cover matriptase and other type II transmembrane serine proteases inhibitors for treating and preventing viral infections, respiratory disorders, inflammatory disorders, pain disorders, tissue disorders, hyperproliferative disorders, and disorders associated with iron overload. The remaining authors declare that they have no competing interests.

8. Hoffmann M, Kleine-Weber H, Schroeder S, Kruger N, Herrler T, Erichsen S, Schiergens TS, Herrler G, Wu NH, Nitsche A, Muller MA, Drosten C, Pohlmann S. SARS-CoV-2 Cell Entry Depends on ACE2 and TMPRSS2 and Is Blocked by a Clinically Proven Protease Inhibitor. Cell. 2020;181(2):271–80 e8. Epub 2020/03/07. doi: 10.1016/j.cell.2020.02.052. PubMed PMID: 32142651; PMCID: PMC7102627.

9. Han DP, Penn-Nicholson A, Cho MW. Identification of critical determinants on ACE2 for SARS-CoV entry and development of a potent entry inhibitor. Virology. 2006;350(1):15–25. Epub 2006/03/03. doi: 10.1016/j.virol.2006.01.029. PubMed PMID: 16510163; PMCID: PMC7111894.

10. Hashemian SMR, Sheida A, Taghizadieh M, Memar MY, Hamblin MR, Bannazadeh Baghi H, Sadri Nahand J, Asemi Z, Mirzaei H. Paxlovid (Nirmatrelvir/Ritonavir): A new approach to Covid-19 therapy? Biomed Pharmacother. 2023;162:114367. Epub 2023/04/06. doi: 10.1016/j.biopha.2023.114367. PubMed PMID: 37018987; PMCID: PMC9899776.

11. Jonsdottir HR, Dijkman R. Coronaviruses and the human airway: a universal system for virus-host interaction studies. Virol J. 2016;13:24. Epub 2016/02/08. doi: 10.1186/s12985-016-0479-5. PubMed PMID: 26852031; PMCID: PMC4744394.

12. Leon Fodoulian JT, Daniel Rossier, Madlaina Boillat, Chenda Kan, Véronique Pauli, Kristof Egervari, Johannes A. Lobrinus, Basile N. Landis, Alan Carleton, Ivan Rodriguez. SARS-CoV-2 receptor and entry genes are expressed by sustentacular cells in the human olfactory neuroepithelium. bioRxiv 20200331013268; doi: https://doiorg/101101/20200331013268. 2020.

13. David H. Brann TT, Caleb Weinreb, Marcela Lipovsek, Koen Van den Berge, Boying Gong, Rebecca Chance, Iain C. Macaulay, Hsin-jung Chou, Russell Fletcher, Diya Das, Kelly Street, Hector Roux de Bezieux, Yoon-Gi Choi, Davide Risso, Sandrine Dudoit, Elizabeth Purdom, Jonathan S. Mill, Ralph Abi Hachem, Hiroaki Matsunami, Darren W. Logan, Bradley J. Goldstein, Matthew S. Grubb, John Ngai, Sandeep Robert Datta. Non-neuronal expression of SARS-CoV-2 entry genes in the olfactory system suggests mechanisms underlying COVID-19-associated anosmia. bioRxiv 20200325009084; doi: https://doiorg/101101/20200325009084. 2020.

14. Iwata-Yoshikawa N, Kakizaki M, Shiwa-Sudo N, Okura T, Tahara M, Fukushi S, Maeda K, Kawase M, Asanuma H, Tomita Y, Takayama I, Matsuyama S, Shirato K, Suzuki T, Nagata N, Takeda M. Essential role of TMPRSS2 in SARS-CoV-2 infection in murine airways. Nat Commun. 2022;13(1):6100. Epub 2022/10/16. doi: 10.1038/s41467-022-33911-8. PubMed PMID: 36243815; PMCID: PMC9568946.

15. Beaudoin CA, Pandurangan AP, Kim SY, Hamaia SW, Huang CL, Blundell TL, Vedithi SC, Jackson AP. In silico analysis of mutations near S1/S2 cleavage site in SARS-CoV-2 spike protein reveals increased propensity of glycosylation in Omicron strain. J Med Virol. 2022;94(9):4181–92. Epub 2022/05/17. doi: 10.1002/jmv.27845. PubMed PMID: 35575289; PMCID: PMC9348480.

16. Koch J, Uckeley ZM, Doldan P, Stanifer M, Boulant S, Lozach PY. TMPRSS2 expression dictates the entry route used by SARS-CoV-2 to infect host cells. Embo J. 2021;40(16):e107821. Epub 2021/06/24. doi: 10.15252/embj.2021107821. PubMed PMID: 34159616; PMCID: PMC8365257.

17. Brann DH, Tsukahara T, Weinreb C, Lipovsek M, Van den Berge K, Gong B, Chance R, Macaulay IC, Chou HJ, Fletcher RB, Das D, Street K, de Bezieux HR, Choi YG, Risso D, Dudoit S, Purdom E, Mill J, Hachem RA, Matsunami H, Logan DW, Goldstein BJ, Grubb MS, Ngai J, Datta SR. Non-neuronal expression of SARS-CoV-2 entry genes in the olfactory system suggests mechanisms underlying COVID-19-associated anosmia. Sci Adv. 2020;6(31). Epub 2020/09/17. doi: 10.1126/sciadv.abc5801. PubMed PMID: 32937591.

18. Fodoulian L, Tuberosa J, Rossier D, Boillat M, Kan C, Pauli V, Egervari K, Lobrinus JA, Landis BN, Carleton A, Rodriguez I. SARS-CoV-2 Receptors and Entry Genes Are Expressed in the Human Olfactory Neuroepithelium and Brain. iScience. 2020;23(12):101839. Epub 2020/12/01. doi: 10.1016/j.isci.2020.101839. PubMed PMID: 33251489; PMCID: PMC7685946.

19. Zazhytska M, Kodra A, Hoagland DA, Frere J, Fullard JF, Shayya H, McArthur NG, Moeller R, Uhl S, Omer AD, Gottesman ME, Firestein S, Gong Q, Canoll PD, Goldman JE, Roussos P, tenOever BR, Jonathan BO, Lomvardas S. Non-cell-autonomous disruption of nuclear architecture as a potential cause of COVID-19-induced anosmia. Cell. 2022;185(6):1052–64 e12. Epub 2022/02/19. doi: 10.1016/j.cell.2022.01.024. PubMed PMID: 35180380; PMCID: PMC8808699.

20. Bubnell J, Pfister P, Sapar ML, Rogers ME, Feinstein P. beta2 adrenergic receptor fluorescent protein fusions traffic to the plasma membrane and retain functionality. PLoS One. 2013;8(9):e74941. Epub 2013/10/03. doi: 10.1371/journal.pone.0074941. PubMed PMID: 24086401; PMCID: PMC3781101.

21. Bubnell J, Jamet S, Tomoiaga D, D’Hulst C, Krampis K, Feinstein P. In Vitro Mutational and Bioinformatics Analysis of the M71 Odorant Receptor and Its Superfamily. PLoS One. 2015;10(10):e0141712. Epub 2015/10/30. doi: 10.1371/journal.pone.0141712. PubMed PMID: 26513476; PMCID: PMC4626375.

22. Jamet S, Bubnell J, Pfister P, Tomoiaga D, Rogers ME, Feinstein P. In Vitro Mutational Analysis of the beta2 Adrenergic Receptor, an In Vivo Surrogate Odorant Receptor. PLoS One. 2015;10(10):e0141696. Epub 2015/10/30. doi: 10.1371/journal.pone.0141696. PubMed PMID: 26513247; PMCID: PMC4626089.

23. Wong SK, Li W, Moore MJ, Choe H, Farzan M. A 193-amino acid fragment of the SARS coronavirus S protein efficiently binds angiotensin-converting enzyme 2. J Biol Chem. 2004;279(5):3197–201. Epub 2003/12/13. doi: 10.1074/jbc.C300520200. PubMed PMID: 14670965; PMCID: PMC7982343.

24. Towler P, Staker B, Prasad SG, Menon S, Tang J, Parsons T, Ryan D, Fisher M, Williams D, Dales NA, Patane MA, Pantoliano MW. ACE2 X-ray structures reveal a large hinge-bending motion important for inhibitor binding and catalysis. J Biol Chem. 2004;279(17):17996–8007. Epub 2004/02/03. doi: 10.1074/jbc.M311191200. PubMed PMID: 14754895; PMCID: PMC7980034.

25. Chan KK, Dorosky D, Sharma P, Abbasi SA, Dye JM, Kranz DM, Herbert AS, Procko E. Engineering human ACE2 to optimize binding to the spike protein of SARS coronavirus 2. Science. 2020;369(6508):1261–5. Epub 2020/08/06. doi: 10.1126/science.abc0870. PubMed PMID: 32753553.

26. Liu K, Pan X, Li L, Yu F, Zheng A, Du P, Han P, Meng Y, Zhang Y, Wu L, Chen Q, Song C, Jia Y, Niu S, Lu D, Qiao C, Chen Z, Ma D, Ma X, Tan S, Zhao X, Qi J, Gao GF, Wang Q. Binding and molecular basis of the bat coronavirus RaTG13 virus to ACE2 in humans and other species. Cell. 2021;184(13):3438–51 e10. Epub 2021/06/18. doi: 10.1016/j.cell.2021.05.031. PubMed PMID: 34139177; PMCID: PMC8142884.

27. Wu K, Li W, Peng G, Li F. Crystal structure of NL63 respiratory coronavirus receptor-binding domain complexed with its human receptor. Proc Natl Acad Sci U S A. 2009;106(47):19970–4. Epub 2009/11/11. doi: 10.1073/pnas.0908837106. PubMed PMID: 19901337; PMCID: PMC2785276.

28. Lan J, Ge J, Yu J, Shan S, Zhou H, Fan S, Zhang Q, Shi X, Wang Q, Zhang L, Wang X. Structure of the SARS-CoV-2 spike receptor-binding domain bound to the ACE2 receptor. Nature. 2020;581(7807):215–20. Epub 2020/04/01. doi: 10.1038/s41586-020-2180-5. PubMed PMID: 32225176.

29. Grishin AM, Dolgova NV, Landreth S, Fisette O, Pickering IJ, George GN, Falzarano D, Cygler M. Disulfide Bonds Play a Critical Role in the Structure and Function of the Receptor-binding Domain of the SARS-CoV-2 Spike Antigen. J Mol Biol. 2022;434(2):167357. Epub 2021/11/16. doi: 10.1016/j.jmb.2021.167357. PubMed PMID: 34780781; PMCID: PMC8588607.

30. Khazaal S, Harb J, Rima M, Annweiler C, Wu Y, Cao Z, Abi Khattar Z, Legros C, Kovacic H, Fajloun Z, Sabatier JM. The Pathophysiology of Long COVID throughout the Renin-Angiotensin System. Molecules. 2022;27(9). Epub 2022/05/15. doi: 10.3390/molecules27092903. PubMed PMID: 35566253; PMCID: PMC9101946.

31. Keck T, Leiacker R, Riechelmann H, Rettinger G. Temperature profile in the nasal cavity. Laryngoscope. 2000;110(4):651–4. Epub 2000/04/14. doi: 10.1097/00005537-200004000-00021. PubMed PMID: 10764013.

32. Jeanneteau F, Diaz J, Sokoloff P, Griffon N. Interactions of GIPC with dopamine D2, D3 but not D4 receptors define a novel mode of regulation of G protein-coupled receptors. Mol Biol Cell. 2004;15(2):696-705. Epub 2003/11/18. doi: 10.1091/mbc.e03-05-0293. PubMed PMID: 14617818; PMCID: PMC329290.

33. Weill C, Ilien B, Goeldner M, Galzi JL. Fluorescent muscarinic EGFP-hM1 chimeric receptors: design, ligand binding and functional properties. J Recept Signal Transduct Res. 1999;19(1-4):423–36. Epub 1999/03/11. doi: 10.3109/10799899909036662. PubMed PMID: 10071775.

34. Parvanova I. Getting at the Surface: A Promoter and Coding Sequence Characterization of an Odorant Receptor. CUNY Academic Works. 2019.

35. McMahon C, Scadding GK. Le Nez du Vin--a quick test of olfaction. Clin Otolaryngol Allied Sci. 1996;21(3):278–80. Epub 1996/06/01. doi: 10.1111/j.1365-2273.1996.tb01741.x. PubMed PMID: 8818503.

36. Lu Y, Zhu Q, Fox DM, Gao C, Stanley SA, Luo K. SARS-CoV-2 down-regulates ACE2 through lysosomal degradation. Mol Biol Cell. 2022;33(14):ar147. Epub 2022/10/27. doi: 10.1091/mbc.E22-02-0045. PubMed PMID: 36287912; PMCID: PMC9727799.

37. Shang C, Zhuang X, Zhang H, Li Y, Zhu Y, Lu J, Ge C, Cong J, Li T, Tian M, Jin N, Li X. Inhibitors of endosomal acidification suppress SARS-CoV-2 replication and relieve viral pneumonia in hACE2 transgenic mice. Virol J. 2021;18(1):46. Epub 2021/03/01. doi: 10.1186/s12985-021-01515-1. PubMed PMID: 33639976; PMCID: PMC7914043.

38. Jackson CB, Farzan M, Chen B, Choe H. Mechanisms of SARS-CoV-2 entry into cells. Nat Rev Mol Cell Biol. 2022;23(1):3–20. Epub 2021/10/07. doi: 10.1038/s41580-021-00418-x. PubMed PMID: 34611326; PMCID: PMC8491763.

39. Szewczyk-Roszczenko OK, Roszczenko P, Shmakova A, Finiuk N, Holota S, Lesyk R, Bielawska A, Vassetzky Y, Bielawski K. The Chemical Inhibitors of Endocytosis: From Mechanisms to Potential Clinical Applications. Cells. 2023;12(18). Epub 2023/09/28. doi: 10.3390/cells12182312. PubMed PMID: 37759535; PMCID: PMC10527932.

40. Bayati A, Kumar R, Francis V, McPherson PS. SARS-CoV-2 infects cells after viral entry via clathrin-mediated endocytosis. J Biol Chem. 2021;296:100306. Epub 2021/01/22. doi: 10.1016/j.jbc.2021.100306. PubMed PMID: 33476648; PMCID: PMC7816624.

41. Starr TN, Greaney AJ, Hilton SK, Ellis D, Crawford KHD, Dingens AS, Navarro MJ, Bowen JE, Tortorici MA, Walls AC, King NP, Veesler D, Bloom JD. Deep Mutational Scanning of SARS-CoV-2 Receptor Binding Domain Reveals Constraints on Folding and ACE2 Binding. Cell. 2020;182(5):1295–310 e20. Epub 2020/08/26. doi: 10.1016/j.cell.2020.08.012. PubMed PMID: 32841599; PMCID: PMC7418704.

42. Jawad B, Adhikari P, Podgornik R, Ching WY. Key Interacting Residues between RBD of SARS-CoV-2 and ACE2 Receptor: Combination of Molecular Dynamics Simulation and Density Functional Calculation. J Chem Inf Model. 2021;61(9):4425–41. Epub 2021/08/25. doi: 10.1021/acs.jcim.1c00560. PubMed PMID: 34428371; PMCID: PMC8409146.

43. Harvey WT, Carabelli AM, Jackson B, Gupta RK, Thomson EC, Harrison EM, Ludden C, Reeve R, Rambaut A, Consortium C-GU, Peacock SJ, Robertson DL. SARS-CoV-2 variants, spike mutations and immune escape. Nat Rev Microbiol. 2021;19(7):409–24. Epub 2021/06/03. doi: 10.1038/s41579-021-00573-0. PubMed PMID: 34075212; PMCID: PMC8167834.

44. Li W, Sui J, Huang IC, Kuhn JH, Radoshitzky SR, Marasco WA, Choe H, Farzan M. The S proteins of human coronavirus NL63 and severe acute respiratory syndrome coronavirus bind overlapping regions of ACE2. Virology. 2007;367(2):367–74. Epub 2007/07/17. doi: 10.1016/j.virol.2007.04.035. PubMed PMID: 17631932; PMCID: PMC2693060.

45. Zhou H, Ji J, Chen X, Bi Y, Li J, Wang Q, Hu T, Song H, Zhao R, Chen Y, Cui M, Zhang Y, Hughes AC, Holmes EC, Shi W. Identification of novel bat coronaviruses sheds light on the evolutionary origins of SARS-CoV-2 and related viruses. Cell. 2021;184(17):4380–91 e14. Epub 2021/06/21. doi: 10.1016/j.cell.2021.06.008. PubMed PMID: 34147139; PMCID: PMC8188299.

46. Temmam S, Vongphayloth K, Baquero E, Munier S, Bonomi M, Regnault B, Douangboubpha B, Karami Y, Chretien D, Sanamxay D, Xayaphet V, Paphaphanh P, Lacoste V, Somlor S, Lakeomany K, Phommavanh N, Perot P, Dehan O, Amara F, Donati F, Bigot T, Nilges M, Rey FA, van der Werf S, Brey PT, Eloit M. Bat coronaviruses related to SARS-CoV-2 and infectious for human cells. Nature. 2022;604(7905):330–6. Epub 2022/02/17. doi: 10.1038/s41586-022-04532-4. PubMed PMID: 35172323.

47. Lin HX, Feng Y, Tu X, Zhao X, Hsieh CH, Griffin L, Junop M, Zhang C. Characterization of the spike protein of human coronavirus NL63 in receptor binding and pseudotype virus entry. Virus Res. 2011;160(1-2):283–93. Epub 2011/07/30. doi: 10.1016/j.virusres.2011.06.029. PubMed PMID: 21798295; PMCID: PMC7114368.

48. Wu K, Chen L, Peng G, Zhou W, Pennell CA, Mansky LM, Geraghty RJ, Li F. A virus-binding hot spot on human angiotensin-converting enzyme 2 is critical for binding of two different coronaviruses. J Virol. 2011;85(11):5331–7. Epub 2011/03/18. doi: 10.1128/JVI.02274-10. PubMed PMID: 21411533; PMCID: PMC3094985.

49. Kapp TG, Rechenmacher F, Neubauer S, Maltsev OV, Cavalcanti-Adam EA, Zarka R, Reuning U, Notni J, Wester HJ, Mas-Moruno C, Spatz J, Geiger B, Kessler H. A Comprehensive Evaluation of the Activity and Selectivity Profile of Ligands for RGD-binding Integrins. Sci Rep. 2017;7:39805. Epub 2017/01/12. doi: 10.1038/srep39805. PubMed PMID: 28074920; PMCID: PMC5225454.

50. Mancek-Keber M, Hafner-Bratkovic I, Lainscek D, Bencina M, Govednik T, Orehek S, Plaper T, Jazbec V, Bergant V, Grass V, Pichlmair A, Jerala R. Disruption of disulfides within RBD of SARS-CoV-2 spike protein prevents fusion and represents a target for viral entry inhibition by registered drugs. FASEB J. 2021;35(6):e21651. Epub 2021/05/19. doi: 10.1096/fj.202100560R. PubMed PMID: 34004056; PMCID: PMC8206760.

51. Lukesh JC, 3rd, Palte MJ, Raines RT. A potent, versatile disulfide-reducing agent from aspartic acid. J Am Chem Soc. 2012;134(9):4057–9. Epub 2012/02/23. doi: 10.1021/ja211931f. PubMed PMID: 22353145; PMCID: PMC3353773.

52. Damas J, Hughes GM, Keough KC, Painter CA, Persky NS, Corbo M, Hiller M, Koepfli KP, Pfenning AR, Zhao H, Genereux DP, Swofford R, Pollard KS, Ryder OA, Nweeia MT, Lindblad-Toh K, Teeling EC, Karlsson EK, Lewin HA. Broad host range of SARS-CoV-2 predicted by comparative and structural analysis of ACE2 in vertebrates. Proc Natl Acad Sci U S A. 2020;117(36):22311–22. Epub 2020/08/23. doi: 10.1073/pnas.2010146117. PubMed PMID: 32826334; PMCID: PMC7486773.

53. Mou H, Quinlan BD, Peng H, Liu G, Guo Y, Peng S, Zhang L, Davis-Gardner ME, Gardner MR, Crynen G, DeVaux LB, Voo ZX, Bailey CC, Alpert MD, Rader C, Gack MU, Choe H, Farzan M. Mutations derived from horseshoe bat ACE2 orthologs enhance ACE2-Fc neutralization of SARS-CoV-2. PLoS Pathog. 2021;17(4):e1009501. Epub 2021/04/10. doi: 10.1371/journal.ppat.1009501. PubMed PMID: 33836016; PMCID: PMC8059821 following competing interests: M.F., H.M. and B.D.Q had filed a patent for the application of ACE2-Fc variants as a SARS-CoV-2 treatment. M.R.G., C.C.B., M.D.A., and M.F. are all cofounders of, and have an equity interest in Emmune Inc., a biotech company that specializes in the development of antibody-like antiviral therapies.

54. Conceicao C, Thakur N, Human S, Kelly JT, Logan L, Bialy D, Bhat S, Stevenson-Leggett P, Zagrajek AK, Hollinghurst P, Varga M, Tsirigoti C, Tully M, Chiu C, Moffat K, Silesian AP, Hammond JA, Maier HJ, Bickerton E, Shelton H, Dietrich I, Graham SC, Bailey D. The SARS-CoV-2 Spike protein has a broad tropism for mammalian ACE2 proteins. PLoS Biol. 2020;18(12):e3001016. Epub 2020/12/22. doi: 10.1371/journal.pbio.3001016. PubMed PMID: 33347434; PMCID: PMC7751883.

55. Li Y, Wang H, Tang X, Fang S, Ma D, Du C, Wang Y, Pan H, Yao W, Zhang R, Zou X, Zheng J, Xu L, Farzan M, Zhong G. SARS-CoV-2 and Three Related Coronaviruses Utilize Multiple ACE2 Orthologs and Are Potently Blocked by an Improved ACE2-Ig. J Virol. 2020;94(22). Epub 2020/08/28. doi: 10.1128/JVI.01283-20. PubMed PMID: 32847856; PMCID: PMC7592233.

56. Liu X, Garriga P, Khorana HG. Structure and function in rhodopsin: correct folding and misfolding in two point mutants in the intradiscal domain of rhodopsin identified in retinitis pigmentosa. Proc Natl Acad Sci U S A. 1996;93(10):4554–9. Epub 1996/05/14. doi: 10.1073/pnas.93.10.4554. PubMed PMID: 8643442; PMCID: PMC39315.

57. Choi CY, Gadhave K, Villano J, Pekosz A, Mao X, Jia H. Generation and characterization of a humanized ACE2 mouse model to study long-term impacts of SARS-CoV-2 infection. J Med Virol. 2024;96(1):e29349. Epub 2024/01/08. doi: 10.1002/jmv.29349. PubMed PMID: 38185937; PMCID: PMC10783855.

58. Cagan RL, Kramer H, Hart AC, Zipursky SL. The bride of sevenless and sevenless interaction: internalization of a transmembrane ligand. Cell. 1992;69(3):393–9. Epub 1992/05/01. doi: 10.1016/0092-8674(92)90442-f. PubMed PMID: 1316239.

59. Langridge PD, Struhl G. Epsin-Dependent Ligand Endocytosis Activates Notch by Force. Cell. 2017;171(6):1383–96 e12. Epub 2017/12/02. doi: 10.1016/j.cell.2017.10.048. PubMed PMID: 29195077; PMCID: PMC6219616.

60. Mathewson AC, Bishop A, Yao Y, Kemp F, Ren J, Chen H, Xu X, Berkhout B, van der Hoek L, Jones IM. Interaction of severe acute respiratory syndrome-coronavirus and NL63 coronavirus spike proteins with angiotensin converting enzyme-2. J Gen Virol. 2008;89(Pt 11):2741–5. Epub 2008/10/22. doi: 10.1099/vir.0.2008/003962-0. PubMed PMID: 18931070; PMCID: PMC2886958.

61. Brown EEF, Rezaei R, Jamieson TR, Dave J, Martin NT, Singaravelu R, Crupi MJF, Boulton S, Tucker S, Duong J, Poutou J, Pelin A, Yasavoli-Sharahi H, Taha Z, Arulanandam R, Surendran A, Ghahremani M, Austin B, Matar C, Diallo JS, Bell JC, Ilkow CS, Azad T. Characterization of Critical Determinants of ACE2-SARS CoV-2 RBD Interaction. Int J Mol Sci. 2021;22(5). Epub 2021/03/07. doi: 10.3390/ijms22052268. PubMed PMID: 33668756; PMCID: PMC7956771.

62. Chan KK, Tan TJC, Narayanan KK, Procko E. An engineered decoy receptor for SARS-CoV-2 broadly binds protein S sequence variants. Sci Adv. 2021;7(8). Epub 2021/02/19. doi: 10.1126/sciadv.abf1738. PubMed PMID: 33597251; PMCID: PMC7888922.

63. Butowt R, Bilinska K, von Bartheld CS. Olfactory dysfunction in COVID-19: new insights into the underlying mechanisms. Trends Neurosci. 2023;46(1):75–90. Epub 2022/12/06. doi: 10.1016/j.tins.2022.11.003. PubMed PMID: 36470705; PMCID: PMC9666374.

64. Essalmani R, Jain J, Susan-Resiga D, Andreo U, Evagelidis A, Derbali RM, Huynh DN, Dallaire F, Laporte M, Delpal A, Sutto-Ortiz P, Coutard B, Mapa C, Wilcoxen K, Decroly E, Nq Pham T, Cohen EA, Seidah NG. Distinctive Roles of Furin and TMPRSS2 in SARS-CoV-2 Infectivity. J Virol. 2022;96(8):e0012822. Epub 2022/03/29. doi: 10.1128/jvi.00128-22. PubMed PMID: 35343766; PMCID: PMC9044946.

65. Papa G, Mallery DL, Albecka A, Welch LG, Cattin-Ortola J, Luptak J, Paul D, McMahon HT, Goodfellow IG, Carter A, Munro S, James LC. Furin cleavage of SARS-CoV-2 Spike promotes but is not essential for infection and cell-cell fusion. PLoS Pathog. 2021;17(1):e1009246. Epub 2021/01/26. doi: 10.1371/journal.ppat.1009246. PubMed PMID: 33493182; PMCID: PMC7861537.

66. Peacock TP, Goldhill DH, Zhou J, Baillon L, Frise R, Swann OC, Kugathasan R, Penn R, Brown JC, Sanchez-David RY, Braga L, Williamson MK, Hassard JA, Staller E, Hanley B, Osborn M, Giacca M, Davidson AD, Matthews DA, Barclay WS. The furin cleavage site in the SARS-CoV-2 spike protein is required for transmission in ferrets. Nat Microbiol. 2021;6(7):899–909. Epub 2021/04/29. doi: 10.1038/s41564-021-00908-w. PubMed PMID: 33907312.

67. Yu S, Zheng X, Zhou B, Li J, Chen M, Deng R, Wong G, Lavillette D, Meng G. SARS-CoV-2 spike engagement of ACE2 primes S2’ site cleavage and fusion initiation. Proc Natl Acad Sci U S A. 2022;119(1). Epub 2021/12/22. doi: 10.1073/pnas.2111199119. PubMed PMID: 34930824; PMCID: PMC8740742.

68. Rawat P, Jemimah S, Ponnuswamy PK, Gromiha MM. Why are ACE2 binding coronavirus strains SARS-CoV/SARS-CoV-2 wild and NL63 mild? Proteins. 2021;89(4):389–98. Epub 2020/11/20. doi: 10.1002/prot.26024. PubMed PMID: 33210300; PMCID: PMC7753379.

69. Tang AT, Buchholz DW, Szigety KM, Imbiakha B, Gao S, Frankfurter M, Wang M, Yang J, Hewins P, Mericko-Ishizuka P, Leu NA, Sterling S, Monreal IA, Sahler J, August A, Zhu X, Jurado KA, Xu M, Morrisey EE, Millar SE, Aguilar HC, Kahn ML. Cell-autonomous requirement for ACE2 across organs in lethal mouse SARS-CoV-2 infection. PLoS Biol. 2023;21(2):e3001989. Epub 2023/02/07. doi: 10.1371/journal.pbio.3001989. PubMed PMID: 36745682; PMCID: PMC9934376.

70. Astuti I, Ysrafil. Severe Acute Respiratory Syndrome Coronavirus 2 (SARS-CoV-2): An overview of viral structure and host response. Diabetes Metab Syndr. 2020;14(4):407–12. Epub 2020/04/27. doi: 10.1016/j.dsx.2020.04.020. PubMed PMID: 32335367; PMCID: PMC7165108.

71. Leijnse N, Barooji YF, Arastoo MR, Sonder SL, Verhagen B, Wullkopf L, Erler JT, Semsey S, Nylandsted J, Oddershede LB, Doostmohammadi A, Bendix PM. Filopodia rotate and coil by actively generating twist in their actin shaft. Nat Commun. 2022;13(1):1636. Epub 2022/03/30. doi: 10.1038/s41467-022-28961-x. PubMed PMID: 35347113; PMCID: PMC8960877.

72. Maslanka Figueroa S, Fleischmann D, Beck S, Tauber P, Witzgall R, Schweda F, Goepferich A. Nanoparticles Mimicking Viral Cell Recognition Strategies Are Superior Transporters into Mesangial Cells. Adv Sci (Weinh). 2020;7(11):1903204. Epub 2020/06/17. doi: 10.1002/advs.201903204. PubMed PMID: 32537398; PMCID: PMC7284201.

73. Lin PC, Lee YR, Liu LT, Chiou SS, Chen PC, Tsai CY, Tsai JJ. Honeysuckle extracts as a potential inhibitor of SARS-CoV-2 infection. Front Pharmacol. 2025;16:1517585. Epub 2025/05/19. doi: 10.3389/fphar.2025.1517585. PubMed PMID: 40385477; PMCID: PMC12083240.

74. Zaharuddin Z, Md Hussin NS, Karuppannan M. Real-world analysis of safety, tolerability, and adherence to nirmatrelvir-ritonavir (paxlovid) in primary care COVID-19 outpatients. Sci Rep. 2024;14(1):24750. Epub 2024/10/22. doi: 10.1038/s41598-024-75192-9. PubMed PMID: 39433826; PMCID: PMC11494129.

75. Havervall S, Marking U, Svensson J, Greilert-Norin N, Bacchus P, Nilsson P, Hober S, Gordon M, Blom K, Klingstrom J, Aberg M, Smed-Sorensen A, Thalin C. Anti-Spike Mucosal IgA Protection against SARS-CoV-2 Omicron Infection. N Engl J Med. 2022;387(14):1333–6. Epub 2022/09/15. doi: 10.1056/NEJMc2209651. PubMed PMID: 36103621; PMCID: PMC9511632.

76. Zhu J, Su, Y., and Tang, Y. Disrupting ACE2 Dimerization Mitigates the Infection by SARS-CoV-2 Pseudovirus. Frontiers in Virology. 2022;Virol., 03 July 2022(2).

77. Lorenzo R, Defelipe LA, Aliperti L, Niebling S, Custodio TF, Low C, Schwarz JJ, Remans K, Craig PO, Otero LH, Klinke S, Garcia-Alai M, Sanchez IE, Alonso LG. Deamidation drives molecular aging of the SARS-CoV-2 spike protein receptor-binding motif. J Biol Chem. 2021;297(4):101175. Epub 2021/09/10. doi: 10.1016/j.jbc.2021.101175. PubMed PMID: 34499924; PMCID: PMC8421091.

78. Beaudoin CA, Petsolari E, Hamaia SW, Hala S, Alofi FS, Pandurangan AP, Blundell TL, Chaitanya Vedithi S, Huang CL, Jackson AP. SARS-CoV-2 Omicron subvariant spike N405 unlikely to rapidly deamidate. Biochem Biophys Res Commun. 2023;666:61–7. Epub 2023/05/14. doi: 10.1016/j.bbrc.2023.04.088. PubMed PMID: 37178506; PMCID: PMC10152834.

79. Yi C, Sun X, Ye J, Ding L, Liu M, Yang Z, Lu X, Zhang Y, Ma L, Gu W, Qu A, Xu J, Shi Z, Ling Z, Sun B. Key residues of the receptor binding motif in the spike protein of SARS-CoV-2 that interact with ACE2 and neutralizing antibodies. Cell Mol Immunol. 2020;17(6):621–30. Epub 2020/05/18. doi: 10.1038/s41423-020-0458-z. PubMed PMID: 32415260; PMCID: PMC7227451.

80. Chopra A, Shukri AH, Adhikary H, Lukinovic V, Hoekstra M, Cowpland M, Biggar KK. A peptide array pipeline for the development of Spike-ACE2 interaction inhibitors. Peptides. 2022;158:170898. Epub 2022/10/25. doi: 10.1016/j.peptides.2022.170898. PubMed PMID: 36279985; PMCID: PMC9585897.

81. Cao L, Goreshnik I, Coventry B, Case JB, Miller L, Kozodoy L, Chen RE, Carter L, Walls AC, Park YJ, Strauch EM, Stewart L, Diamond MS, Veesler D, Baker D. De novo design of picomolar SARS-CoV-2 miniprotein inhibitors. Science. 2020;370(6515):426–31. Epub 2020/09/11. doi: 10.1126/science.abd9909. PubMed PMID: 32907861; PMCID: PMC7857403.

82. Beaudoin CA, Jamasb AR, Alsulami AF, Copoiu L, van Tonder AJ, Hala S, Bannerman BP, Thomas SE, Vedithi SC, Torres PHM, Blundell TL. Predicted structural mimicry of spike receptor-binding motifs from highly pathogenic human coronaviruses. Comput Struct Biotechnol J. 2021;19:3938–53. Epub 2021/07/09. doi: 10.1016/j.csbj.2021.06.041. PubMed PMID: 34234921; PMCID: PMC8249111.

83. Eslami N, Aghbash PS, Shamekh A, Entezari-Maleki T, Nahand JS, Sales AJ, Baghi HB. SARS-CoV-2: Receptor and Co-receptor Tropism Probability. Curr Microbiol. 2022;79(5):133. Epub 2022/03/17. doi: 10.1007/s00284-022-02807-7. PubMed PMID: 35292865; PMCID: PMC8923825.

84. Alipoor SD, Mirsaeidi M. SARS-CoV-2 cell entry beyond the ACE2 receptor. Mol Biol Rep. 2022;49(11):10715–27. Epub 2022/06/27. doi: 10.1007/s11033-022-07700-x. PubMed PMID: 35754059; PMCID: PMC9244107.

85. Beddingfield BJ, Iwanaga N, Chapagain PP, Zheng W, Roy CJ, Hu TY, Kolls JK, Bix GJ. The Integrin Binding Peptide, ATN-161, as a Novel Therapy for SARS-CoV-2 Infection. JACC Basic Transl Sci. 2021;6(1):1-8. Epub 2020/10/27. doi: 10.1016/j.jacbts.2020.10.003. PubMed PMID: 33102950; PMCID: PMC7566794.

86. Beaudoin CA, Hamaia SW, Huang CL, Blundell TL, Jackson AP. Can the SARS-CoV-2 Spike Protein Bind Integrins Independent of the RGD Sequence? Front Cell Infect Microbiol. 2021;11:765300. Epub 2021/12/07. doi: 10.3389/fcimb.2021.765300. PubMed PMID: 34869067; PMCID: PMC8637727.

87. Kliche J, Kuss H, Ali M, Ivarsson Y. Cytoplasmic short linear motifs in ACE2 and integrin beta(3) link SARS-CoV-2 host cell receptors to mediators of endocytosis and autophagy. Sci Signal. 2021;14(665). Epub 2021/01/14. doi: 10.1126/scisignal.abf1117. PubMed PMID: 33436498; PMCID: PMC7928716.

88. Zamorano Cuervo N, Grandvaux N. ACE2: Evidence of role as entry receptor for SARS-CoV-2 and implications in comorbidities. Elife. 2020;9. Epub 2020/11/10. doi: 10.7554/eLife.61390. PubMed PMID: 33164751; PMCID: PMC7652413.

89. Sui Y, Li J, Venzon DJ, Berzofsky JA. SARS-CoV-2 Spike Protein Suppresses ACE2 and Type I Interferon Expression in Primary Cells From Macaque Lung Bronchoalveolar Lavage. Front Immunol. 2021;12:658428. Epub 2021/06/22. doi: 10.3389/fimmu.2021.658428. PubMed PMID: 34149696; PMCID: PMC8213020.

90. Ugo Bastolla1* PC, David Abia1, Maria-Laura Garcia-Bermejo3 and Manuel Fresno1. Is Covid-19 Severity Associated With ACE2 Degradation? Frontiers in Drug Discovery. 2022(January 2022 | Volume 1 | Article 789710). Epub 10.3389/fddsv.2021.789710.

91. Trougakos IP, Terpos E, Alexopoulos H, Politou M, Paraskevis D, Scorilas A, Kastritis E, Andreakos E, Dimopoulos MA. Adverse effects of COVID-19 mRNA vaccines: the spike hypothesis. Trends Mol Med. 2022;28(7):542–54. Epub 2022/05/11. doi: 10.1016/j.molmed.2022.04.007. PubMed PMID: 35537987; PMCID: PMC9021367.

92. Angeli F, Spanevello A, Reboldi G, Visca D, Verdecchia P. SARS-CoV-2 vaccines: Lights and shadows. Eur J Intern Med. 2021;88:1–8. Epub 2021/05/11. doi: 10.1016/j.ejim.2021.04.019. PubMed PMID: 33966930; PMCID: PMC8084611.

93. Zappa M, Verdecchia P, Spanevello A, Visca D, Angeli F. Blood pressure increase after Pfizer/BioNTech SARS-CoV-2 vaccine. Eur J Intern Med. 2021;90:111–3. Epub 2021/06/24. doi: 10.1016/j.ejim.2021.06.013. PubMed PMID: 34158234; PMCID: PMC8206586.

94. Forte E. Circulating spike protein may contribute to myocarditis after COVID-19 vaccination. Nat Cardiovasc Res. 2023;2(2):100. Epub 2023/02/01. doi: 10.1038/s44161-023-00222-0. PubMed PMID: 39196056.

95. Walsh EE, Frenck RW, Jr., Falsey AR, Kitchin N, Absalon J, Gurtman A, Lockhart S, Neuzil K, Mulligan MJ, Bailey R, Swanson KA, Li P, Koury K, Kalina W, Cooper D, Fontes-Garfias C, Shi PY, Tureci O, Tompkins KR, Lyke KE, Raabe V, Dormitzer PR, Jansen KU, Sahin U, Gruber WC. Safety and Immunogenicity of Two RNA-Based Covid-19 Vaccine Candidates. N Engl J Med. 2020;383(25):2439–50. Epub 2020/10/15. doi: 10.1056/NEJMoa2027906. PubMed PMID: 33053279; PMCID: PMC7583697.

96. Haas JW, Bender FL, Ballou S, Kelley JM, Wilhelm M, Miller FG, Rief W, Kaptchuk TJ. Frequency of Adverse Events in the Placebo Arms of COVID-19 Vaccine Trials: A Systematic Review and Meta-analysis. JAMA Netw Open. 2022;5(1):e2143955. Epub 2022/01/19. doi: 10.1001/jamanetworkopen.2021.43955. PubMed PMID: 35040967; PMCID: PMC8767431 scholarship from the German Academic Exchange Service (Deutscher Akademischer Austauschdienst) during the conduct of the study. No other disclosures were reported.

97. Illing N, Boolay S, Siwoski JS, Casper D, Lucero MT, Roskams AJ. Conditionally immortalized clonal cell lines from the mouse olfactory placode differentiate into olfactory receptor neurons. Mol Cell Neurosci. 2002;20(2):225–43. Epub 2002/07/03. doi: 10.1006/mcne.2002.1106. PubMed PMID: 12093156.

98. Ma G, Syu GD, Shan X, Henson B, Wang S, Desai PJ, Zhu H, Tao N. Measuring Ligand Binding Kinetics to Membrane Proteins Using Virion Nano-oscillators. J Am Chem Soc. 2018;140(36):11495–501. Epub 2018/08/17. doi: 10.1021/jacs.8b07461. PubMed PMID: 30114365; PMCID: PMC6211285.

99. Hati S, Bhattacharyya S. Impact of Thiol-Disulfide Balance on the Binding of Covid-19 Spike Protein with Angiotensin-Converting Enzyme 2 Receptor. ACS Omega. 2020;5(26):16292–8. Epub 2020/07/14. doi: 10.1021/acsomega.0c02125. PubMed PMID: 32656452; PMCID: PMC7346263.

100. Argentinian AntiCovid C. Structural and functional comparison of SARS-CoV-2-spike receptor binding domain produced in Pichia pastoris and mammalian cells. Sci Rep. 2020;10(1):21779. Epub 2020/12/15. doi: 10.1038/s41598-020-78711-6. PubMed PMID: 33311634; PMCID: PMC7732851.

