## Supplementary figures and images for "Visualizing the Rapid Degradation and Differential Binding of Membrane-Bound Human ACE2 Protein Upon Binding and Internalization of the SARS-CoV-2 Spike RBD Protein"

### Supplemental Figure S1

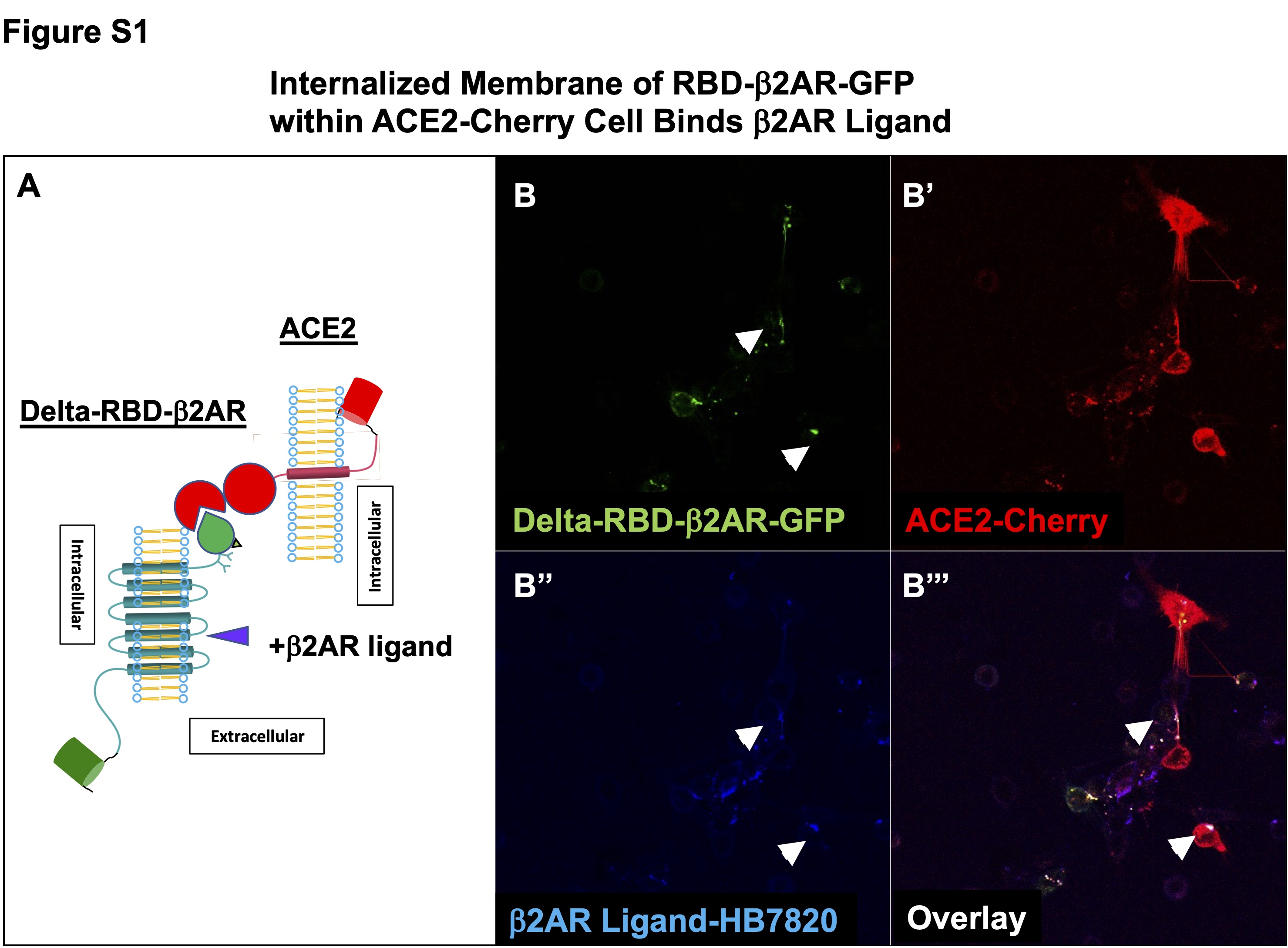

### Supplemental Figure S2

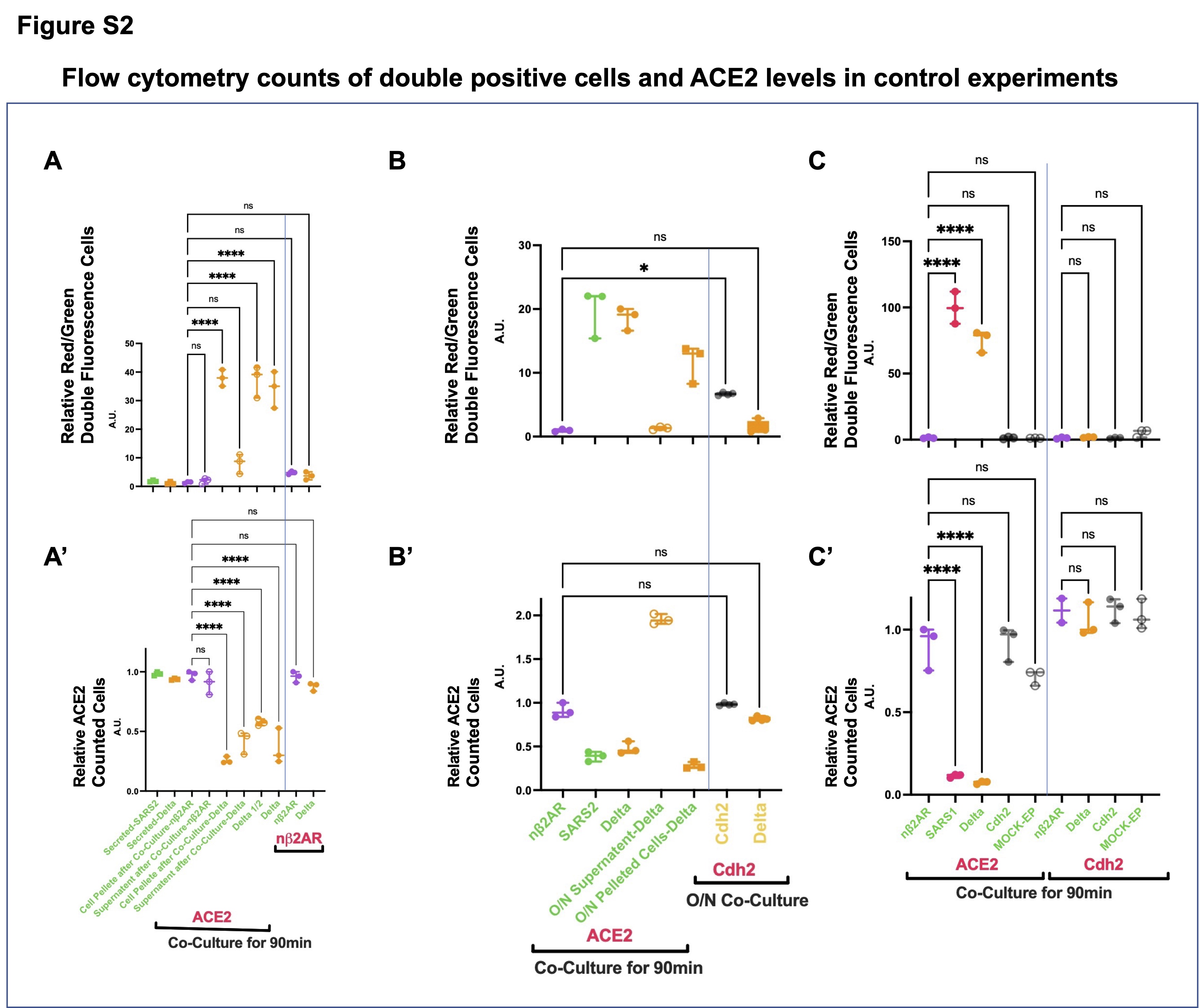

### Supplemental Figure S3

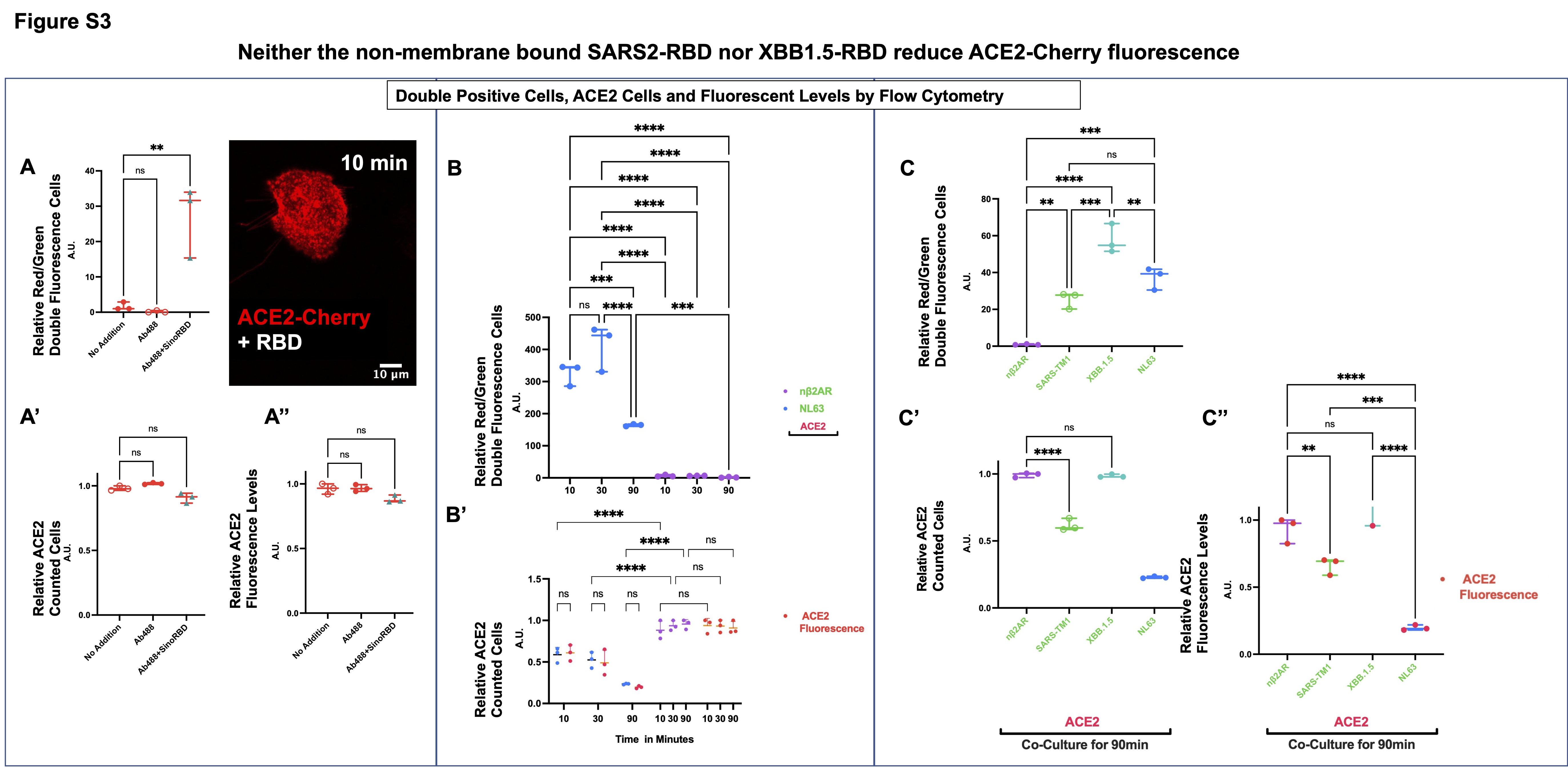

### Supplemental Figure S4

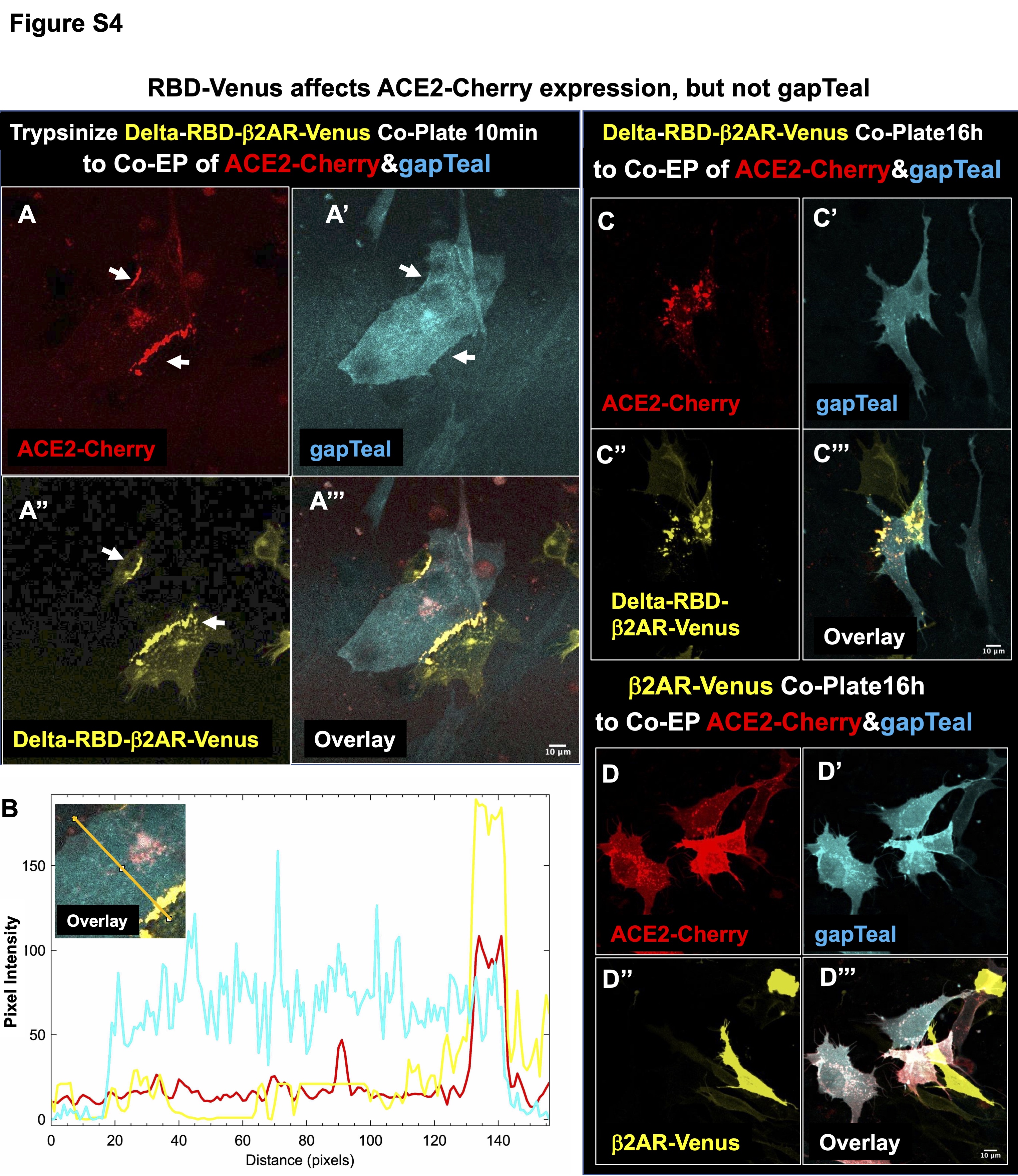

### Supplemental Figure S5

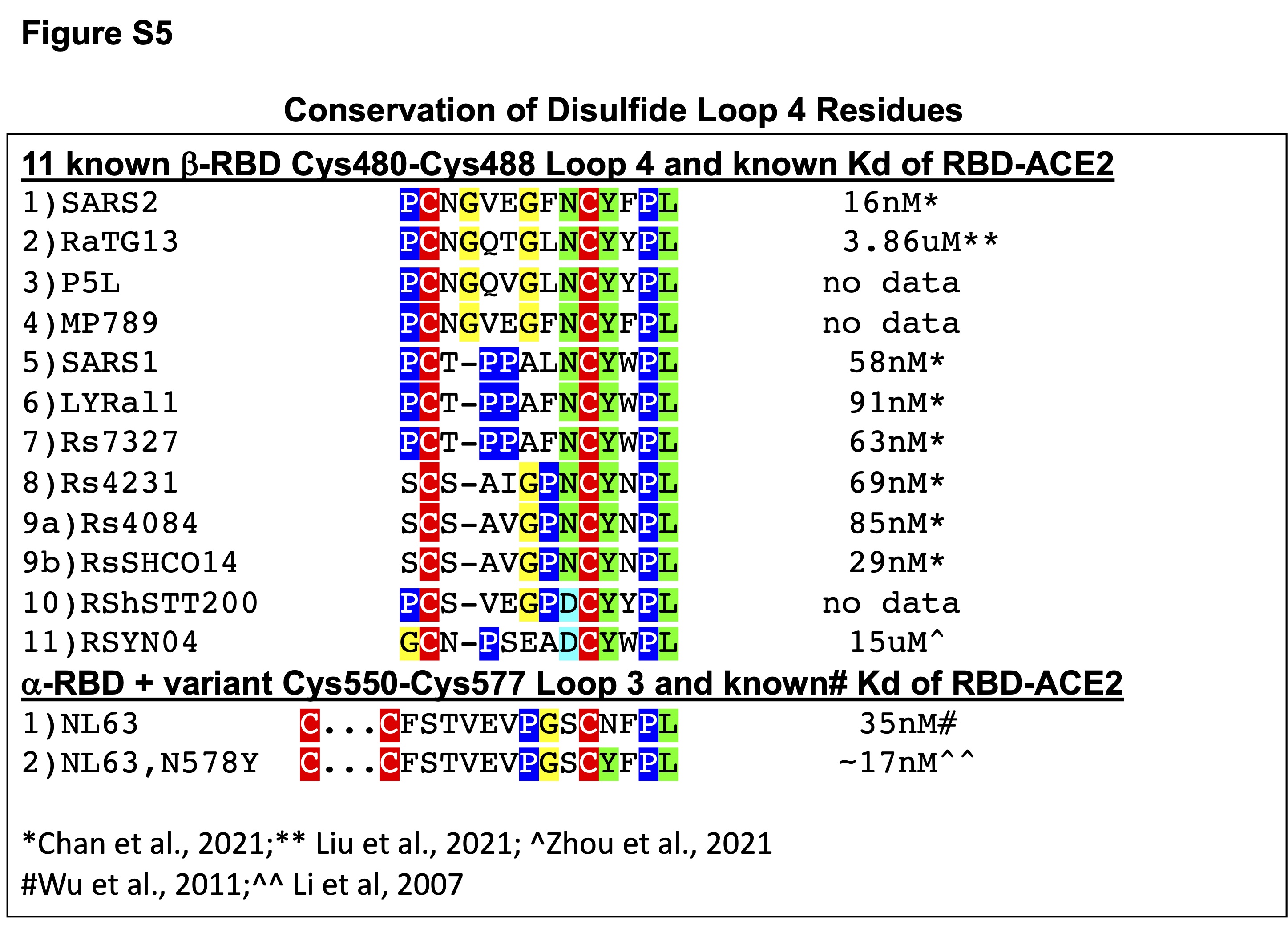

### Supplemental Figure S6

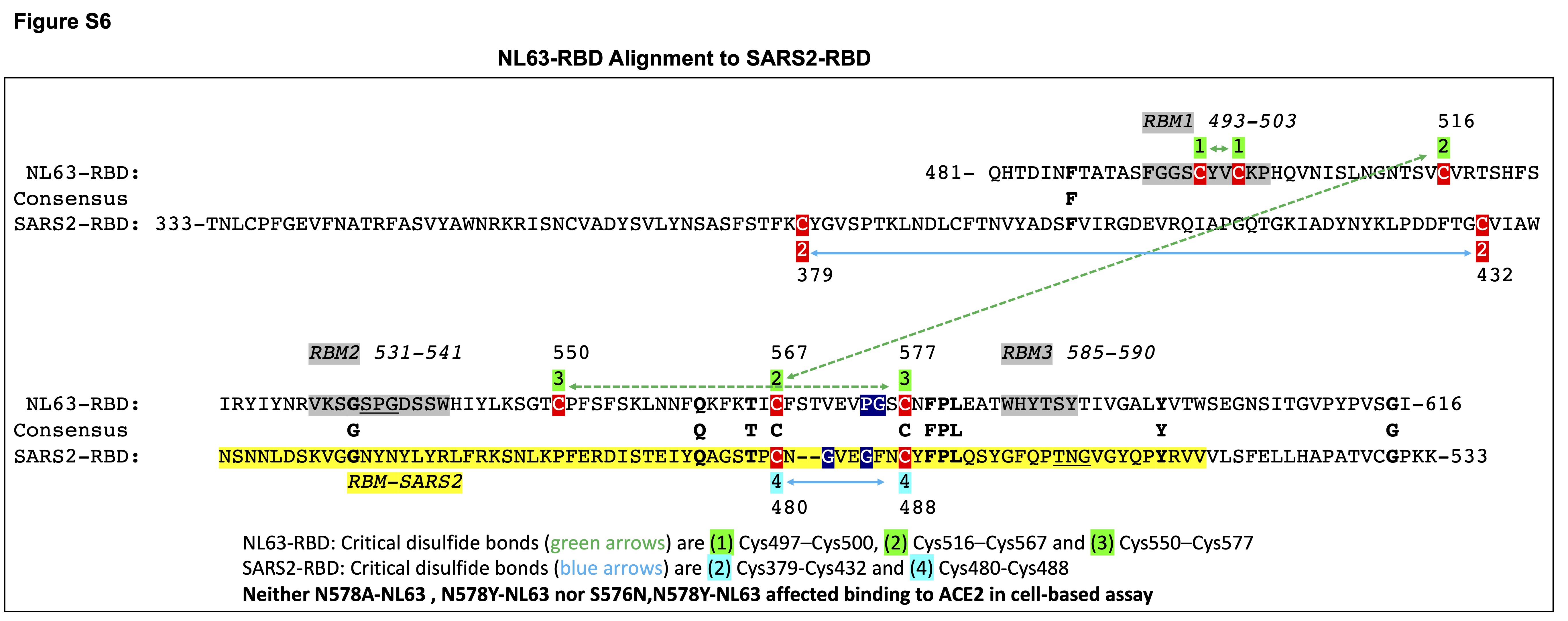

### Supplemental Figure S7

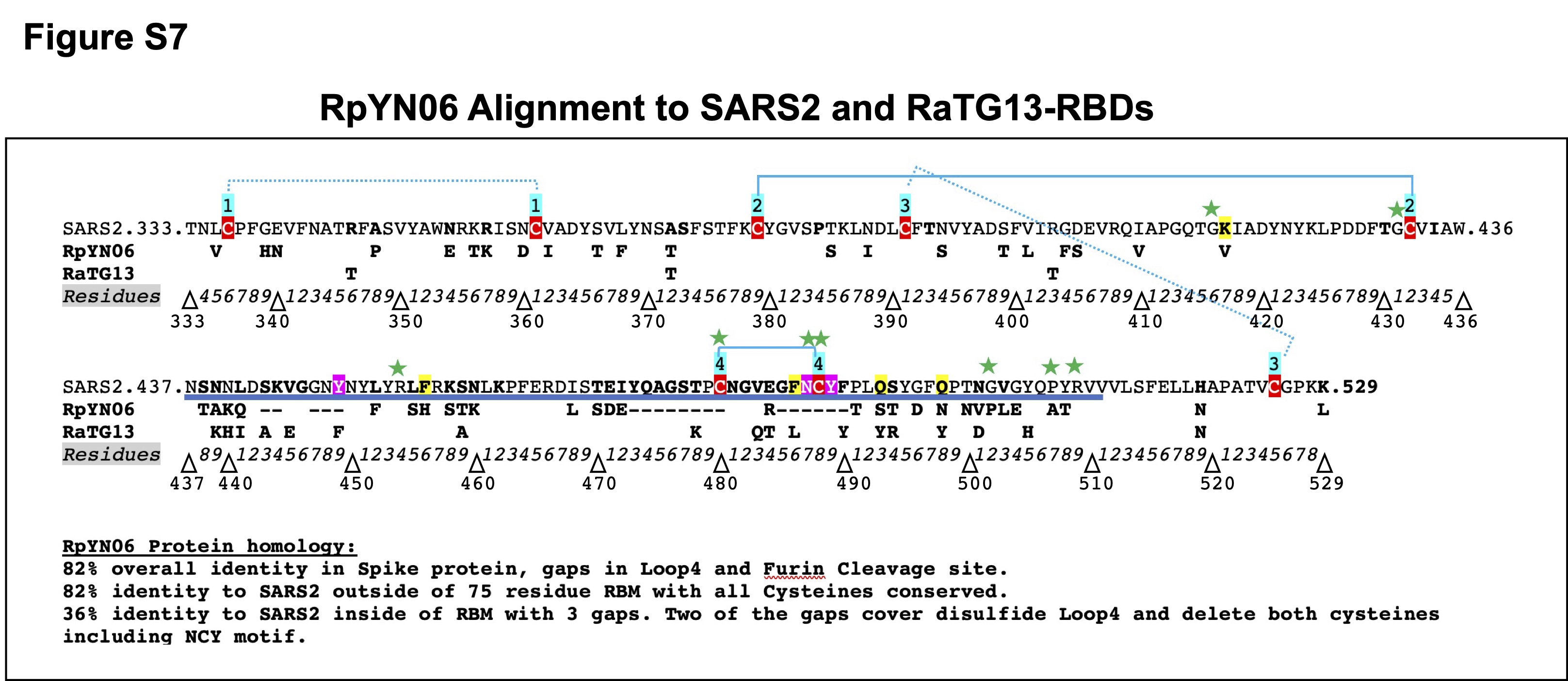

### Supplemental Figure S8

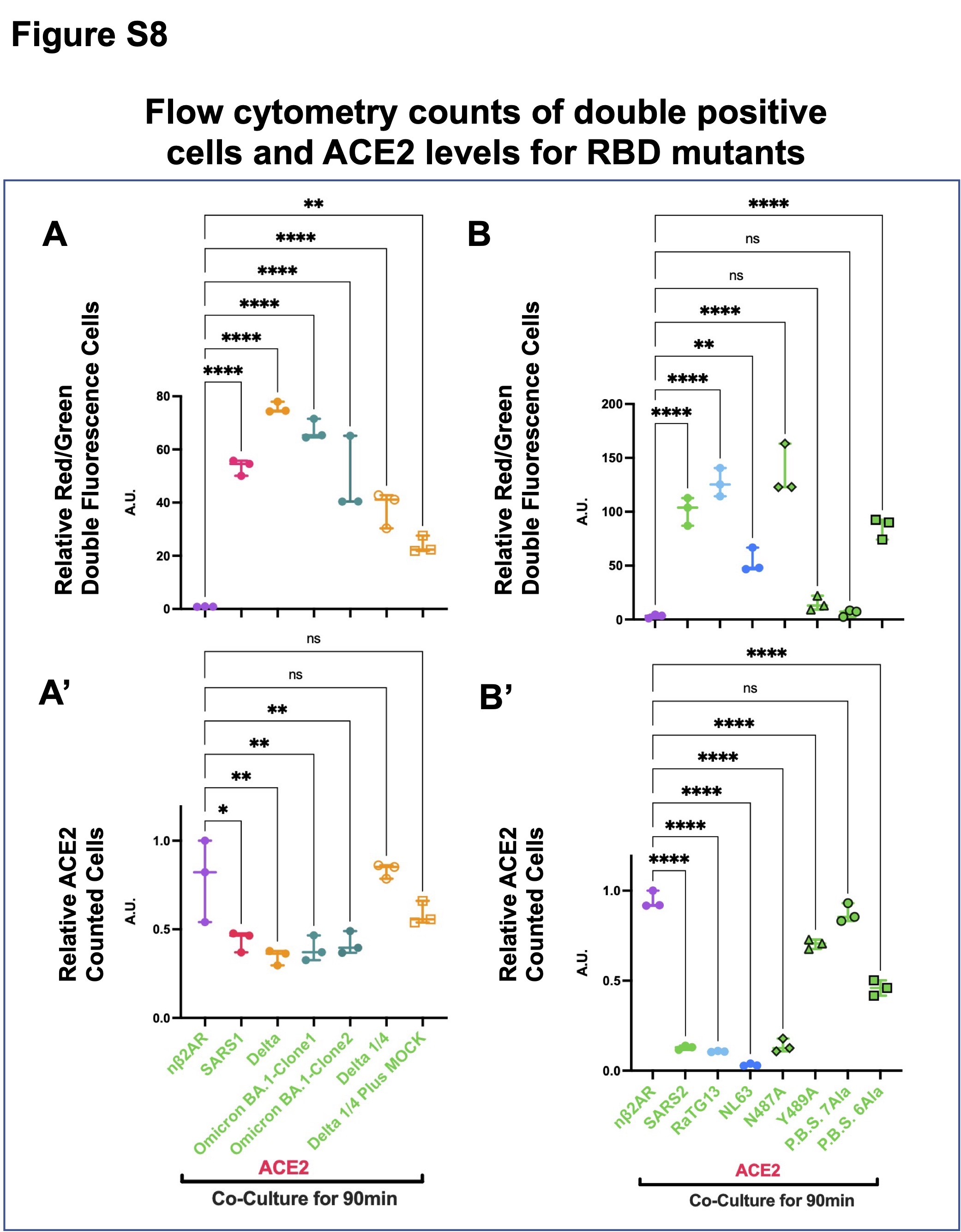

### Supplemental Figure S9

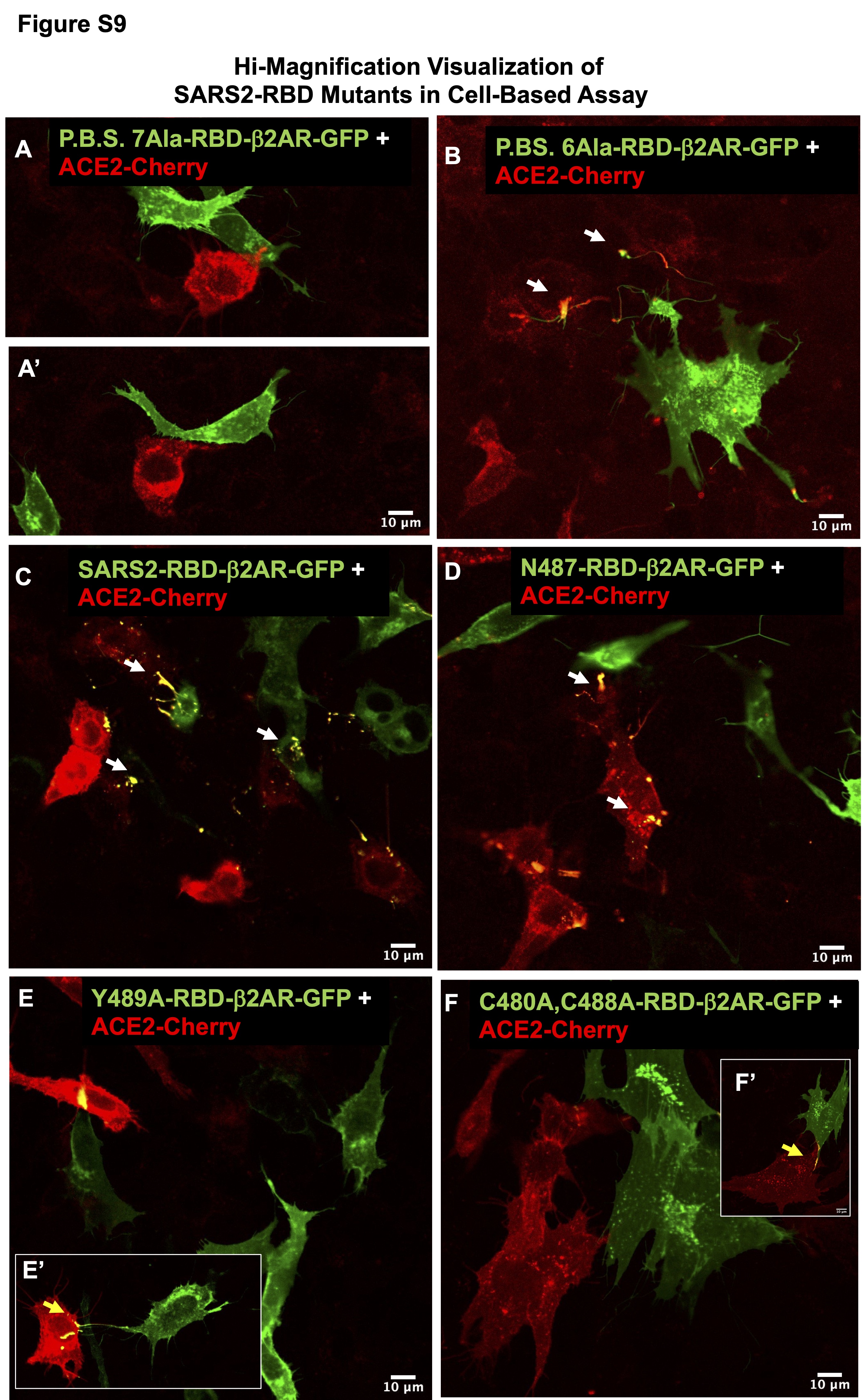

### Supplemental Figure S10

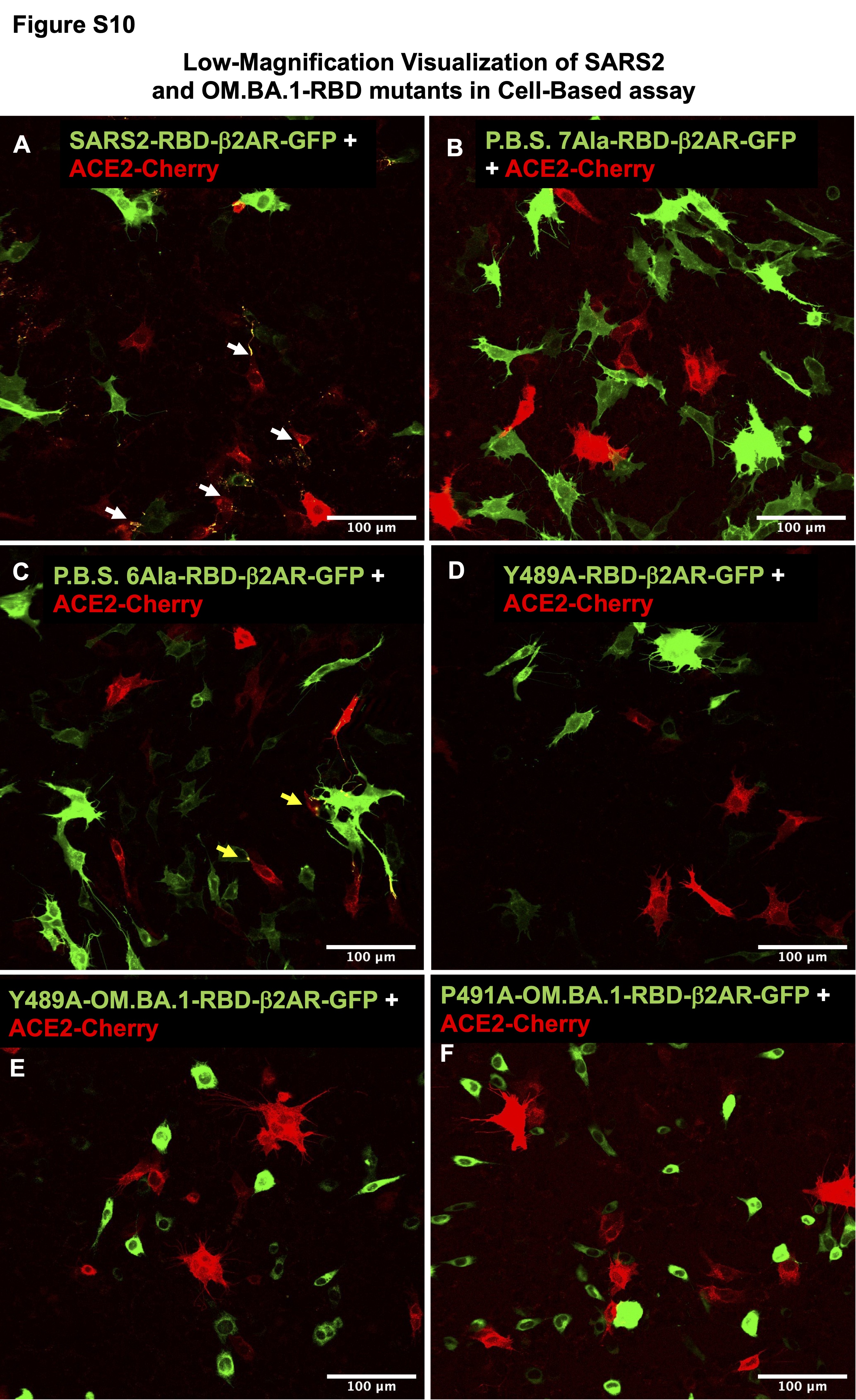

### Supplemental Figure S11

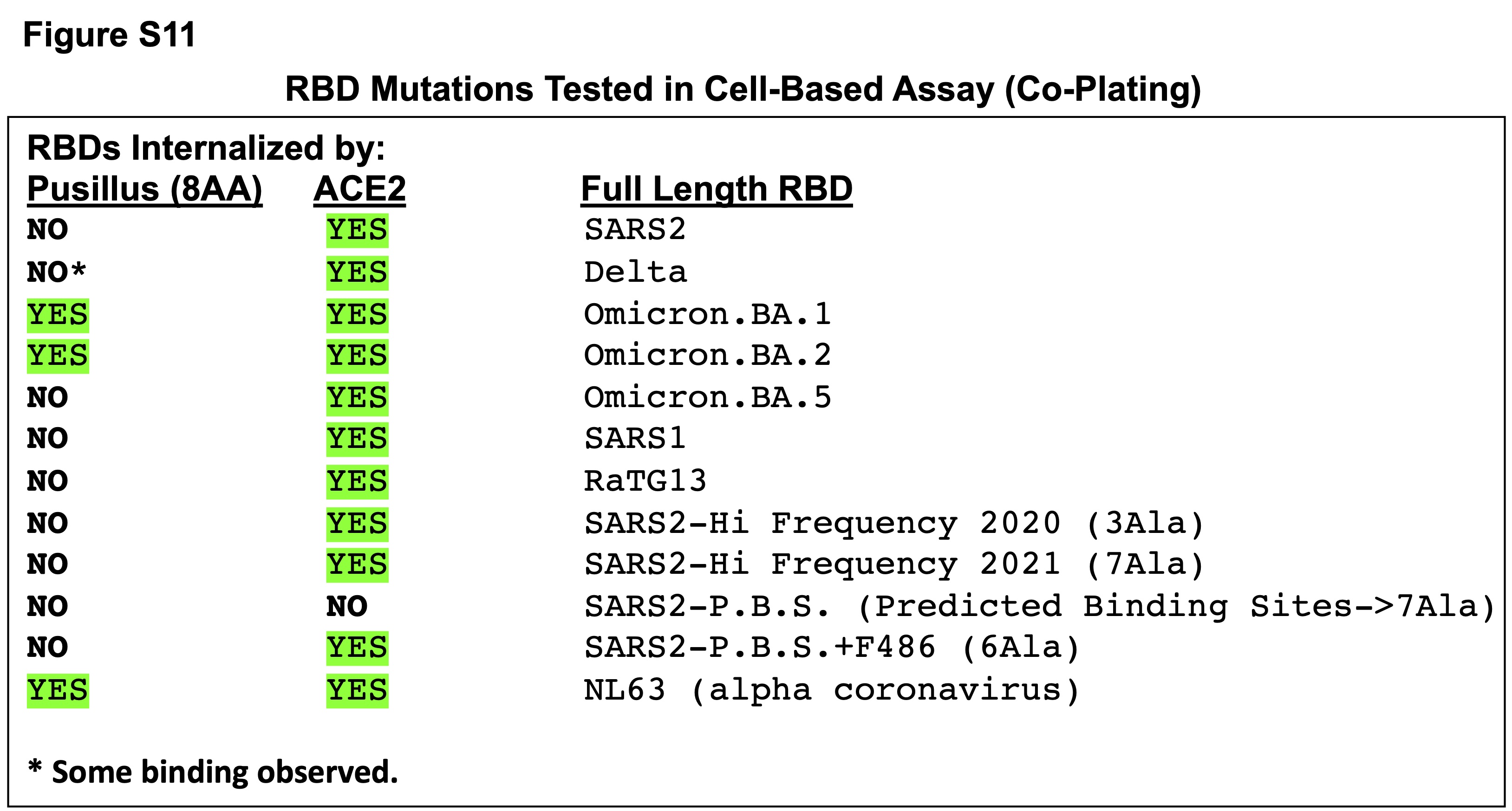

### Supplemental Figure S12

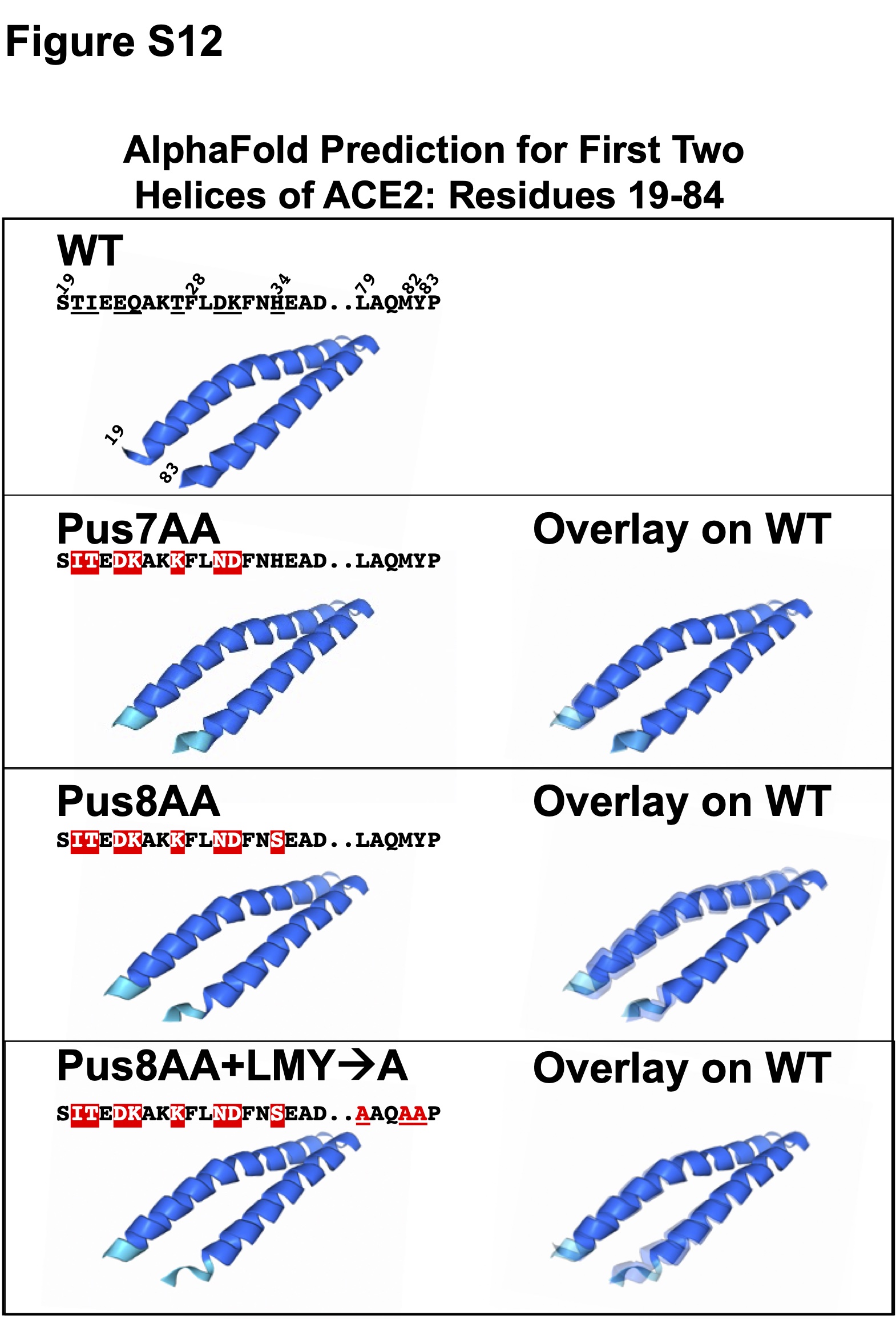

### Supplemental Figure S13

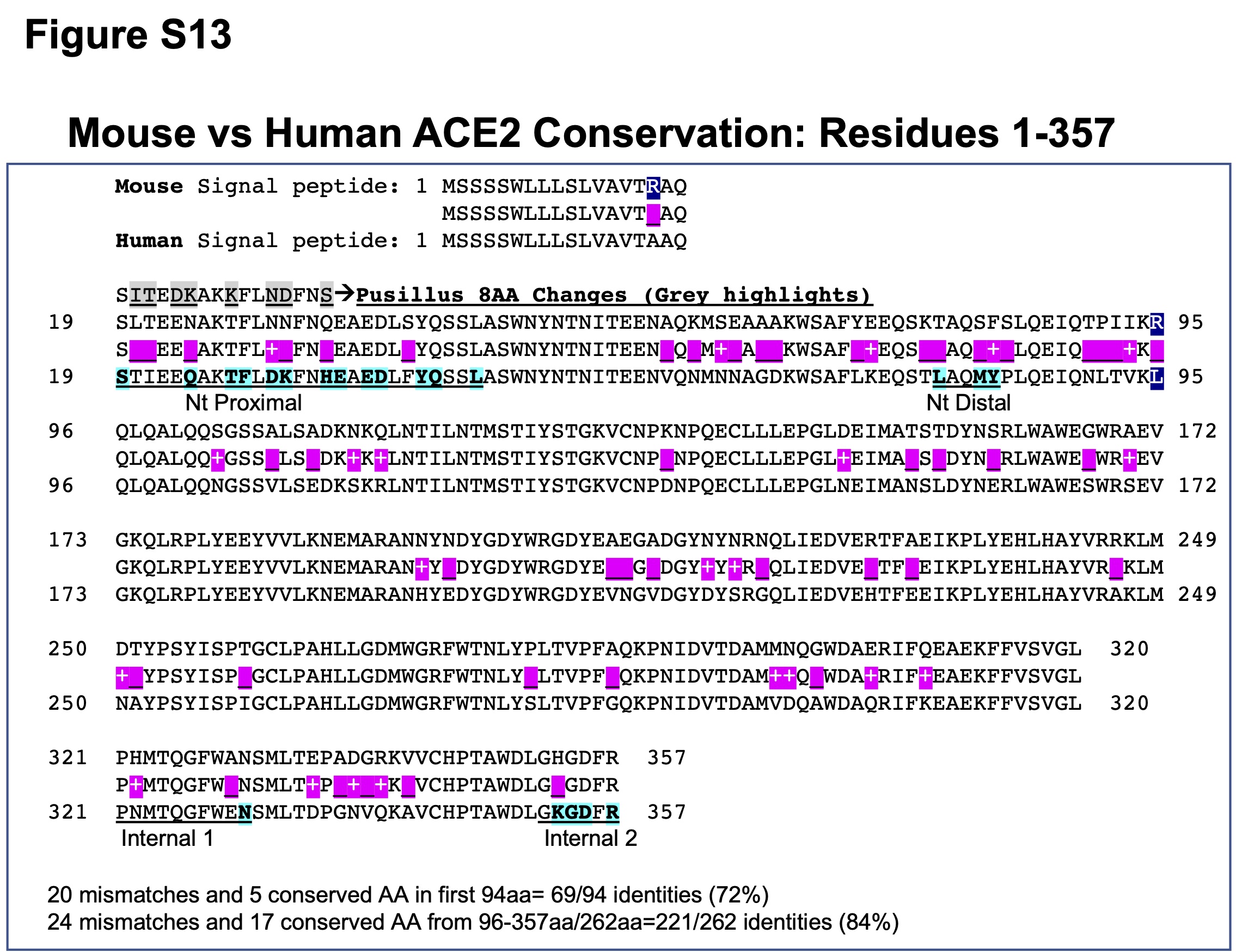

### Supplemental Figure S14

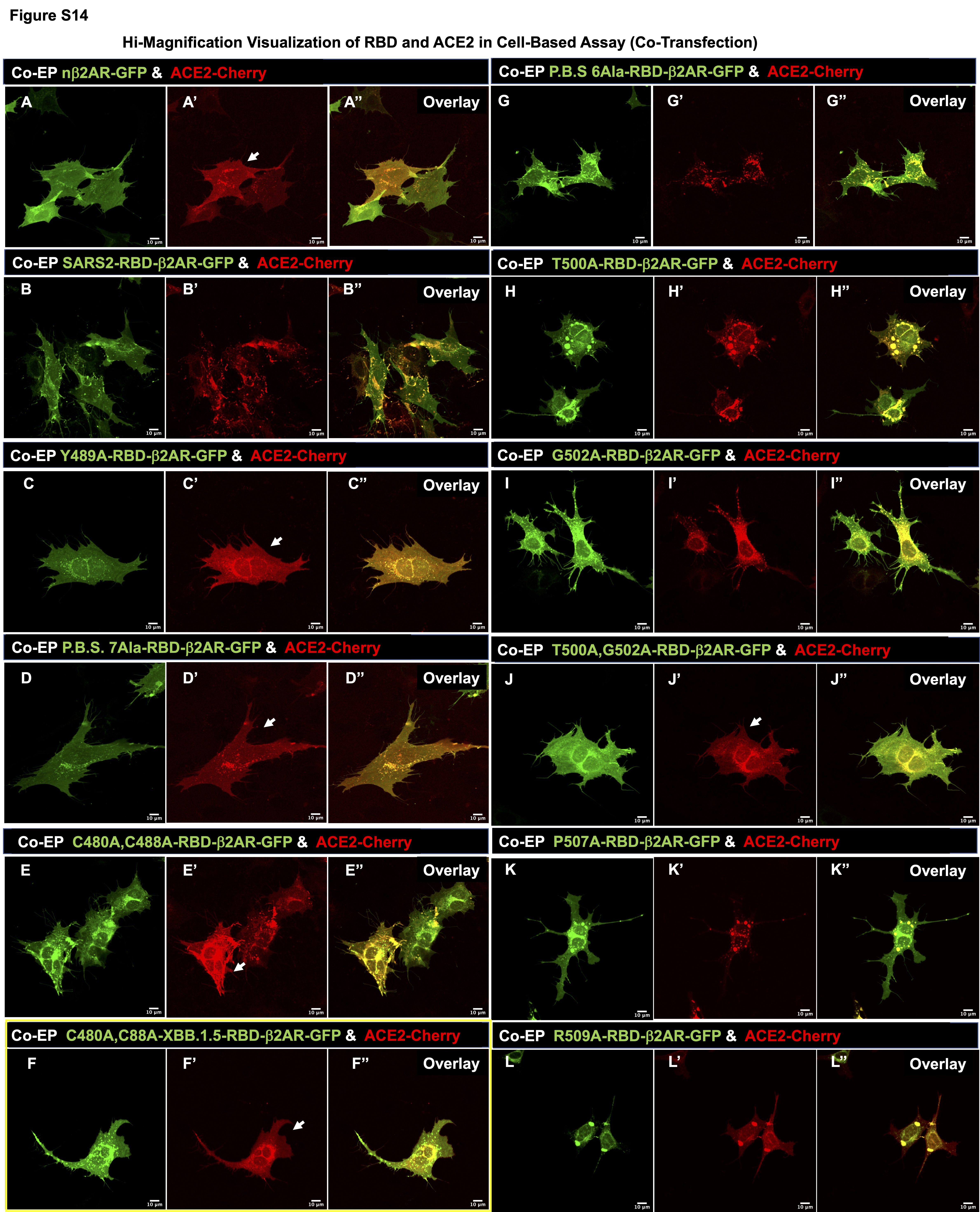

### Supplemental Figure S15

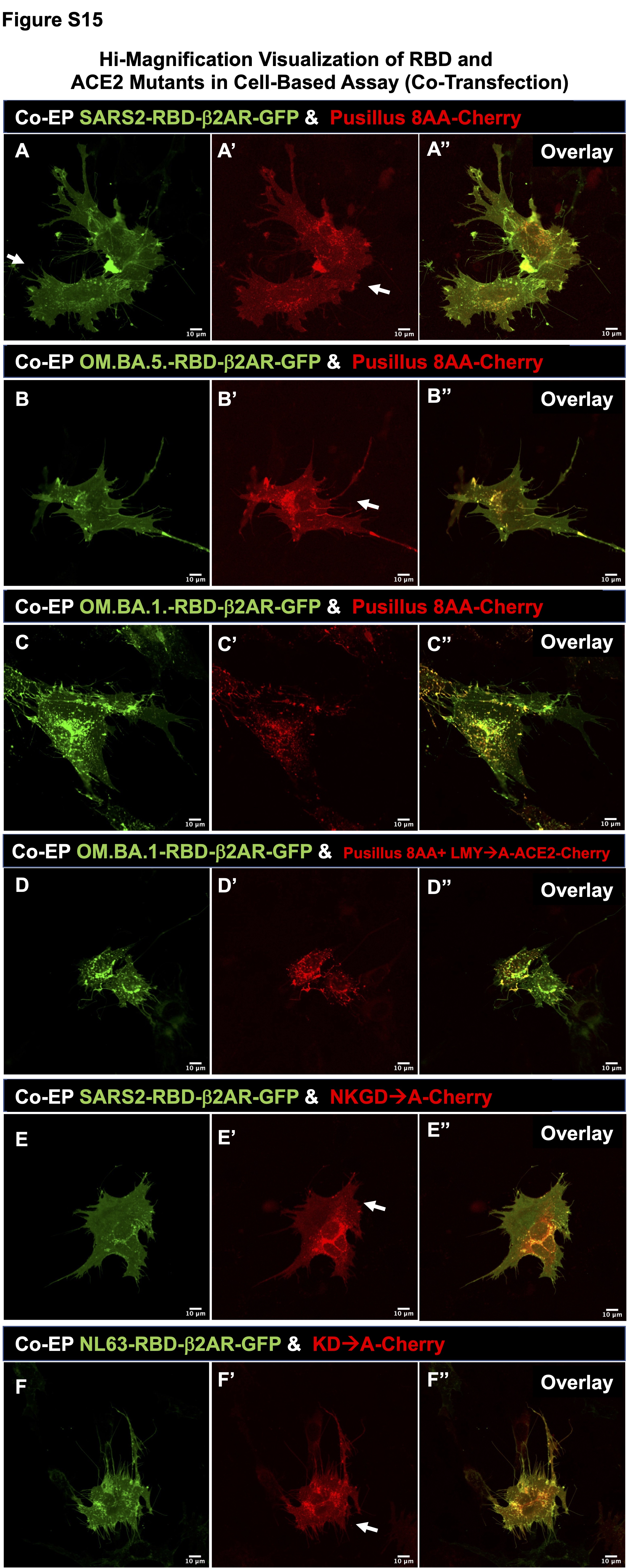

### Supplemental Figure S16

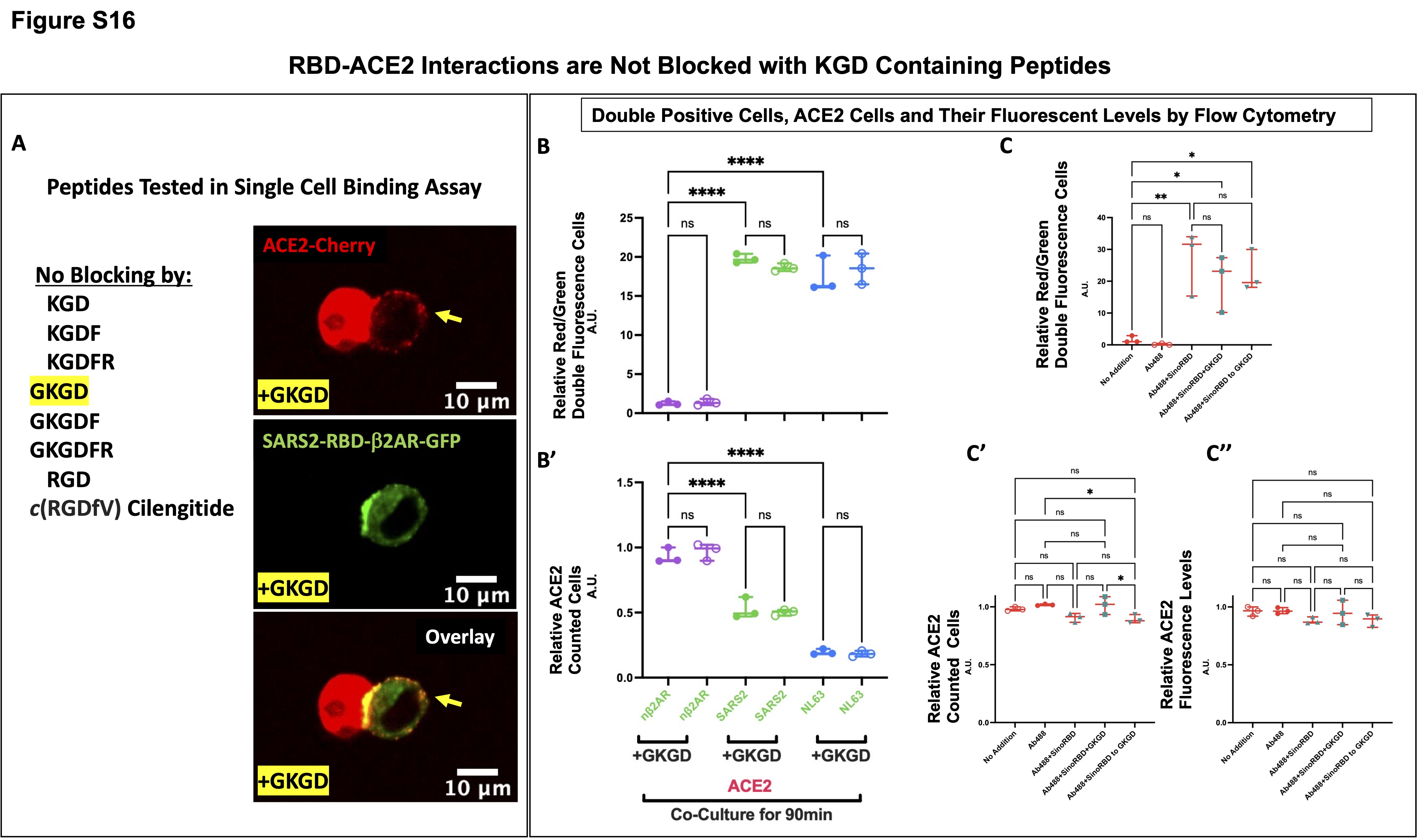

### Supplemental Figure S17

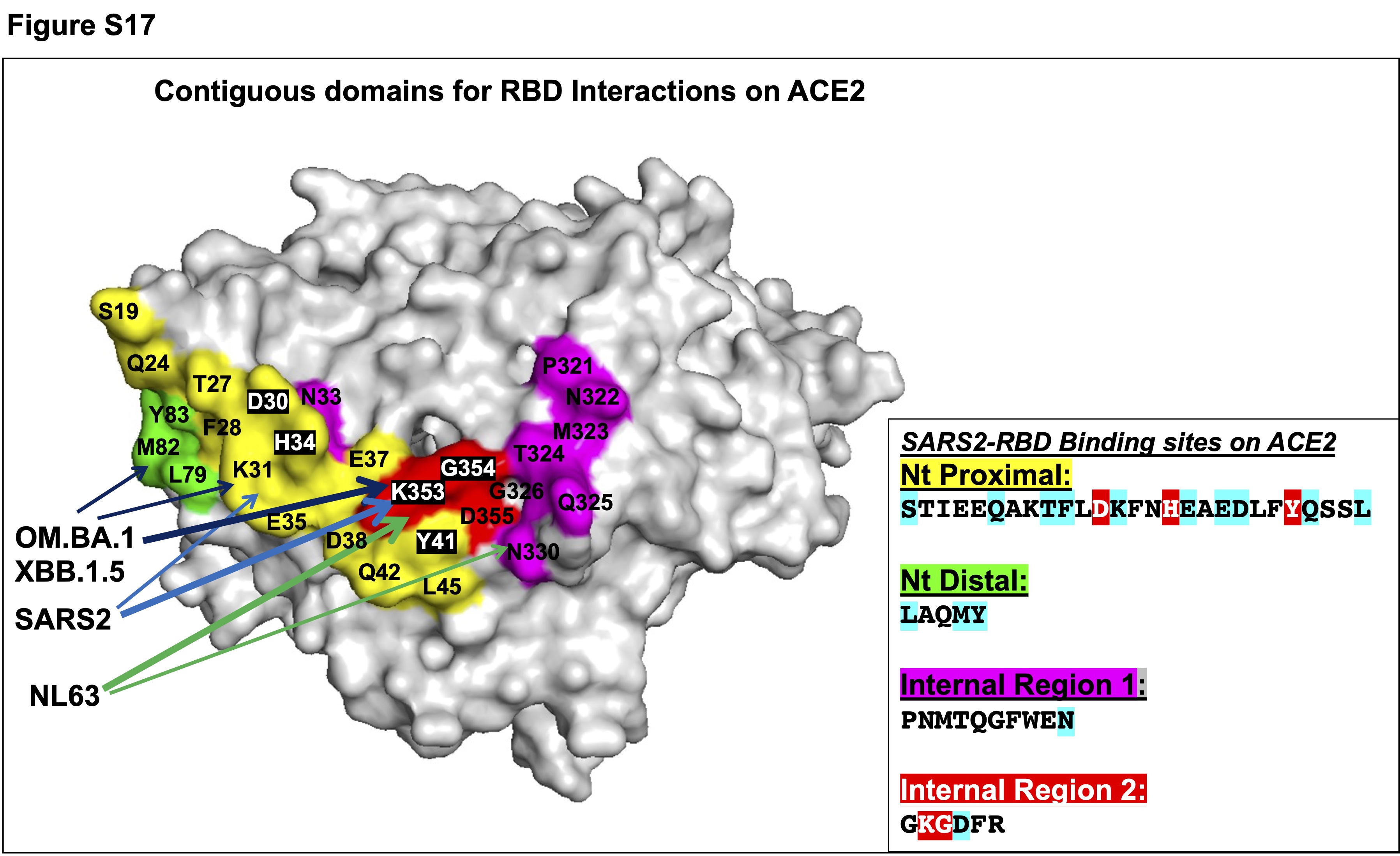
